# Highly Pathogenic Avian Influenza Viruses H5N1 and H5N5 in Red Foxes (*Vulpes vulpes*) in Norway during 2022-2024

**DOI:** 10.64898/2026.09.25.754393

**Authors:** Malin Rokseth Reiten, Cathrine A. Bøe, Line Olsen, Maryam Saghafian, Andreas Rohringer, Johan Åkerstedt, Bjørnar Ytrehus, Britt Gjerset, Silje Granstad, Olav Hungnes, Ragnhild Tønnessen

## Abstract

Since 2020, highly pathogenic avian influenza (HPAI) A(H5Nx) clade 2.3.4.4b viruses have spread globally, causing extensive outbreaks in domestic and wild birds. Increased circulation has resulted in frequent spillover to mammals, and occasional mammal-to-mammal transmission. Although human infections remain rare, the zoonotic potential of these viruses continues to be a public health concern. We investigated six cases of HPAI in red foxes (*Vulpes vulpes*) in Norway during 2022-2024 and identified infections with H5N1 and H5N5 viruses. Pathological and virological investigations demonstrated systemic infection, with prominent lesions in the brain and/or lungs. Phylogenetic analyses showed high similarity between viruses detected in foxes and those concurrently circulating in wild birds. The mammalian adaptation marker PB2:E627K was identified in one H5N1 virus and as a minority variant in a second H5N1 virus.

## Introduction

Highly pathogenic avian influenza (HPAI) A(H5Nx) clade 2.3.4.4b viruses, derived from the A/Goose/Guangdong/1/1996 lineage, emerged in Europe in 2016 (1). In 2021, HPAI H5N1 became established in wild bird populations across Europe (2) and subsequently spread globally via migratory birds (3, 4). Concurrent with the extensive circulation in birds, clade 2.3.4.4b HPAI viruses (HPAIVs) have increasingly been detected in wild and domestic mammals worldwide, with infections reported in more than 80 species (3, 5).

Although most mammalian infections are considered independent spillover events from birds, mammal-to-mammal transmission has been implicated in outbreaks among farmed mink in Spain, fur animals in Finland, and marine mammals in South America (6–9). More recently, widespread transmission among dairy cattle in the United States with spillover to cats and humans has further raised concerns regarding the pandemic potential of these viruses (10–12).

Red foxes (*Vulpes vulpes*) are widely distributed carnivores in the Northern Hemisphere and are opportunistic predators and scavengers. Frequent contact with sick and dead wild birds likely increases their exposure to HPAIVs during outbreaks. Consequently, red foxes have emerged as an important sentinel species for monitoring HPAI circulation and are among the wild mammalian species most frequently reported with HPAIV infection (5). Reported infections have been associated with respiratory and neurological disease, systemic viral dissemination, and detection of mammalian-adaptive mutations, including PB2:E627K (13–16). Recent serological evidence suggests that avian influenza virus exposure in red foxes may be more widespread than indicated by passive surveillance alone (17, 18).

In Northern Europe, HPAIV H5N1 predominated in wild birds throughout 2022-2024, with EA-2020-C, EA-2021-AB, and EA-2022-BB constituting the major genotypes (1). The H5N1 subtype first emerged in Norway in 2021 in wild birds and poultry (19, 20).

The HPAIV H5N5 subtype was first reported in wild birds in Europe in 2016 (21) and was subsequently identified in Norway in 2022 in a white-tailed eagle (*Haliaeetus albicilla*) (20, 22). During 2023 and 2024, H5N5 genotype EA-2021-I also spread to wild birds in other parts of Northern Europe, Canada (23, 24) and Japan (25). Although less frequently detected than H5N1, H5N5 circulation may be underrepresented by routine surveillance activities. Recent studies from the Arctic have reported only sporadic H5N5 detections, yet widespread seroreactivity in polar bears (*Ursus maritimus*) and high H5N5 seroprevalence in Arctic foxes (*Vulpes lagopus*) on Svalbard suggest that H5N5 circulation may be more extensive in this ecosystem than virus detection data alone indicate (26–28). The zoonotic potential of H5N5 was further illustrated by the first reported human infection in the United States in 2025 (29). However, despite increasing reports of infection in wildlife and humans, H5N5 has been far less studied in mammals than H5N1.

Between 2022 and 2024, HPAIV infection was confirmed in wild red foxes in Norway. We investigated these cases to characterize the pathological, virological and genomic features of HPAI infection in red foxes and to determine whether fox-derived viruses were consistent with spillover from contemporaneously circulating avian viruses and showed evidence of mammalian adaptation. By documenting spillover of both H5N1 and H5N5 viruses into red foxes, this study provides insights into HPAI ecology, mammalian adaptation, and potential animal and public health risks.

## Materials and Methods

Detailed descriptions of sample collection, diagnostic testing, pathological investigations, whole-genome sequencing (WGS), phylogenetic analyses, and mutation screening are provided in the Appendix. Virus sequences and associated metadata included in the phylogenetic analyses are listed in Supplementary File 1 and 2 and can also be found at https://doi.org/10.55876/gis8.260923pq.

Between 2022 and 2024, six red foxes sampled through wildlife surveillance in mainland Norway were tested at the Norwegian Veterinary Institute by influenza A matrix (M) gene real-time reverse transcription PCR (rRT-PCR) and confirmed positive for HPAIV. Results from passive and active surveillance data from wild birds were used to provide epidemiological context, and spatial analyses were performed using R-based tools.

Pathological investigations included necropsy, histopathology, immunohistochemistry (IHC), and RNAscope *in situ* hybridization to assess viral distribution and tissue tropism. Quantification cycle (Cq) values from the semi-quantitative rRT-PCR targeting the influenza A virus M gene were used as a proxy for viral load.

To investigate the relationships between fox-derived viruses and viruses circulating in wild birds, WGS, assembly, genotyping, phylogenetic analysis and screening for mammalian-adaptive mutations were performed. Viral genomes were evaluated for mutations associated with mammalian adaptation according to published criteria (30).

## Results

### HPAIV Detections in Red Foxes and Wild Birds During 2022-2024

Between 2022 and 2024, 18 red foxes were submitted and tested for HPAIV, of which six were found positive (Figure 1). Four foxes were infected with H5N1: Fox Selje (2022), Fox Refvik (2022), Fox Averøy (2022), and Fox Tromsø (2023); two were infected with H5N5: Fox Skibotn 1 (2024) and Fox Skibotn 2 (2024). All viruses belonged to H5 HA clade 2.3.4.4b. Five foxes had been observed with neurological signs before death or euthanasia, including lethargy, reduced shyness, circling, lameness and inability to walk. The sixth fox was found dead (Appendix Table 1).

**Figure 1.**
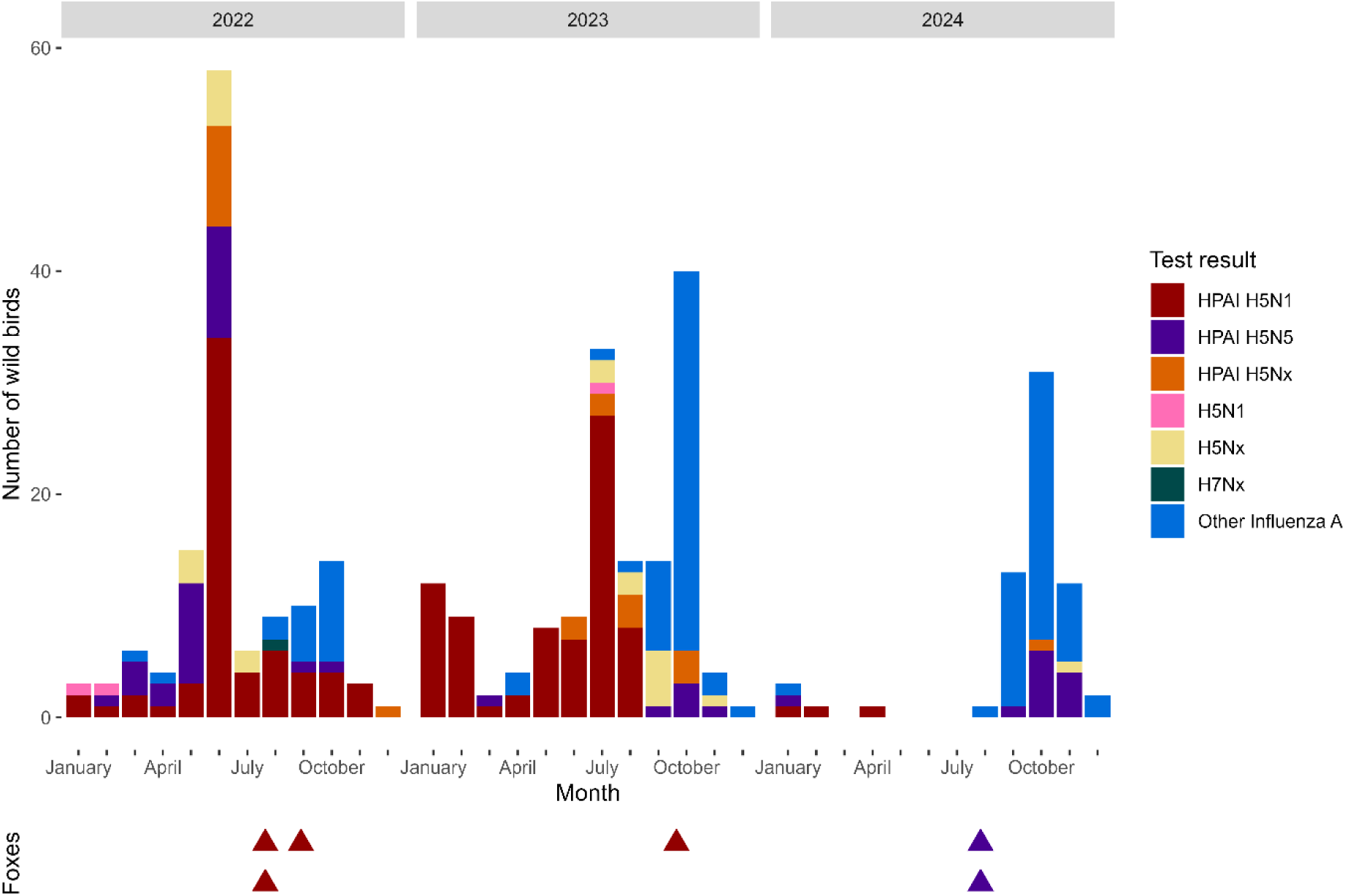
Number of red foxes (*Vulpes vulpes*) that tested positive for highly pathogenic avian influenza virus (HPAIV), 2022-2024, (triangles), in relation to wild birds from mainland Norway that tested positive during the same period (bars). Bars and triangles are color-coded by pathotype and subtype, as indicated in the legend on the right. Other InfA = non H5/H7 detections.

During the same period, 207 out of 2,393 wild birds tested through active and passive surveillance in mainland Norway were positive for HPAIV (Table 1, Figure 1).

**Table 1.**
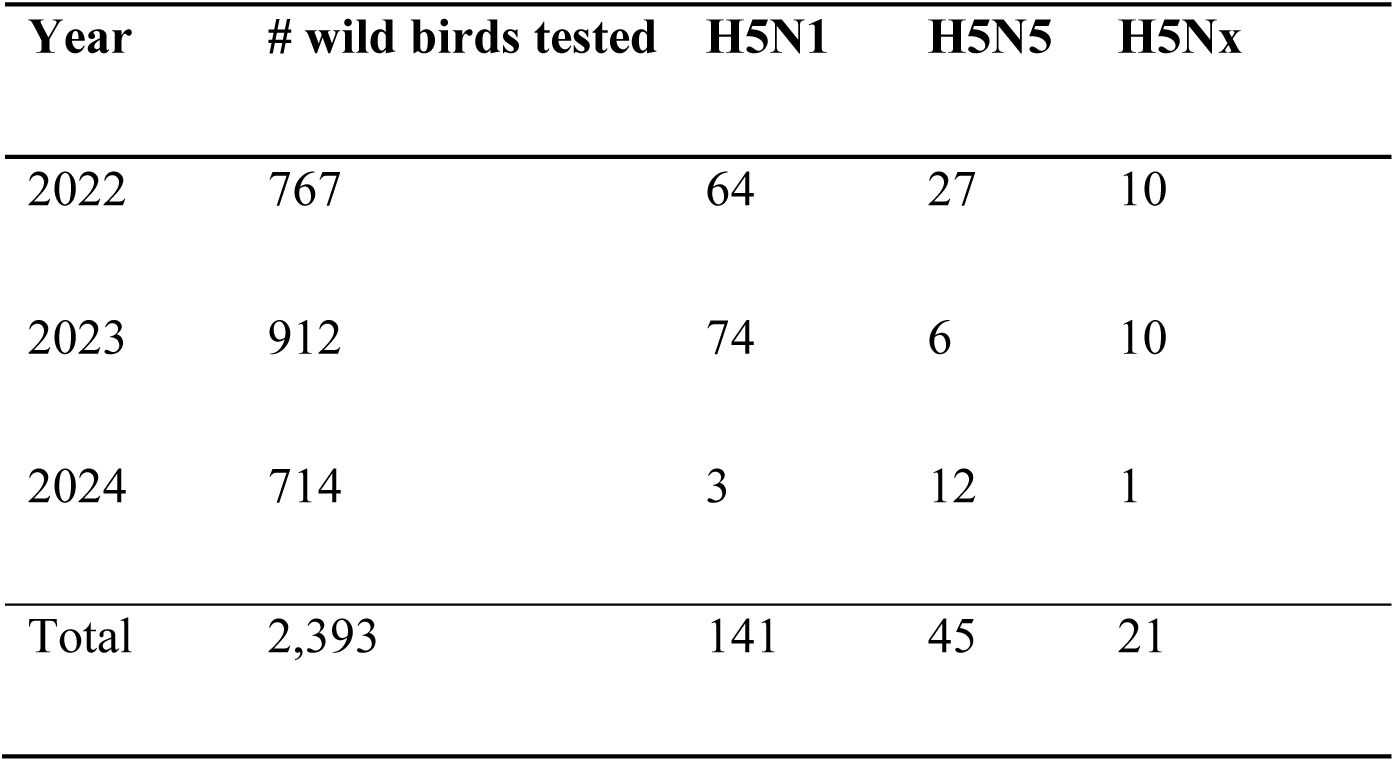
Detection of highly pathogenic avian influenza viruses H5N1 and H5N5 in wild birds from mainland Norway, 2022-2024, through active and passive surveillance. Cases were categorized as H5Nx when NA subtyping was not successful due to low viral load.

In 2022, three foxes and 64 wild birds tested positive for H5N1 (Figure 1, Table 1). The fox detections occurred one to two months after the summer peak of H5N1 detections in wild birds along the west coast of Norway, particularly affecting Northern gannets (*Morus bassanus*) (Figure 1 and 2). In 2023, one fox and 74 wild birds tested positive for H5N1. Again, the virus detection in the fox occurred a few months after a large outbreak of H5N1 in black-legged kittiwakes (*Rissa tridactyla*) and other gull species in Northern Norway (31). In February 2024, two foxes from Northern Norway tested positive for H5N5. These detections preceded a number of H5N5 cases subsequently identified in wild birds during October-December, mainly in large gull species (*Larus* spp.), hooded crows (*Corvus cornix*) and white-tailed eagles.

**Figure 2.**
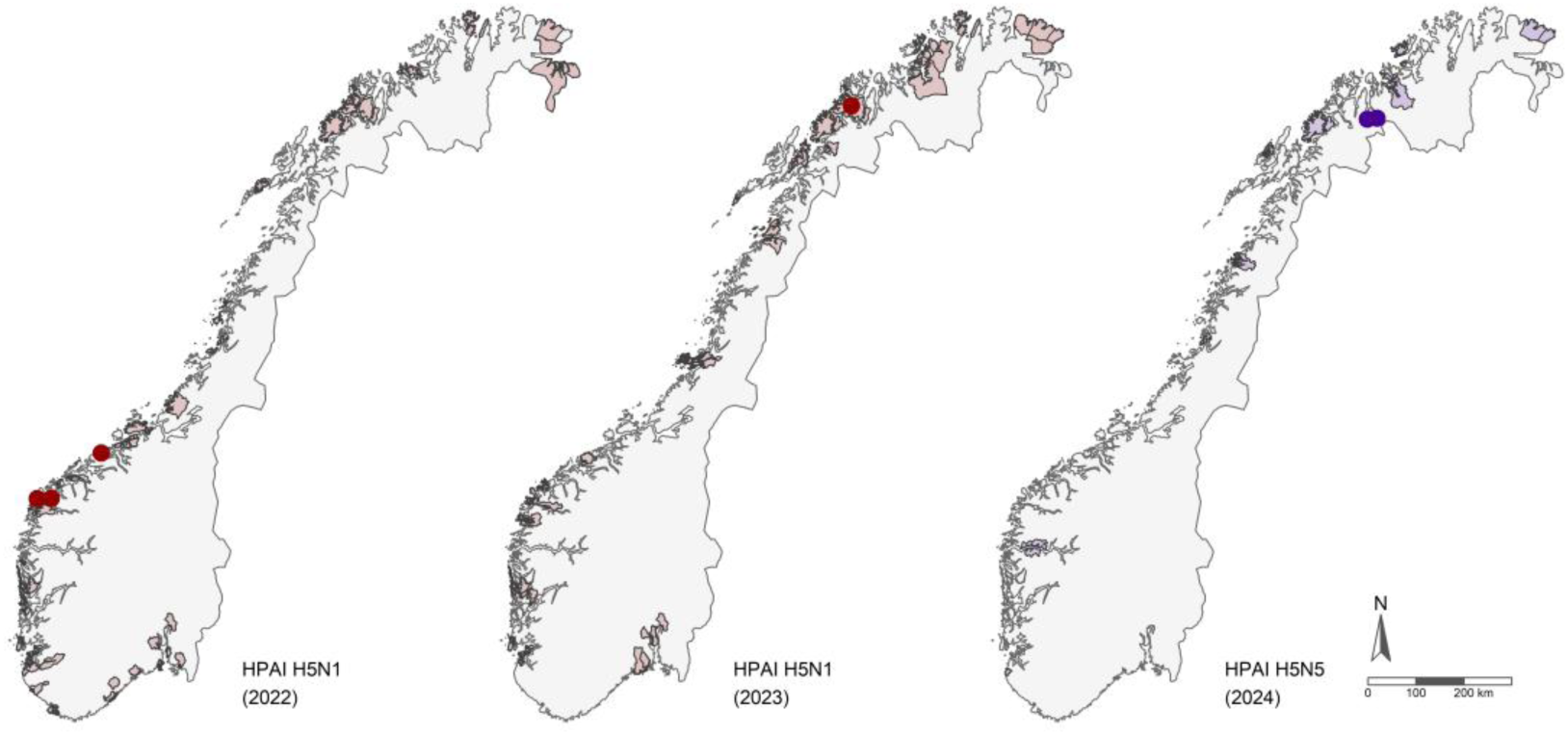
Maps showing detections of highly pathogenic avian influenza virus (HPAIV) of the H5N1 (red dots) and H5N5 (purple dots) subtypes in red foxes in Norway, 2022-2024. The corresponding-colored areas show municipalities where the same HPAIV subtypes were detected in wild birds during the same years.

### PCR Results from Swab Samples

Initial screening of tracheal and rectal swabs by rRT-PCR confirmed HPAIV in all six foxes (Table 2). Influenza A virus RNA was detected in tracheal swabs from all foxes, whereas rectal swabs were positive in only two cases. Genotyping identified H5Nx genotypes EA-2020-C (2022), H5N1 EA-2022-BB (2023), and H5N5 EA-2021-I (2024).

**Table 2.** Red foxes (*Vulpes vulpes*) positive for clade 2.3.4.4b highly pathogenic avian influenza viruses (H5N1 and H5N5) in Norway, 2022-2024. Initial screening results (influenza A virus rRT-PCR) in swab samples (T = trachea, R = rectum, N = nose), virus subtype and genotype, and detection of the mammalian-adaptation marker PB2:E627K are shown (ND = not detected). Whole-genome sequencing was performed on tracheal swabs.

| Fox # | Location year | Swab Cq | Subtype genotype | PB2 E627K | GISAID# |
| --- | --- | --- | --- | --- | --- |
| **1** | Selje, 2022 | T: 31.97<br>R: No Cq | H5N1,<br>EA-2020-C | 15.3%* | EPI_ISL_18901718 |
| **2** | Refvik, 2022 | T: 27.83<br>R: No Cq | H5N1,<br>EA-2020-C | ND | EPI_ISL_18455203 |
| **3** | Averøy, 2022 | T: 25.97<br>R: 34.31 | H5N1,<br>EA-2020-C | Present | EPI_ISL_18901720 |
| **4** | Tromsø, 2023 | T: 19.93<br>N: 28.97<br>R: 27.05 | H5N1,<br>EA-2022-BB | ND | EPI_ISL_19232456 |
| **5** | Skibotn 1, 2024 | T: 25.49<br>R: No Cq | H5N5,<br>EA-2021-I | ND | EPI_ISL_19232150 |
| **6** | Skibotn 2, 2024 | T: 19.74<br>R: No Cq | H5N5,<br>EA-2021-I | ND | EPI_ISL_19232151 |

\* Variant frequency 15.3% PB2: E627K in raw reads as determined by IRMA.

### Pathological Examination and Virus Distribution in Tissues

A summary of the pathological findings in the six foxes is provided in Appendix Table 1.

All foxes infected with H5N1 were juvenile males, whereas the two foxes infected with H5N5 were adult females. Foxes diagnosed with HPAI H5N1 in 2022 and 2023 were in moderate to low body condition (Figure 3A) and showed prominent pulmonary lesions characterized by dark red to purple lungs with marked edema, congestion and multifocal hemorrhages (Figure 3B). Both H5N5-infected foxes were adult females in moderate to moderate-low body condition. Fox Skibotn 2 (2024) had minimal pulmonary lesions, characterized by collapsed, brick red lungs with few small areas of hemorrhage in the right lung lobes (Figure 3C), whereas pulmonary assessment of Fox Skibotn 1 (2024) was not possible because of extensive gunshot trauma to the thorax. No consistent gross lesions were observed in other organs. Interpretation of gross pathology was complicated by freezing and thawing artifacts in the foxes from 2022 and 2023.

**Figure 3.**
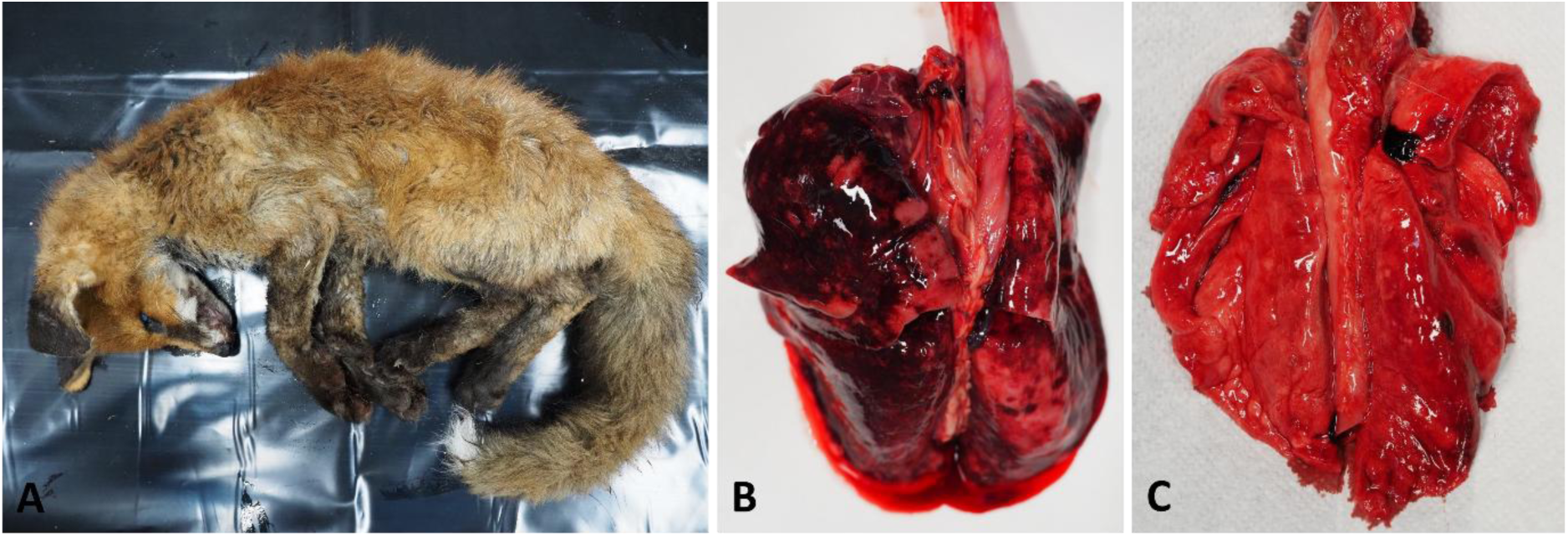
A) Fox Refvik (2022) was a juvenile male in moderate to low body condition. B) The lungs from Fox Refvik infected with HPAIV H5N1 were dark red and congested with marked edema. C) The lungs from Fox Skibotn 2 (2024) infected with HPAIV H5N5 were, in contrast, brick red and collapsed, with only a few small dark red spots.

Major histopathological changes were observed in the lungs and brain (Figure 4). Foxes with H5N1 infection showed a multifocal to diffuse fibrinonecrotizing pneumonia of varying severity (Figure 4, A-F). In the three foxes from 2022, inflammation was predominantly multifocal and peribronchiolar with mononuclear inflammatory infiltrates, bronchiolar necrosis, alveolar fibrinous exudation, and pulmonary necrosis. In Fox Tromsø (2023), the pulmonary lesions were diffuse rather than multifocal and were accompanied by widespread alveolar hemorrhages. Histological findings in this fox also suggested a concurrent inflammatory process in addition to the HPAIV-associated lesions. No histological changes were observed in the lungs of the H5N5-infected Fox Skibotn 2 (2024).

**Figure 4.**
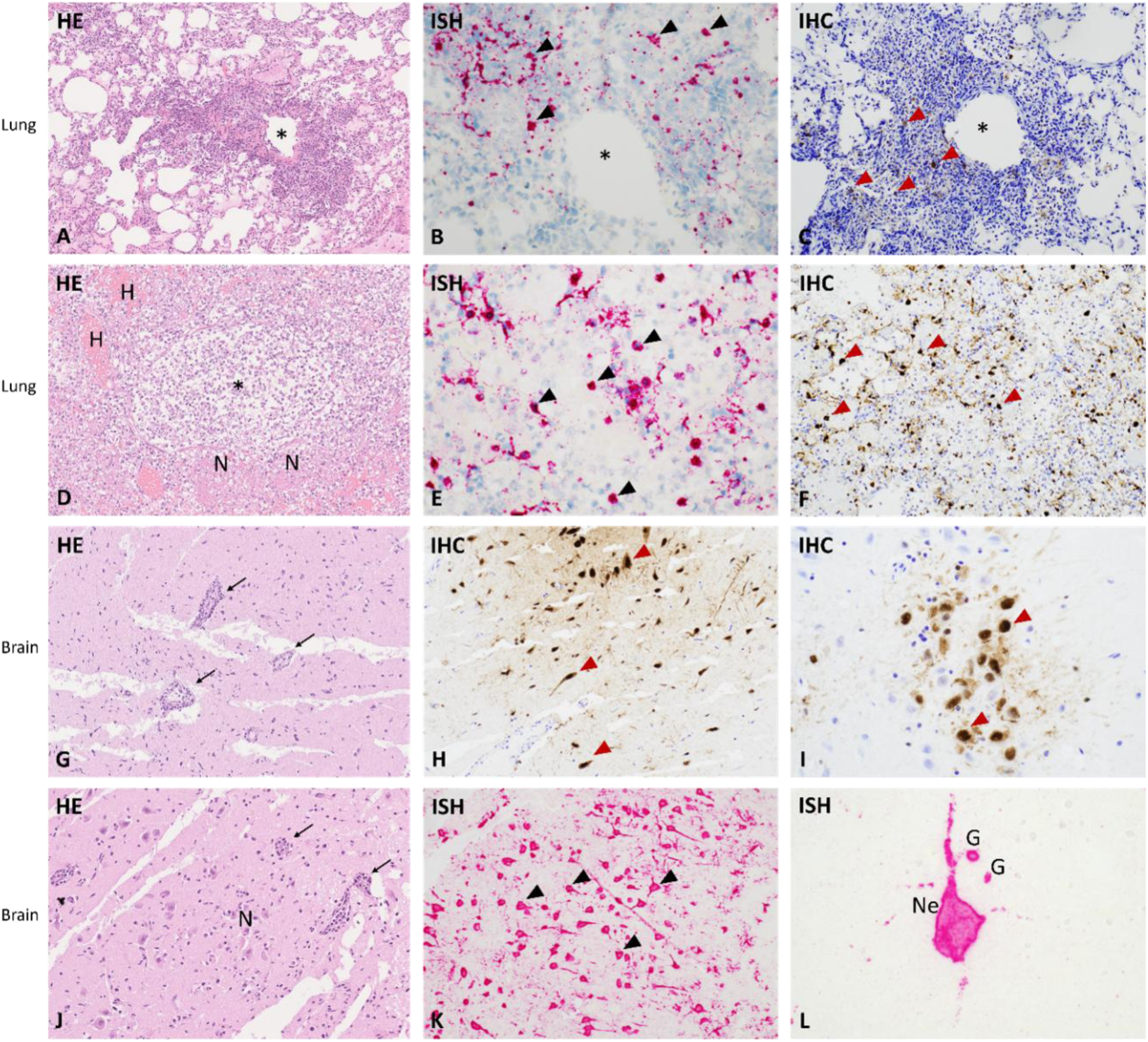
Histological changes and virus distribution in lung tissue (A-F) and brain tissue (G-L) of HPAIV H5N1 infected red foxes. Hematoxylin and eosin (HE), *in situ* hybridization (ISH) and immunohistochemistry (IHC). In the lungs of Fox Selje (2022) (A-C) there was multifocal and peribronchiolar infiltration of mononuclear inflammatory cells (A). Central bronchiole shown with *. Virus was detected within macrophages by ISH (black arrowheads) (B) and IHC (red arrowheads) (C). In Fox Tromsø (2023) (D-F), there were diffuse infiltrates of inflammatory cells within bronchioles (*) and alveoli, and multifocal alveolar hemorrhages (shown with an H) and necroses (shown with an N) (D). Virus was detected within macrophages by ISH (E) and IHC (F). In the brain of Fox Averøy (2022) (G-I), multifocal necrosis of gray matter and lymphohistiocytic perivascular cuffing (black arrows) were found (G). Virus was detected by IHC (H-I) in neurons (red arrowheads) and glia cells. Signal was predominately detected in the nuclei, but also in the cytoplasm of both neurons and glia cells. Endothelial cells and cells in perivascular cuffing were consistently negative. Similarly, in Fox Selje (2022) (J-L), multifocal necroses (shown with an N) of gray matter and perivascular cuffing were detected in the brain (black arrows) (J). By ISH (K-L), viral RNA was detected in neurons (black arrowheads) and glia cells in gray matter either in focal accumulations of neurons (K) or in single neurons (labeled as Ne) and glia cells (shown as G)(L). In slide G and J, white lines in the slides are artefacts.

Histopathological changes in the brain were detected in all examined foxes except Fox Tromsø (2023) and consisted of multifocal, subacute to chronic necrotizing encephalitis and vasculitis with neuronal necrosis, neuronophagia, inflammatory infiltrates and perivascular cuffing (Figure 4 G-L and Appendix Figure 1). Additional lesions were detected in the myocardium, liver and spleen of several animals (Appendix Figure 2), whereas no histopathological changes were observed in the pancreas or kidney.

Viral antigens and RNA detected by IHC and *in situ* hybridization, corresponded closely with the histopathological lesions. Inflammatory lesions in lungs and brain consistently contained viral antigen or RNA. Viral detection in other organs was limited, although viral signal was detected in the endothelium and intima of a single arteriole in Fox Averøy (2022). Fox Tromsø (2023) was negative on brain sections by both IHC and ISH.

### PCR Results from Tissue Samples

Influenza A virus M-gene RNA was detected by rRT-PCR in the brain and/or lung of all six red foxes (Table 3). The highest viral loads, estimated from Cq values, were detected in the brain in five of six animals. In Fox Tromsø (2023), the highest viral load was detected in the lung, and viral RNA was not detected in the brain. Influenza A virus was also detected in the trachea of all foxes, and in the anterior and posterior conchae from three foxes (Table 2 and 3). In all foxes, except Fox Tromsø (2023), viral RNA was detected in one or more internal organs including the liver, spleen, heart and kidney. Overall, tissue samples generally yielded lower Cq values than tracheal and rectal swab samples.

**Table 3.** Influenza A virus M gene rRT-PCR Cq values in samples from six wild red foxes naturally infected with clade 2.3.4.4b highly pathogenic avian influenza viruses in Norway, 2022-2024. Cq <25 indicate high, 25-30 moderate, and >30 low viral loads. No Cq indicates no detectable influenza A virus RNA.

| Fox ID/<br>Sample type | Selje<br>(2022) | Refvik<br>(2022) | Averøy<br>(2022) | Tromsø<br>(2023) | Skibotn 1<br>(2024) | Skibotn 2<br>(2024) |
| --- | --- | --- | --- | --- | --- | --- |
| Brain swab | 19.57 | 16.93* | 15.25 | NA | 20.31 | 19.36 |
| Brain | NA | NA | 16.91 | No Cq** | 26.50 | 22.98 |
| Tracheal swab*** | 27.2 | 30.73 | 20.05 | NA | 32.52 | 19.74 |
| Lung | 27.39 | 25.84 | 19.05 | 21.31 | 27.30 | 30.67 |
| Lung swab | 33.27 | 31.06 | 18.4 | NA | 25.64 | 24.90 |
| Liver | 33.59 | 28.74 | 21.72 | NA | No Cq | 31.03 |
| Rectal swab*** | No Cq | No Cq | No Cq | NA | No Cq | NA |
| Kidney | No Cq | No Cq | No Cq | NA | 33.03 | 30.97 |
| Spleen | No Cq | 28.81 | No Cq | NA | 26.85 | 29.64 |
| Intestine | No Cq | 31.27 | No Cq | NA | 34.85 | 29.76 |
| Ant. concha | NA | NA | 24.35 | NA | 28.45 | 20.15 |
| Post. concha | NA | NA | 25.19 | NA | 27.58 | 25.14 |
| Heart | NA | NA | 24.31 | NA | 34.23 | 31.81 |
| Lung lymph node | NA | NA | No Cq | 25.81 | NA | 26.75 |

NA, not tested or unsuitable sample, \* Sequenced deposited in GISAID (EPI\_ISL\_19232149),
\*\* Sample briefly exposed to formalin, \*\*\*Additional swabs collected after initial screening

### Phylogenetic Analyses

Phylogenetic analyses showed that the viruses detected in the foxes clustered closely with contemporaneous HPAIVs circulating in wild birds in Norway and Northern Europe (Figure 5A and Appendix Figures 3, 4 and 5). The H5N1 viruses detected in foxes in 2022 clustered with genotype EA-2020-C viruses circulating in Europe during the same period, including viruses detected in northern gannets (Figure 5A). The H5N1 virus detected in Fox Tromsø (2023) clustered with Norwegian gull-derived EA-2022-BB viruses primarily from northern Norway during summer 2023 (Figure 5B). The H5N5 viruses detected in foxes in 2024 were most closely related to EA-2021-I H5N5 viruses detected in gulls, crows, and a common eider (*Somateria mollissima*) in the Northern Hemisphere between autumn 2023 and winter 2024 (Figure 5C).

**Figure 5.**
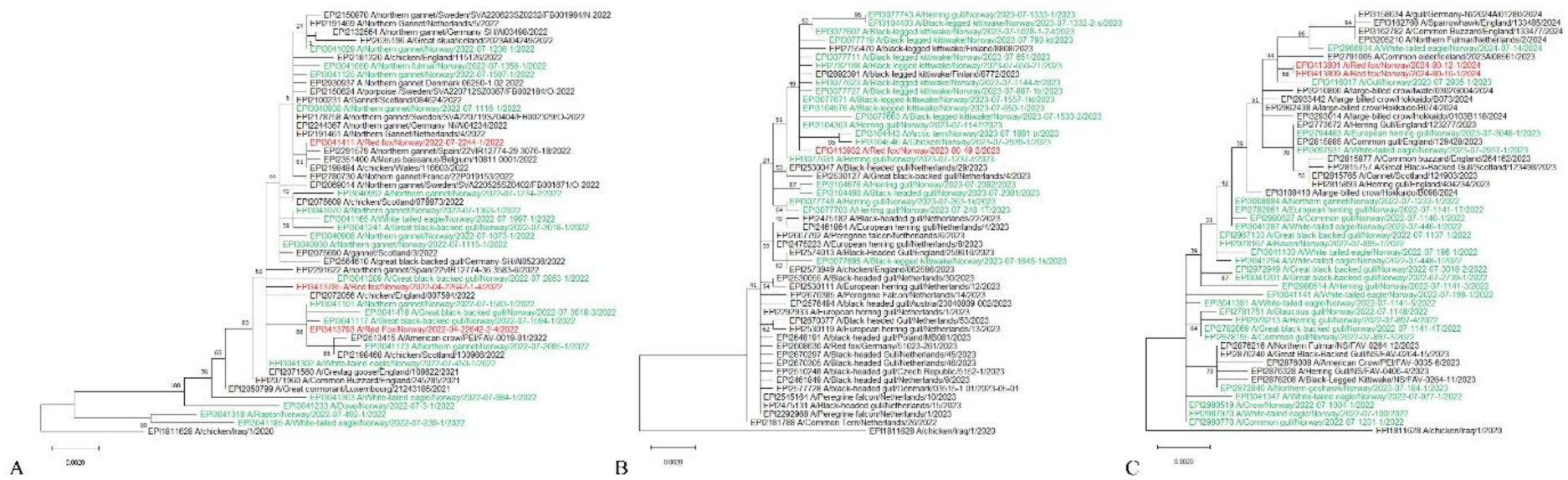
Genetic relationship of the HA segments of HPAI A(H5Nx) viruses obtained from red foxes (red) and wild birds (green) in Norway and reference sequences from Europe (black), 2022-2024. Maximum likelihood trees were constructed based on the coding region of the HA gene sequences for each genotype; A) H5N1 EA-2020-C, B) H5N1 EA-2022-BB, C) H5N5 EA-2021-I. Bootstrap support was calculated using 1000 replicates.

### Mutation Analysis

PB2:E627K, a recognized marker of mammalian adaptation, was detected in one H5N1 virus and as a minor variant in a second H5N1 virus from 2022 (Table 2 and Appendix Table 2). The same substitution was also identified in one H5N5 virus from a white-tailed eagle. Several additional substitutions previously linked to mammalian adaptation were identified in the HA protein, including S133A, T156A, K218Q and S223R. However, these substitutions were also present in avian viruses collected from the same geographic area and period. Complete mutation screening results are provided in Appendix Table 2.

## Discussion

HPAIV infection has been widely reported in red foxes and other carnivores across several continents (5). Because red foxes are opportunistic scavengers and predators with frequent exposure to sick and dead wild birds, they may serve as useful sentinel species for HPAI circulation. The six cases identified in Norway likely represent only a subset of infections occurring in red foxes. Recent serological evidence from red foxes in northeastern Germany and Ireland, together with findings from Arctic foxes in Svalbard, suggests that avian influenza virus exposure in wild canids may be more widespread than indicated by virus detection data alone (17, 18, 28). The extent of HPAIV exposure in Norwegian red foxes remains unknown, and further surveillance and serological studies are needed to better understand infection dynamics and outcomes in this species (17, 18).

Neurological disease is frequently reported in mammals infected with clade 2.3.4.4b HPAIV and reflects the marked neurotropism of viruses derived from the A/Goose/Guangdong/1/96 lineage (32). Clinical signs observed in the foxes described here, including circling, abnormal behaviour, lameness and reduced fear of humans, corresponded closely with the detection of high viral loads in the brain. Histopathological lesions and viral distribution were consistent with findings previously reported in red foxes and other carnivores from Europe and North America (13–16).

The close correspondence between histopathological lesions and viral detection by rRT-PCR, IHC and ISH supports HPAIV as the primary cause of disease in these foxes, although concurrent pathological processes could not be excluded in individual animals. Rabies is absent from mainland Norway (33), and canine distemper virus is only sporadically reported in Norwegian wild canids (34). Severe lesions were primarily confined to the lungs and central nervous system, although viral RNA was also detected in several peripheral organs, confirming systemic infection. In contrast, among the H5N5-infected foxes, the only animal with evaluable lung tissue showed minimal pulmonary pathology despite the presence of viral RNA. Viral RNA was detected less consistently in peripheral tissues of H5N5-infected foxes than in the H5N1-infected foxes. These observations may suggest differences in tissue distribution, pathogenesis or disease progression between the two subtypes, although the limited number of animals precludes firm conclusions. Interpretation of subtype-specific differences was constrained by the small sample size available for detailed pathological examination and the opportunistic nature of wildlife surveillance. Notably, all H5N1-infected foxes were juveniles, whereas both H5N5-infected foxes were adults. However, these observations are insufficient to draw conclusions regarding age-related susceptibility or disease severity. Although the study included relatively few foxes, the predominance of juvenile animals among the H5N1 cases is noteworthy. Similar observations have been reported in other mammals such as Arctic foxes infected with clade 2.3.4.4b HPAIVs (26), but it remains unclear whether this reflects increased susceptibility, greater exposure or potential surveillance bias.

Together with surveillance data from wild birds, the phylogenetic analyses strongly support that the infections described here represented independent spillover events from avian hosts rather than sustained transmission among foxes. The temporal association between fox detections and outbreaks in seabirds, combined with the close genetic relationship between fox and bird viruses, suggests that infection was most likely acquired through predation or scavenging on infected birds. The 2022 fox viruses clustered closely with H5N1 EA-2020-C viruses detected in northern gannets during the major North Atlantic seabird mortality event in 2022 (35). Notably, all three H5N1-infected foxes from 2022 were detected in coastal areas where HPAI-infected northern gannets had been found. Similarly, the H5N1 EA-2022-BB virus detected in Fox Tromsø (2023) was closely related to viruses circulating during the large outbreak among black-legged kittiwakes (*Rissa tridactyla*) and other gull species in northern Norway (31). The two H5N5-infected foxes were detected in the same area in northern Norway within a short time period, suggesting exposure to a common local source of infection.

The detection of H5N5 in two red foxes several months before most H5N5 detections subsequently recorded in wild birds in mainland Norway suggests that circulation of this subtype may have been more widespread than indicated by routine surveillance. A similar pattern was observed in Svalbard, where HPAIV H5N5 was detected in Arctic foxes in 2025 (26) and subsequently in a polar bear and a walrus (*Odobenus rosmarus rosmarus*) in 2026 (27), despite the absence of prior detections in local wild birds. Although H5N5 was later detected in large gulls on Jan Mayen in fall 2025, these observations suggest that infections in scavenging mammals may help identify otherwise undetected virus circulation in wildlife populations. Together with previous findings in white-tailed eagles (20), and subsequent detections of H5N5 in multiple mammalian species across mainland Norway and Svalbard (36), this pattern suggest that scavenging birds and mammals may provide complementary information for HPAIV surveillance. The occurrence of H5N5 across diverse host species and geographic regions also raises questions about the longer-term ecology and evolution of this subtype. Notably, an additional H5N5 infection in a red fox was reported from Troms, northern Norway, in April 2025 after completion of the present study, further indicating continued spillover of this genotype into mammalian hosts.

An intriguing feature of the EA-2021-I H5N5 genotype is its persistence across multiple years and host species despite limited documented changes in genome constellation. Its repeated detection contrasts with the emergence of several H5N1 genotypes in Northern Europe during the same period. Continued genomic surveillance may help clarify the ecological and evolutionary mechanisms underlying the apparent stability of this genotype.

Mutation analysis identified the mammalian adaptation marker PB2:E627K in one H5N1 virus and as a minority variant in a second H5N1 virus. Detection of PB2:E627K as a minority variant may represent an intermediate state of within host adaptation following avian-to-mammalian transmission. Experimental studies have demonstrated rapid selection of this substitution after infection of mammals (37), supporting the hypothesis that adaptive evolution can occur shortly after spillover from birds. PB2:E627K has repeatedly been reported in HPAI-infected carnivores and other mammals (6, 13) and remains one of the most important genetic markers associated with adaptation to mammalian hosts (38). Notably, PB2:E627K was also detected in a contemporaneous H5N5 virus from a white-tailed eagle in Norway, indicating that this marker was not unique to the fox-derived viruses. In contrast, the HA substitutions identified in the fox viruses were also present in contemporary avian viruses from the same geographic region and period and therefore did not provide evidence of host-specific adaptation.

In conclusion, HPAIV infection in Norwegian red foxes was associated with severe systemic disease characterized by prominent involvement of the central nervous system and lungs. Phylogenetic and epidemiological evidence supports that these cases represented independent spillover events from contemporaneously circulating avian viruses most likely acquired through predation or scavenging. The detection of PB2:E627K in one virus and as a minority variant in another demonstrates the capacity of these viruses to acquire adaptive changes following transmission to mammals. These findings highlight the importance of continued surveillance of both avian and mammalian species to detect spillover events, monitor viral adaptation following cross-species transmission, and assess potential animal and public health risks.

## Acknowledgements

We gratefully acknowledge all data contributors, i.e., the Authors and their Originating laboratories responsible for obtaining the specimens, and their Submitting laboratories for generating the genetic sequence and metadata and sharing via the GISAID Initiative, on which this research is based. We thank the Norwegian Food Safety Authority (NFSA) and other collaborators for collecting and submitting carcasses and samples to the Norwegian Veterinary Institute (NVI). We are especially grateful to the technical and laboratory staff at NVI for their extensive support throughout the investigation, including specimen reception and processing, assistance during necropsies, sample preparation, laboratory diagnostics, biosafety assessments and advice. We also thank Rasmus Kopperud Riis, Norwegian Institute of Public Health (NIPH), for developing and sharing the script used to screen viral sequences for markers of mammalian adaptation. Microsoft Copilot was used for language editing and improvement of readability. The authors take full responsibility for all scientific content, interpretations, and conclusions.

## Ethics Statement

The study was based on animals submitted through routine wildlife disease surveillance. No animals were euthanized for research purposes, and no experimental procedures were performed. The study therefore did not require approval by an animal research ethics committee.

## Data availability

Genome sequences generated in this study have been deposited in the GISAID EpiFlu database under the accession numbers listed in Table 2 and Appendix Table 2. The virus sequences included in the phylogenetic analysis are provided in Supplementary File 1. Additional data, including raw sequencing data, supporting the findings of this study are available from the corresponding author upon reasonable request.

