## Appendix for "Highly Pathogenic Avian Influenza Viruses H5N1 and H5N5 in Red Foxes (*Vulpes vulpes*) in Norway during 2022-2024"

### Appendix Materials and Methods

#### *Sample Collection and Virus Detection*

During 2022-2024, six sick or dead red foxes were reported to and sampled by inspectors from the Norwegian Food Safety Authority (NFSA). For initial testing, swabs were collected from trachea and rectum and placed in transport medium (Amies). In addition, a nasal swab was collected from one fox. The samples were sent by overnight express parcel service to the Norwegian Veterinary Institute (NVI) for analyses by PCR.

Nucleic acids were extracted on a MagNA Pure 24 or 96 (Roche), and real-time reverse transcriptase-PCR (rRT-PCR) targeting the matrix (M) gene of the influenza A virus was used for viral RNA detection. Influenza A positive samples were further characterized by subtype-specific rRT-PCRs, which also includes determination of pathogenicity, as described in Bøe *et al.* (1).

#### *Surveillance Systems, Data Extraction and Visualization*

Samples from sick or dead wild birds and other animals were collected all-year round by NFSA as part of passive surveillance. In addition, samples from hunted wild aquatic birds were collected during autumn (active surveillance). Data on HPAIV detections in mainland Norway were extracted from the Laboratory Information Management System at NVI using R (R Core Team, 2025). Several R packages were employed for data processing and visualization. A figure was created using the ggplot2 package (2), showing both positive wild birds and red foxes, while the tmap package (3) was used to visualize municipalities with positive cases in wild birds and to map the locations of infected red foxes. Spatial data processing was carried out using the sf package (4) which enabled the handling and manipulation of geographic data for accurate mapping and analysis.

*Pathological Investigation*

For pathological investigations, the six fox carcasses were frozen locally and either transported by car or sent by overnight express parcel service to the NVI laboratories in Tromsø or Ås. The foxes were either stored at –20°C and later thawed for necropsy or directly transferred to a BSL3 facility for necropsy and sampling following standardized procedures. Swabs for HPAIV confirmation by PCR were collected from the trachea, lung and brain. To assess virus distribution, samples from brain, lung, trachea, liver, spleen, kidney, intestine, heart, lung-associated lymph node, and, when available, anterior and posterior nasal conchae were collected for PCR analyses. Representative tissue samples from major organs, including brain, lung, heart, liver, spleen, kidney and pancreas, were collected for histopathology. Because sample availability varied among foxes, not all tissues were examined in every animal.

For PCR analyses, tissues were preserved in RNAlater® (Thermo Fisher Scientific, Waltham, MA, USA) or transport medium and stored at –80°C until further processed and analyzed. Nucleic acids were extracted as previously described.

For histology, tissues were fixed in 10% buffered formalin, dehydrated and embedded in paraffin, sectioned and stained with hematoxylin and eosin according to standardized procedures before examination by light microscopy.

*Virus Detection and Visualization by RNAscope™ and Immunohistochemistry*

Formalin fixated, paraffin-embedded tissue sections were utilized to detect viral RNA expression using the RNAScope™ *in situ* hybridization (ISH) technique. RNAscope ISH enables the visualization of individual RNA molecules in cytological and histological samples at the single-cell level (5).

We employed a probe that targets the M gene of Avian Influenza A Virus (AIAV), which is conserved across all influenza A viruses. This probe was procured from ACD Bio (2.5 LS probe-V-Influenza A-H5N8-M2M1 C1, Cat. Nr. 1048228-C1). The slides were stained using the RNAscope LS Red automated kit (ACD Bio, Cat. Nr. 322750) on a Leica BOND RXm from Leica Biosystems. As a negative control, we used a probe targeting the dihydrodipicolinate reductase (*dapB*) gene from *Bacillus subtilis*.

For immunohistochemical (IHC) analysis of tissues from all foxes except Fox Tromsø (2023), sections were deparaffinized and rehydrated prior to proteinase K treatment for 6 minutes and peroxidase block for 5 minutes. Sections were incubated at room temperature for 30 minutes with an anti-influenza A primary monoclonal antibody diluted 1:1000 (Progen, EBS-I-238). For detection, EnVision+/HRP was added for 30 minutes followed by DAB+ for 10 minutes. Washing was performed between each step. Sections were counterstained with hematoxylin, dehydrated and mounted.

For immunohistochemical (IHC) analysis of tissues from Fox Tromsø (2023), sections were deparaffinized and rehydrated prior to proteinase K treatment for 30 minutes and peroxidase block for 10 minutes. Sections were incubated at room temperature for 60 minutes with an anti-influenza A primary monoclonal antibody diluted 1:2000 (Statens Serum Institut, HYB 340-05). For detection, EnVision HRP was added for 30 minutes followed by Romulin AEX Chromogen for 15 minutes. Washing was performed between each step. Sections were counterstained with hematoxylin, dehydrated and mounted.

*Whole-Genome Sequencing, Assembly and Genotyping*

Samples were subjected to cDNA synthesis as described by Zhou et al. (6) before whole-genome sequencing (WGS) was performed as described (1). DNA libraries were prepared from 200 to 400 ng DNA using Illumina DNA Prep (Illumina, San Diego, CA, USA) and sequenced using Illumina NextSeq/MiSeq (Illumina).

Consensus sequences for the viral genomes were obtained using IRMA (Iterative Refinement Meta-Assembler) v1.0.3 with default settings (7). The consensus sequences and metadata were uploaded to GISAID EpiFlu (8). Genotyping was performed as described by Fusaro et al.(9). Briefly, each viral segment and the Eurasian avian (EA) reference sequences provided by the EURL were included in cluster analysis performed using MEGA X v10.1.8.

*Phylogenetic Analyses*

The genetic similarity of all eight segments from the six fox-derived viruses was assessed using BLAST in GISAID EpiFlu. For each of the eight viral segments, the full-length coding nucleotide sequences were aligned by MUSCLE, and phylogenetic trees were constructed using the Maximum Likelihood algorithm and the Tamura-Nei model in MEGA X v10.1.8 (10). Bootstrap values of 1000 were used to assess nodal support. For each segment, the 50 most genetically similar sequences in GISAID, along with available H5N1 and H5N5 sequences from Norway from the relevant period, were included in the initial analysis. The final sequences included in the trees are listed in Supplementary File 1. All sequences and associated metadata included in the analyses are also available at https://doi.org/10.55876/gis8.260923pq (Supplementary File 2).

*Mutation Analysis*

The FluSeq script (https://github.com/RasmusKoRiis/nf-core-fluseq/tree/master) processes FASTQ files using IRMA's IRMA-minion module to generate consensus sequences with ambiguous bases at positions where the major base frequency is under 70% (https://wonder.cdc.gov/amd/flu/irma/). Subtype identification is done by aligning hemagglutinin (HA) and neuraminidase (NA) sequences to a reference FASTA with multiple subtypes (https://github.com/epi2me-labs/wf-flu/blob/master/data/primer_schemes/V1/consensus_irma.fasta). Consensus sequences were translated to amino acids using Nextclade (11) and mutation analysis compared these sequences to a reference using Python (v. 3.9.19). Mutations were evaluated against a curated list of subtype-relevant mutations (12). Key references included A/Viet Nam/1203/2004 H5N1 for HA, A/Aichi/2/1968 (H3N2) for NA, and A/Goose/Guangdong/1/96 H5N1 for all the other segments.

### Appendix Figures

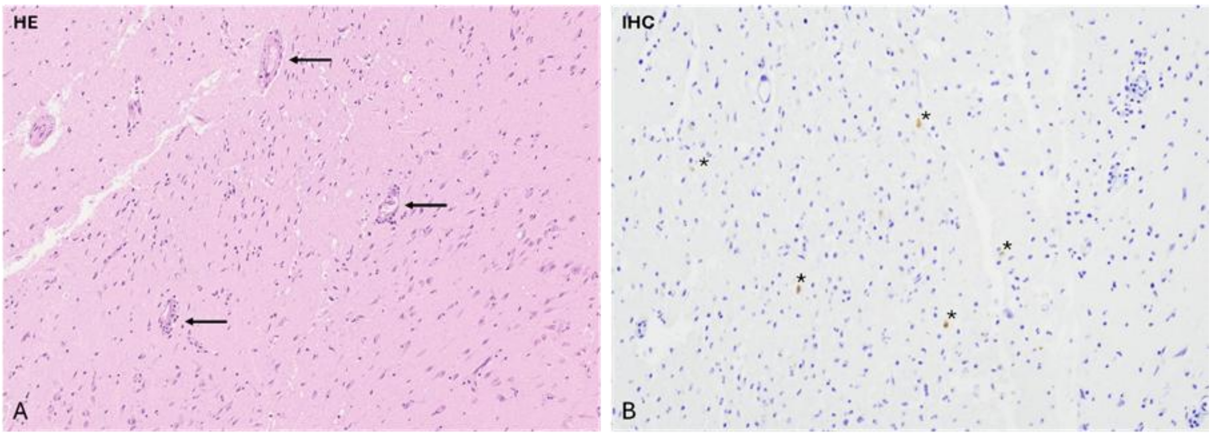

**Appendix Figure 1**: Histological changes and virus distribution in brain tissue from the H5N5 infected Fox Skibotn 1 (2024). A) Hematoxylin and eosin staining (HE). Multifocal mild lymphohistiocytic perivascular cuffing was detected in some parts of the brain (black arrows). Few areas of increased cell infiltration were seen in the brain of this H5N5 infected fox (not shown). B) Immunohistochemistry (IHC). Few scattered single neurons were positive for Influenza A (marked by *).

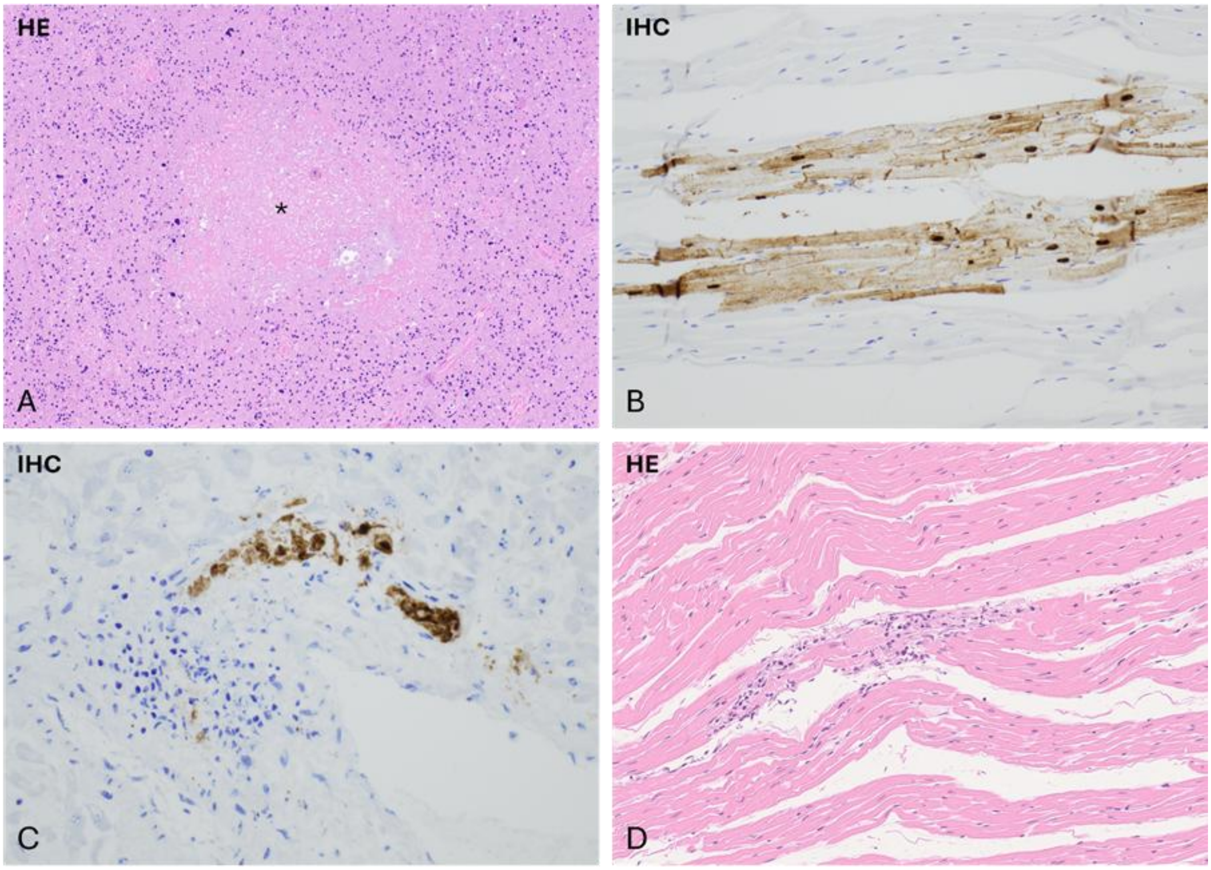

**Appendix Figure 2**: Histological changes and virus distribution in various organs from HPAIV infected red foxes. Hematoxylin and eosin (HE) and immunohistochemistry (IHC). A-C) spleen, heart and liver from Fox Averøy (2022) (H5N1). D) Heart from Fox Skibotn 1 (2024) (H5N5). In the spleen (A), multifocal necroses of various sizes were noted (*). Positive signal was detected in necrotic matter by IHC (not shown). By IHC, virus was detected in the myofibers in the myocardium of the heart (B). Virus was detected in the macrophages within triades in the liver (C). No histological changes were present by investigation of HE slides of the heart and liver from Fox Averøy (2022). In the heart from Fox Skibotn 1 (2024), multifocal degeneration and mononuclear cell accumulation were detected in the myocardium (D). No virus was detected by IHC (not shown).

**Appendix Figure 3.** Midpoint rooted maximum likelihood phylogenetic trees showing the genetic relationship between highly pathogenic avian influenza H5N1 EA-2020-C viruses from red foxes (red), wild birds (green) and other highly similar viruses obtained from GISAID (black), 2022. Trees are shown for the gene segments coding A) PB2, B) PB1, C) PA, D) NP, E) NA, F) M and G) NS.

A)

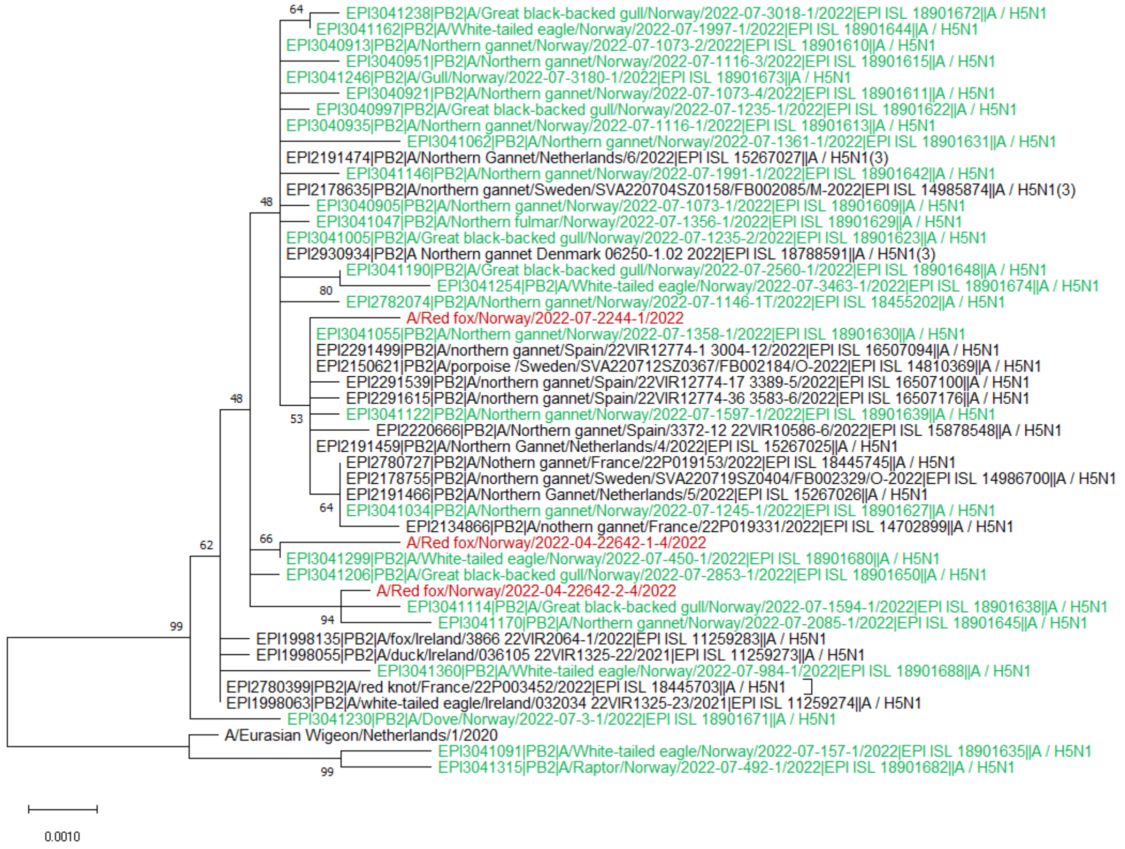

B)

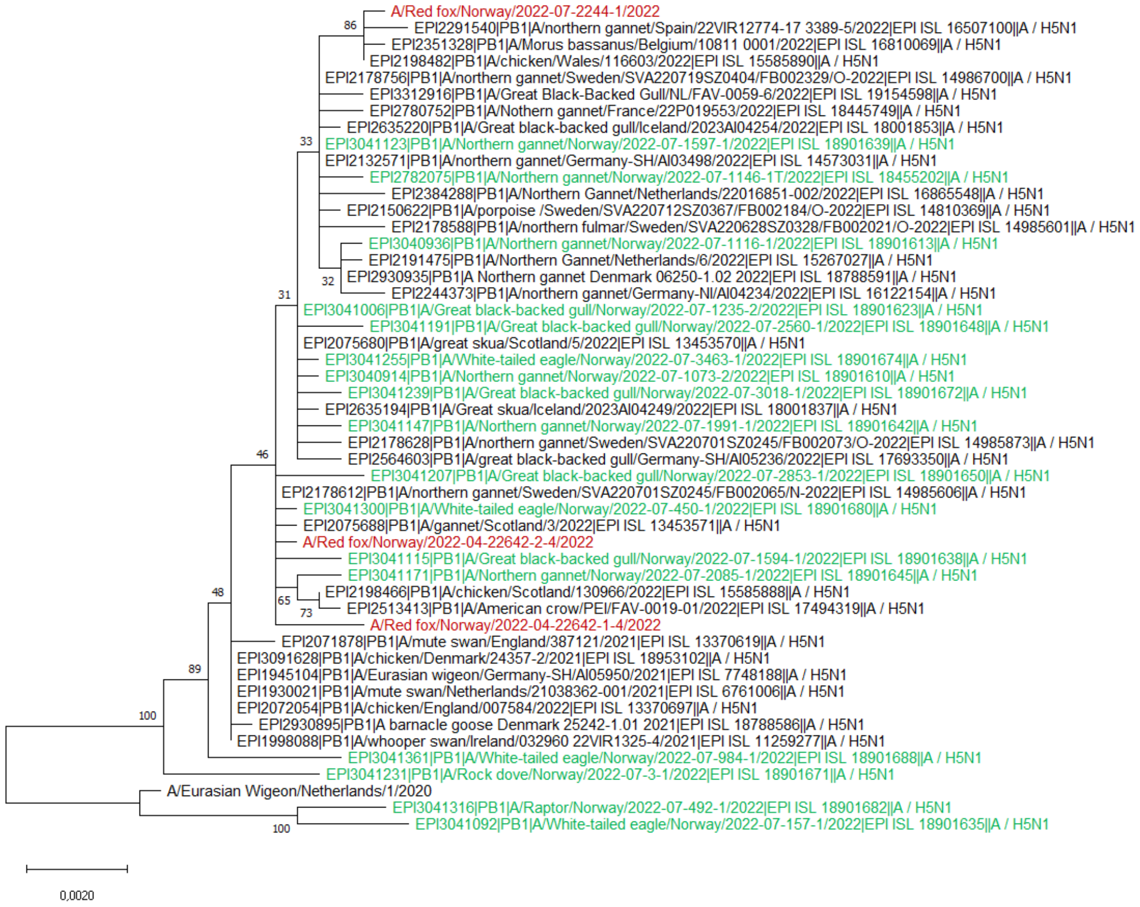
A/Red fox/Norway/2022-07-2244-1/2022
86
EPI2291540|PB1|A/northern gannet/Spain/22VIR12774-17 3389-5/2022|EPI ISL 16507100||A / H5N1
EPI2351328|PB1|A/Morus bassanus/Belgium/10811 0001/2022|EPI ISL 16810069||A / H5N1
EPI2198482|PB1|A/chicken/Wales/116603/2022|EPI ISL 15585890||A / H5N1
EPI2178756|PB1|A/northern gannet/Sweden/SVA220719SZ0404/FB002329/O-2022|EPI ISL 14986700||A / H5N1
EPI3312916|PB1|A/Great Black-Backed Gull/NL/FAV-0059-6/2022|EPI ISL 19154598||A / H5N1
EPI2780752|PB1|A/Nothern gannet/France/22P019553/2022|EPI ISL 18445749||A / H5N1
EPI2635220|PB1|A/Great black-backed gull/Iceland/2023AI04254/2022|EPI ISL 18001853||A / H5N1
33
EPI3041123|PB1|A/Northern gannet/Norway/2022-07-1597-1/2022|EPI ISL 18901639||A / H5N1
EPI2132571|PB1|A/northern gannet/Germany-SH/AI03498/2022|EPI ISL 14573031||A / H5N1
EPI2782075|PB1|A/Northern gannet/Norway/2022-07-1146-1T/2022|EPI ISL 18455202||A / H5N1
EPI2384288|PB1|A/Northern Gannet/Netherlands/22016851-002/2022|EPI ISL 16865548||A / H5N1
EPI2150622|PB1|A/porpoise /Sweden/SVA220712SZ0367/FB002184/O-2022|EPI ISL 14810369||A / H5N1
EPI2178588|PB1|A/northern fulmar/Sweden/SVA220628SZ0328/FB002021/O-2022|EPI ISL 14985601||A / H5N1
EPI3040936|PB1|A/Northern gannet/Norway/2022-07-1116-1/2022|EPI ISL 18901613||A / H5N1
EPI2191475|PB1|A/Northern Gannet/Netherlands/6/2022|EPI ISL 15267027||A / H5N1
32
EPI2930935|PB1|A Northern gannet Denmark 06250-1.02 2022|EPI ISL 18788591||A / H5N1
EPI2244373|PB1|A/northern gannet/Germany-NI/AI04234/2022|EPI ISL 16122154||A / H5N1
31
EPI3041006|PB1|A/Great black-backed gull/Norway/2022-07-1235-2/2022|EPI ISL 18901623||A / H5N1
EPI3041191|PB1|A/Great black-backed gull/Norway/2022-07-2560-1/2022|EPI ISL 18901648||A / H5N1
EPI2075680|PB1|A/great skua/Scotland/5/2022|EPI ISL 13453570||A / H5N1
EPI3041255|PB1|A/White-tailed eagle/Norway/2022-07-3463-1/2022|EPI ISL 18901674||A / H5N1
EPI3040914|PB1|A/Northern gannet/Norway/2022-07-1073-2/2022|EPI ISL 18901610||A / H5N1
EPI3041239|PB1|A/Great black-backed gull/Norway/2022-07-3018-1/2022|EPI ISL 18901672||A / H5N1
EPI2635194|PB1|A/Great skua/Iceland/2023AI04249/2022|EPI ISL 18001837||A / H5N1
EPI3041147|PB1|A/Northern gannet/Norway/2022-07-1991-1/2022|EPI ISL 18901642||A / H5N1
EPI2178628|PB1|A/northern gannet/Sweden/SVA220701SZ0245/FB002073/O-2022|EPI ISL 14985873||A / H5N1
EPI2564603|PB1|A/great black-backed gull/Germany-SH/AI05236/2022|EPI ISL 17693350||A / H5N1
46
EPI3041207|PB1|A/Great black-backed gull/Norway/2022-07-2853-1/2022|EPI ISL 18901650||A / H5N1
EPI2178612|PB1|A/northern gannet/Sweden/SVA220701SZ0245/FB002065/N-2022|EPI ISL 14985606||A / H5N1
EPI3041300|PB1|A/White-tailed eagle/Norway/2022-07-450-1/2022|EPI ISL 18901680||A / H5N1
EPI2075688|PB1|A/gannet/Scotland/3/2022|EPI ISL 13453571||A / H5N1
A/Red fox/Norway/2022-04-22642-2-4/2022
EPI3041115|PB1|A/Great black-backed gull/Norway/2022-07-1594-1/2022|EPI ISL 18901638||A / H5N1
EPI3041171|PB1|A/Northern gannet/Norway/2022-07-2085-1/2022|EPI ISL 18901645||A / H5N1
48
65
EPI2198466|PB1|A/chicken/Scotland/130966/2022|EPI ISL 15585888||A / H5N1
73
EPI2513413|PB1|A/American crow/PEI/FAV-0019-01/2022|EPI ISL 17494319||A / H5N1
A/Red fox/Norway/2022-04-22642-1-4/2022
EPI2071878|PB1|A/mute swan/England/387121/2021|EPI ISL 13370619||A / H5N1
EPI3091628|PB1|A/chicken/Denmark/24357-2/2021|EPI ISL 18953102||A / H5N1
89
EPI1945104|PB1|A/Eurasian wigeon/Germany-SH/AI05950/2021|EPI ISL 7748188||A / H5N1
EPI1930021|PB1|A/mute swan/Netherlands/21038362-001/2021|EPI ISL 6761006||A / H5N1
EPI2072054|PB1|A/chicken/England/007584/2022|EPI ISL 13370697||A / H5N1
100
EPI2930895|PB1|A barnacle goose Denmark 25242-1.01 2021|EPI ISL 18788586||A / H5N1
EPI1998088|PB1|A/whooper swan/Ireland/032960 22VIR1325-4/2021|EPI ISL 11259277||A / H5N1
EPI3041361|PB1|A/White-tailed eagle/Norway/2022-07-984-1/2022|EPI ISL 18901688||A / H5N1
EPI3041231|PB1|A/Rock dove/Norway/2022-07-3-1/2022|EPI ISL 18901671||A / H5N1
A/Eurasian Wigeon/Netherlands/1/2020
EPI3041316|PB1|A/Raptor/Norway/2022-07-492-1/2022|EPI ISL 18901682||A / H5N1
100
EPI3041092|PB1|A/White-tailed eagle/Norway/2022-07-157-1/2022|EPI ISL 18901635||A / H5N1
0,0020

C)

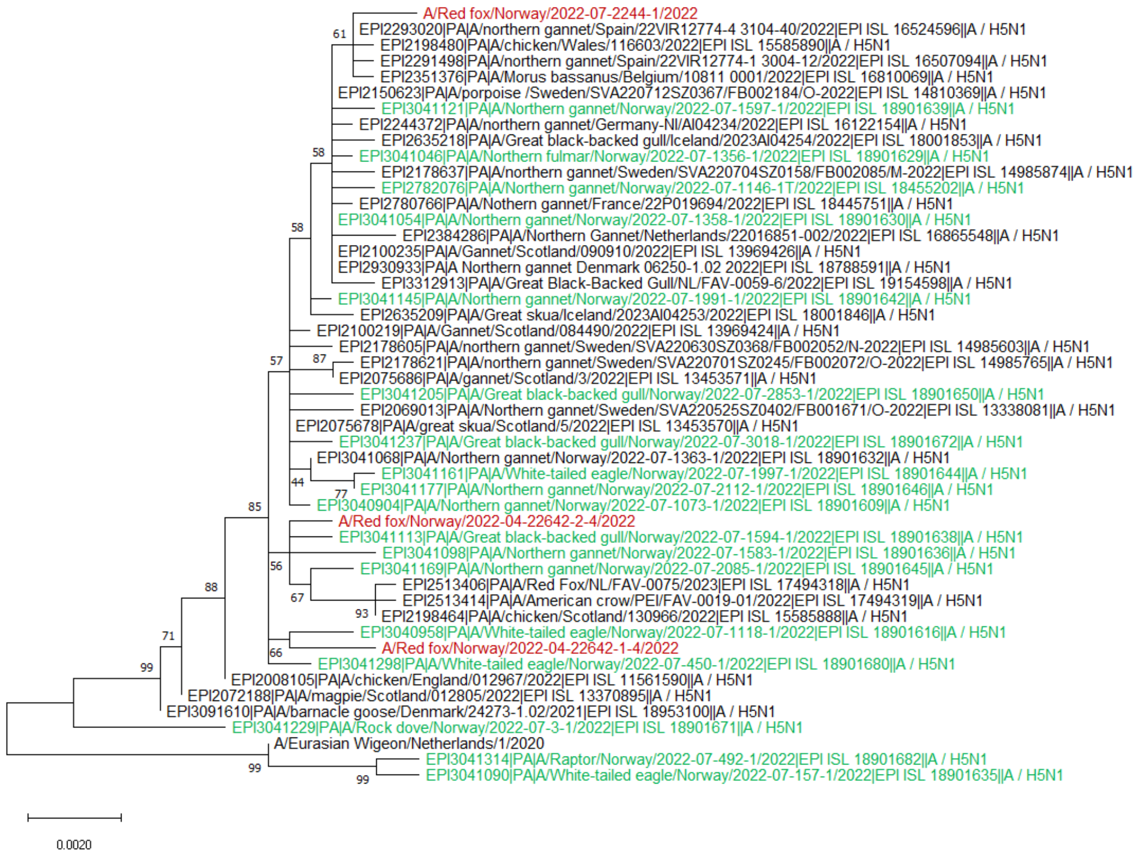
A/Red fox/Norway/2022-07-2244-1/2022
EPI2293020|PA|A/northern gannet/Spain/22VIR12774-4 3104-40/2022|EPI ISL 16524596||A / H5N1
EPI2198480|PA|A/chicken/Wales/116603/2022|EPI ISL 15585890||A / H5N1
EPI2291498|PA|A/northern gannet/Spain/22VIR12774-1 3004-12/2022|EPI ISL 16507094||A / H5N1
EPI2351376|PA|A/Morus bassanus/Belgium/10811 0001/2022|EPI ISL 16810069||A / H5N1
EPI2150623|PA|A/porpoise /Sweden/SVA220712SZ0367/FB002184/O-2022|EPI ISL 14810369||A / H5N1
EPI3041121|PA|A/Northern gannet/Norway/2022-07-1597-1/2022|EPI ISL 18901639||A / H5N1
EPI2244372|PA|A/northern gannet/Germany-NI/AI04234/2022|EPI ISL 16122154||A / H5N1
EPI2635218|PA|A/Great black-backed gull/Iceland/2023AI04254/2022|EPI ISL 18001853||A / H5N1
EPI3041046|PA|A/Northern fulmar/Norway/2022-07-1356-1/2022|EPI ISL 18901629||A / H5N1
EPI2178637|PA|A/northern gannet/Sweden/SVA220704SZ0158/FB002085/M-2022|EPI ISL 14985874||A / H5N1
EPI2782076|PA|A/Northern gannet/Norway/2022-07-1146-1T/2022|EPI ISL 18455202||A / H5N1
EPI2780766|PA|A/Nothern gannet/France/22P019694/2022|EPI ISL 18445751||A / H5N1
EPI3041054|PA|A/Northern gannet/Norway/2022-07-1358-1/2022|EPI ISL 18901630||A / H5N1
EPI2384286|PA|A/Northern Gannet/Netherlands/22016851-002/2022|EPI ISL 16865548||A / H5N1
EPI2100235|PA|A/Gannet/Scotland/090910/2022|EPI ISL 13969426||A / H5N1
EPI2930933|PA|A Northern gannet Denmark 06250-1.02 2022|EPI ISL 18788591||A / H5N1
EPI3312913|PA|A/Great Black-Backed Gull/NL/FAV-0059-6/2022|EPI ISL 19154598||A / H5N1
EPI3041145|PA|A/Northern gannet/Norway/2022-07-1991-1/2022|EPI ISL 18901642||A / H5N1
EPI2635209|PA|A/Great skua/Iceland/2023AI04253/2022|EPI ISL 18001846||A / H5N1
EPI2100219|PA|A/Gannet/Scotland/084490/2022|EPI ISL 13969424||A / H5N1
EPI2178605|PA|A/northern gannet/Sweden/SVA220630SZ0368/FB002052/N-2022|EPI ISL 14985603||A / H5N1
EPI2178621|PA|A/northern gannet/Sweden/SVA220701SZ0245/FB002072/O-2022|EPI ISL 14985765||A / H5N1
EPI2075686|PA|A/gannet/Scotland/3/2022|EPI ISL 13453571||A / H5N1
EPI3041205|PA|A/Great black-backed gull/Norway/2022-07-2853-1/2022|EPI ISL 18901650||A / H5N1
EPI2069013|PA|A/Northern gannet/Sweden/SVA220525SZ0402/FB001671/O-2022|EPI ISL 13338081||A / H5N1
EPI2075678|PA|A/great skua/Scotland/5/2022|EPI ISL 13453570||A / H5N1
EPI3041237|PA|A/Great black-backed gull/Norway/2022-07-3018-1/2022|EPI ISL 18901672||A / H5N1
EPI3041068|PA|A/Northern gannet/Norway/2022-07-1363-1/2022|EPI ISL 18901632||A / H5N1
EPI3041161|PA|A/White-tailed eagle/Norway/2022-07-1997-1/2022|EPI ISL 18901644||A / H5N1
EPI3041177|PA|A/Northern gannet/Norway/2022-07-2112-1/2022|EPI ISL 18901646||A / H5N1
EPI3040904|PA|A/Northern gannet/Norway/2022-07-1073-1/2022|EPI ISL 18901609||A / H5N1
A/Red fox/Norway/2022-04-22642-2-4/2022
EPI3041113|PA|A/Great black-backed gull/Norway/2022-07-1594-1/2022|EPI ISL 18901638||A / H5N1
EPI3041098|PA|A/Northern gannet/Norway/2022-07-1583-1/2022|EPI ISL 18901636||A / H5N1
EPI3041169|PA|A/Northern gannet/Norway/2022-07-2085-1/2022|EPI ISL 18901645||A / H5N1
EPI2513406|PA|A/Red Fox/NL/FAV-0075/2023|EPI ISL 17494318||A / H5N1
EPI2513414|PA|A/American crow/PEI/FAV-0019-01/2022|EPI ISL 17494319||A / H5N1
EPI2198464|PA|A/chicken/Scotland/130966/2022|EPI ISL 15585888||A / H5N1
EPI3040958|PA|A/White-tailed eagle/Norway/2022-07-1118-1/2022|EPI ISL 18901616||A / H5N1
A/Red fox/Norway/2022-04-22642-1-4/2022
EPI3041298|PA|A/White-tailed eagle/Norway/2022-07-450-1/2022|EPI ISL 18901680||A / H5N1
EPI2008105|PA|A/chicken/England/012967/2022|EPI ISL 11561590||A / H5N1
EPI2072188|PA|A/magpie/Scotland/012805/2022|EPI ISL 13370895||A / H5N1
EPI3091610|PA|A/barnacle goose/Denmark/24273-1.02/2021|EPI ISL 18953100||A / H5N1
EPI3041229|PA|A/Rock dove/Norway/2022-07-3-1/2022|EPI ISL 18901671||A / H5N1
A/Eurasian Wigeon/Netherlands/1/2020
EPI3041314|PA|A/Raptor/Norway/2022-07-492-1/2022|EPI ISL 18901682||A / H5N1
EPI3041090|PA|A/White-tailed eagle/Norway/2022-07-157-1/2022|EPI ISL 18901635||A / H5N1
61
58
58
57
87
44
77
85
56
67
93
66
88
71
99
99
99
0.0020

D)

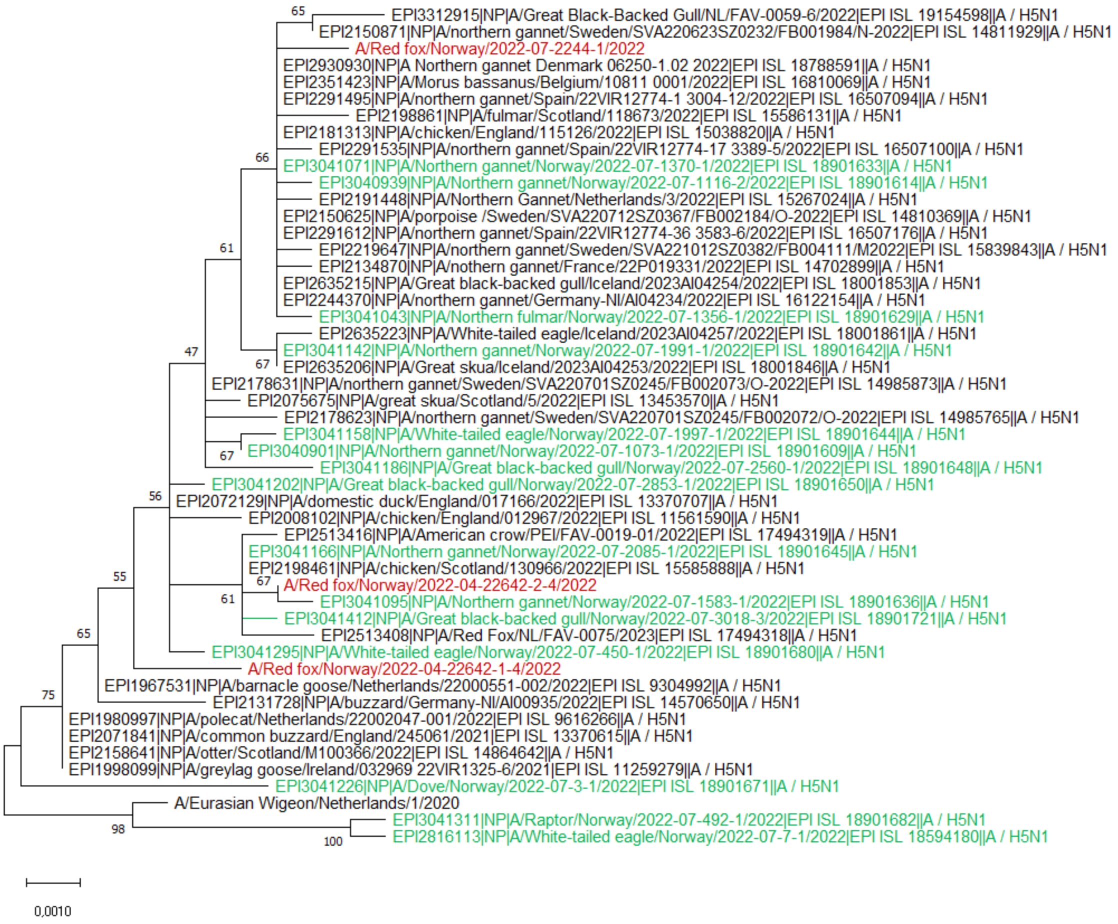

EPI3312915|NP|A/Great Black-Backed Gull/NL/FAV-0059-6/2022|EPI ISL 19154598||A / H5N1
EPI2150871|NP|A/northern gannet/Sweden/SVA220623SZ0232/FB001984/N-2022|EPI ISL 14811929||A / H5N1
A/Red fox/Norway/2022-07-2244-1/2022
EPI2930930|NP|A Northern gannet Denmark 06250-1.02 2022|EPI ISL 18788591||A / H5N1
EPI2351423|NP|A/Morus bassanus/Belgium/10811 0001/2022|EPI ISL 16810069||A / H5N1
EPI2291495|NP|A/northern gannet/Spain/22VIR12774-1 3004-12/2022|EPI ISL 16507094||A / H5N1
EPI2198861|NP|A/fulmar/Scotland/118673/2022|EPI ISL 15586131||A / H5N1
EPI2181313|NP|A/chicken/England/115126/2022|EPI ISL 15038820||A / H5N1
EPI2291535|NP|A/northern gannet/Spain/22VIR12774-17 3389-5/2022|EPI ISL 16507100||A / H5N1
EPI3041071|NP|A/Northern gannet/Norway/2022-07-1370-1/2022|EPI ISL 18901633||A / H5N1
EPI3040939|NP|A/Northern gannet/Norway/2022-07-1116-2/2022|EPI ISL 18901614||A / H5N1
EPI2191448|NP|A/Northern Gannet/Netherlands/3/2022|EPI ISL 15267024||A / H5N1
EPI2150625|NP|A/porpoise /Sweden/SVA220712SZ0367/FB002184/O-2022|EPI ISL 14810369||A / H5N1
EPI2291612|NP|A/northern gannet/Spain/22VIR12774-36 3583-6/2022|EPI ISL 16507176||A / H5N1
EPI2219647|NP|A/northern gannet/Sweden/SVA221012SZ0382/FB004111/M2022|EPI ISL 15839843||A / H5N1
EPI2134870|NP|A/nothern gannet/France/22P019331/2022|EPI ISL 14702899||A / H5N1
EPI2635215|NP|A/Great black-backed gull/Iceland/2023AI04254/2022|EPI ISL 18001853||A / H5N1
EPI2244370|NP|A/northern gannet/Germany-NI/AI04234/2022|EPI ISL 16122154||A / H5N1
EPI3041043|NP|A/Northern fulmar/Norway/2022-07-1356-1/2022|EPI ISL 18901629||A / H5N1
EPI2635223|NP|A/White-tailed eagle/Iceland/2023AI04257/2022|EPI ISL 18001861||A / H5N1
EPI3041142|NP|A/Northern gannet/Norway/2022-07-1991-1/2022|EPI ISL 18901642||A / H5N1
EPI2635206|NP|A/Great skua/Iceland/2023AI04253/2022|EPI ISL 18001846||A / H5N1
EPI2178631|NP|A/northern gannet/Sweden/SVA220701SZ0245/FB002073/O-2022|EPI ISL 14985873||A / H5N1
EPI2075675|NP|A/great skua/Scotland/5/2022|EPI ISL 13453570||A / H5N1
EPI2178623|NP|A/northern gannet/Sweden/SVA220701SZ0245/FB002072/O-2022|EPI ISL 14985765||A / H5N1
EPI3041158|NP|A/White-tailed eagle/Norway/2022-07-1997-1/2022|EPI ISL 18901644||A / H5N1
EPI3040901|NP|A/Northern gannet/Norway/2022-07-1073-1/2022|EPI ISL 18901609||A / H5N1
EPI3041186|NP|A/Great black-backed gull/Norway/2022-07-2560-1/2022|EPI ISL 18901648||A / H5N1
EPI3041202|NP|A/Great black-backed gull/Norway/2022-07-2853-1/2022|EPI ISL 18901650||A / H5N1
EPI2072129|NP|A/domestic duck/England/017166/2022|EPI ISL 13370707||A / H5N1
EPI2008102|NP|A/chicken/England/012967/2022|EPI ISL 11561590||A / H5N1
EPI2513416|NP|A/American crow/PEI/FAV-0019-01/2022|EPI ISL 17494319||A / H5N1
EPI3041166|NP|A/Northern gannet/Norway/2022-07-2085-1/2022|EPI ISL 18901645||A / H5N1
EPI2198461|NP|A/chicken/Scotland/130966/2022|EPI ISL 15585888||A / H5N1
A/Red fox/Norway/2022-04-22642-2-4/2022
EPI3041095|NP|A/Northern gannet/Norway/2022-07-1583-1/2022|EPI ISL 18901636||A / H5N1
EPI3041412|NP|A/Great black-backed gull/Norway/2022-07-3018-3/2022|EPI ISL 18901721||A / H5N1
EPI2513408|NP|A/Red Fox/NL/FAV-0075/2023|EPI ISL 17494318||A / H5N1
EPI3041295|NP|A/White-tailed eagle/Norway/2022-07-450-1/2022|EPI ISL 18901680||A / H5N1
A/Red fox/Norway/2022-04-22642-1-4/2022
EPI1967531|NP|A/barnacle goose/Netherlands/22000551-002/2022|EPI ISL 9304992||A / H5N1
EPI2131728|NP|A/buzzard/Germany-NI/AI00935/2022|EPI ISL 14570650||A / H5N1
EPI1980997|NP|A/polecat/Netherlands/22002047-001/2022|EPI ISL 9616266||A / H5N1
EPI2071841|NP|A/common buzzard/England/245061/2021|EPI ISL 13370615||A / H5N1
EPI2158641|NP|A/otter/Scotland/M100366/2022|EPI ISL 14864642||A / H5N1
EPI1998099|NP|A/greylag goose/Ireland/032969 22VIR1325-6/2021|EPI ISL 11259279||A / H5N1
EPI3041226|NP|A/Dove/Norway/2022-07-3-1/2022|EPI ISL 18901671||A / H5N1
A/Eurasian Wigeon/Netherlands/1/2020
EPI3041311|NP|A/Raptor/Norway/2022-07-492-1/2022|EPI ISL 18901682||A / H5N1
EPI2816113|NP|A/White-tailed eagle/Norway/2022-07-7-1/2022|EPI ISL 18594180||A / H5N1
0,0010

E)

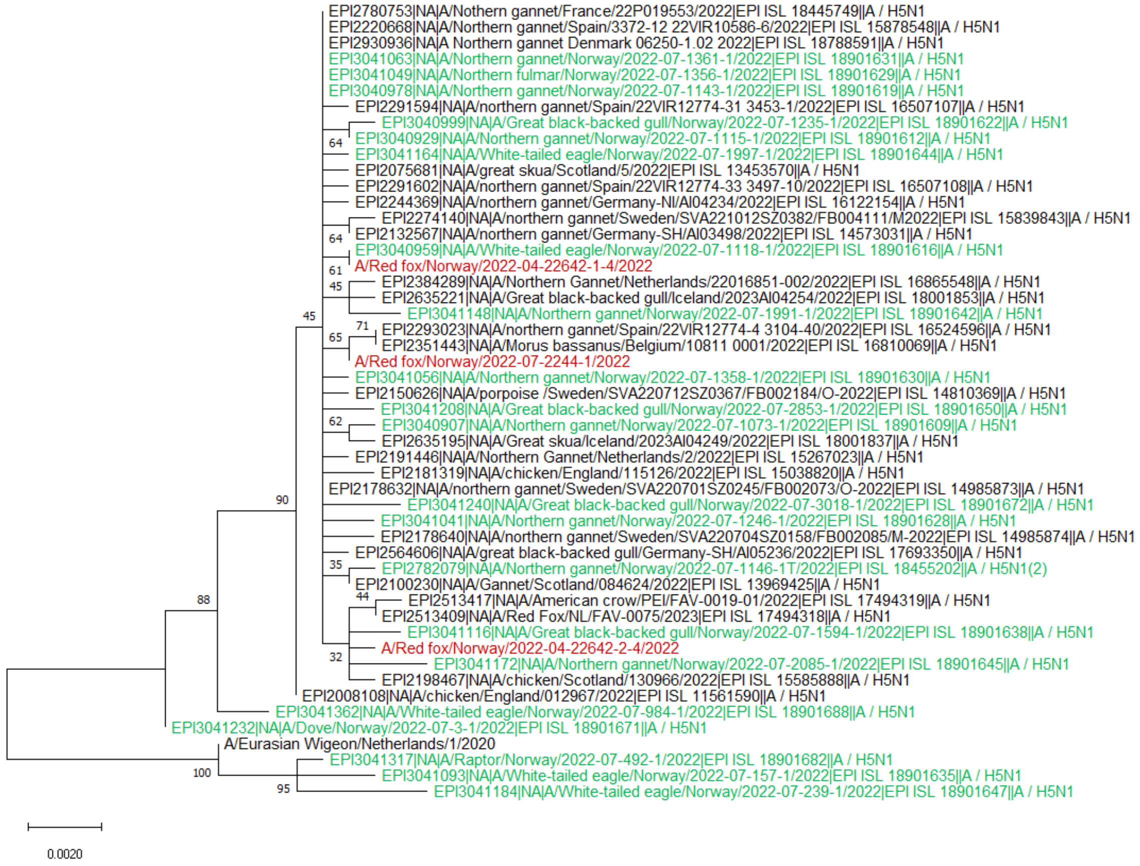

EPI2780753|NA|A/Nothern gannet/France/22P019553/2022|EPI ISL 18445749||A / H5N1
EPI2220668|NA|A/Northern gannet/Spain/3372-12 22VIR10586-6/2022|EPI ISL 15878548||A / H5N1
EPI2930936|NA|A Northern gannet Denmark 06250-1.02 2022|EPI ISL 18788591||A / H5N1
EPI3041063|NA|A/Northern gannet/Norway/2022-07-1361-1/2022|EPI ISL 18901631||A / H5N1
EPI3041049|NA|A/Northern fulmar/Norway/2022-07-1356-1/2022|EPI ISL 18901629||A / H5N1
EPI3040978|NA|A/Northern gannet/Norway/2022-07-1143-1/2022|EPI ISL 18901619||A / H5N1
EPI2291594|NA|A/northern gannet/Spain/22VIR12774-31 3453-1/2022|EPI ISL 16507107||A / H5N1
EPI3040999|NA|A/Great black-backed gull/Norway/2022-07-1235-1/2022|EPI ISL 18901622||A / H5N1
64
EPI3040929|NA|A/Northern gannet/Norway/2022-07-1115-1/2022|EPI ISL 18901612||A / H5N1
EPI3041164|NA|A/White-tailed eagle/Norway/2022-07-1997-1/2022|EPI ISL 18901644||A / H5N1
EPI2075681|NA|A/great skua/Scotland/5/2022|EPI ISL 13453570||A / H5N1
EPI2291602|NA|A/northern gannet/Spain/22VIR12774-33 3497-10/2022|EPI ISL 16507108||A / H5N1
EPI2244369|NA|A/northern gannet/Germany-NI/AI04234/2022|EPI ISL 16122154||A / H5N1
EPI2274140|NA|A/northern gannet/Sweden/SVA221012SZ0382/FB004111/M2022|EPI ISL 15839843||A / H5N1
64
EPI2132567|NA|A/northern gannet/Germany-SH/AI03498/2022|EPI ISL 14573031||A / H5N1
EPI3040959|NA|A/White-tailed eagle/Norway/2022-07-1118-1/2022|EPI ISL 18901616||A / H5N1
61
A/Red fox/Norway/2022-04-22642-1-4/2022
45
EPI2384289|NA|A/Northern Gannet/Netherlands/22016851-002/2022|EPI ISL 16865548||A / H5N1
EPI2635221|NA|A/Great black-backed gull/Iceland/2023AI04254/2022|EPI ISL 18001853||A / H5N1
45
EPI3041148|NA|A/Northern gannet/Norway/2022-07-1991-1/2022|EPI ISL 18901642||A / H5N1
71
EPI2293023|NA|A/northern gannet/Spain/22VIR12774-4 3104-40/2022|EPI ISL 16524596||A / H5N1
65
EPI2351443|NA|A/Morus bassanus/Belgium/10811 0001/2022|EPI ISL 16810069||A / H5N1
A/Red fox/Norway/2022-07-2244-1/2022
EPI3041056|NA|A/Northern gannet/Norway/2022-07-1358-1/2022|EPI ISL 18901630||A / H5N1
EPI2150626|NA|A/porpoise /Sweden/SVA220712SZ0367/FB002184/O-2022|EPI ISL 14810369||A / H5N1
EPI3041208|NA|A/Great black-backed gull/Norway/2022-07-2853-1/2022|EPI ISL 18901650||A / H5N1
62
EPI3040907|NA|A/Northern gannet/Norway/2022-07-1073-1/2022|EPI ISL 18901609||A / H5N1
EPI2635195|NA|A/Great skua/Iceland/2023AI04249/2022|EPI ISL 18001837||A / H5N1
EPI2191446|NA|A/Northern Gannet/Netherlands/2/2022|EPI ISL 15267023||A / H5N1
EPI2181319|NA|A/chicken/England/115126/2022|EPI ISL 15038820||A / H5N1
EPI2178632|NA|A/northern gannet/Sweden/SVA220701SZ0245/FB002073/O-2022|EPI ISL 14985873||A / H5N1
90
EPI3041240|NA|A/Great black-backed gull/Norway/2022-07-3018-1/2022|EPI ISL 18901672||A / H5N1
EPI3041041|NA|A/Northern gannet/Norway/2022-07-1246-1/2022|EPI ISL 18901628||A / H5N1
EPI2178640|NA|A/northern gannet/Sweden/SVA220704SZ0158/FB002085/M-2022|EPI ISL 14985874||A / H5N1
EPI2564606|NA|A/great black-backed gull/Germany-SH/AI05236/2022|EPI ISL 17693350||A / H5N1
35
EPI2782079|NA|A/Northern gannet/Norway/2022-07-1146-1T/2022|EPI ISL 18455202||A / H5N1(2)
EPI2100230|NA|A/Gannet/Scotland/084624/2022|EPI ISL 13969425||A / H5N1
88
44
EPI2513417|NA|A/American crow/PEI/FAV-0019-01/2022|EPI ISL 17494319||A / H5N1
EPI2513409|NA|A/Red Fox/NL/FAV-0075/2023|EPI ISL 17494318||A / H5N1
EPI3041116|NA|A/Great black-backed gull/Norway/2022-07-1594-1/2022|EPI ISL 18901638||A / H5N1
A/Red fox/Norway/2022-04-22642-2-4/2022
32
EPI3041172|NA|A/Northern gannet/Norway/2022-07-2085-1/2022|EPI ISL 18901645||A / H5N1
EPI2198467|NA|A/chicken/Scotland/130966/2022|EPI ISL 15585888||A / H5N1
EPI2008108|NA|A/chicken/England/012967/2022|EPI ISL 11561590||A / H5N1
EPI3041362|NA|A/White-tailed eagle/Norway/2022-07-984-1/2022|EPI ISL 18901688||A / H5N1
EPI3041232|NA|A/Dove/Norway/2022-07-3-1/2022|EPI ISL 18901671||A / H5N1
A/Eurasian Wigeon/Netherlands/1/2020
EPI3041317|NA|A/Raptor/Norway/2022-07-492-1/2022|EPI ISL 18901682||A / H5N1
100
EPI3041093|NA|A/White-tailed eagle/Norway/2022-07-157-1/2022|EPI ISL 18901635||A / H5N1
95
EPI3041184|NA|A/White-tailed eagle/Norway/2022-07-239-1/2022|EPI ISL 18901647||A / H5N1
0.0020

F)

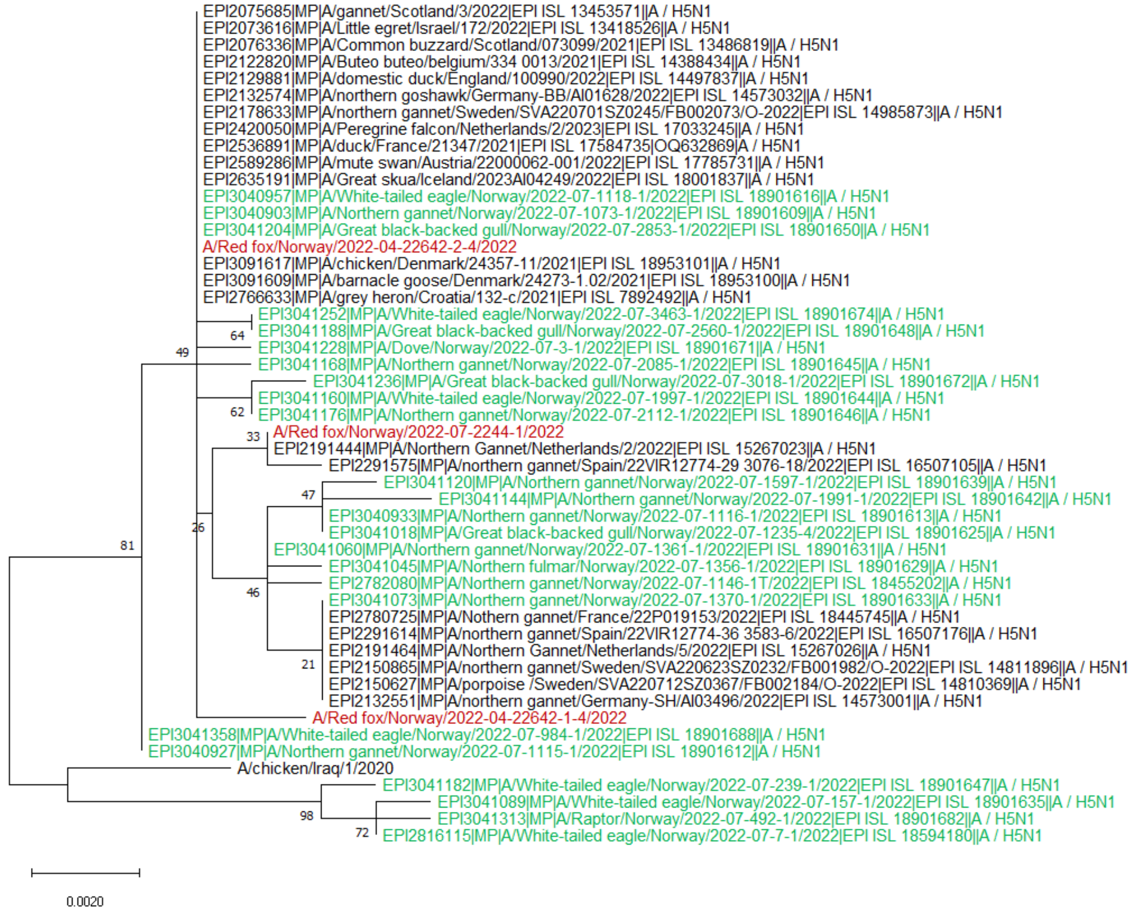
EPI2075685|MP|A/gannet/Scotland/3/2022|EPI ISL 13453571||A / H5N1
EPI2073616|MP|A/Little egret/Israel/172/2022|EPI ISL 13418526||A / H5N1
EPI2076336|MP|A/Common buzzard/Scotland/073099/2021|EPI ISL 13486819||A / H5N1
EPI2122820|MP|A/Buteo buteo/belgium/334 0013/2021|EPI ISL 14388434||A / H5N1
EPI2129881|MP|A/domestic duck/England/100990/2022|EPI ISL 14497837||A / H5N1
EPI2132574|MP|A/northern goshawk/Germany-BB/AI01628/2022|EPI ISL 14573032||A / H5N1
EPI2178633|MP|A/northern gannet/Sweden/SVA220701SZ0245/FB002073/O-2022|EPI ISL 14985873||A / H5N1
EPI2420050|MP|A/Peregrine falcon/Netherlands/2/2023|EPI ISL 17033245||A / H5N1
EPI2536891|MP|A/duck/France/21347/2021|EPI ISL 17584735|OQ632869|A / H5N1
EPI2589286|MP|A/mute swan/Austria/22000062-001/2022|EPI ISL 17785731||A / H5N1
EPI2635191|MP|A/Great skua/Iceland/2023AI04249/2022|EPI ISL 18001837||A / H5N1
EPI3040957|MP|A/White-tailed eagle/Norway/2022-07-1118-1/2022|EPI ISL 18901616||A / H5N1
EPI3040903|MP|A/Northern gannet/Norway/2022-07-1073-1/2022|EPI ISL 18901609||A / H5N1
EPI3041204|MP|A/Great black-backed gull/Norway/2022-07-2853-1/2022|EPI ISL 18901650||A / H5N1
A/Red fox/Norway/2022-04-22642-2-4/2022
EPI3091617|MP|A/chicken/Denmark/24357-11/2021|EPI ISL 18953101||A / H5N1
EPI3091609|MP|A/barnacle goose/Denmark/24273-1.02/2021|EPI ISL 18953100||A / H5N1
EPI2766633|MP|A/grey heron/Croatia/132-c/2021|EPI ISL 7892492||A / H5N1
EPI3041252|MP|A/White-tailed eagle/Norway/2022-07-3463-1/2022|EPI ISL 18901674||A / H5N1
EPI3041188|MP|A/Great black-backed gull/Norway/2022-07-2560-1/2022|EPI ISL 18901648||A / H5N1
EPI3041228|MP|A/Dove/Norway/2022-07-3-1/2022|EPI ISL 18901671||A / H5N1
EPI3041168|MP|A/Northern gannet/Norway/2022-07-2085-1/2022|EPI ISL 18901645||A / H5N1
EPI3041236|MP|A/Great black-backed gull/Norway/2022-07-3018-1/2022|EPI ISL 18901672||A / H5N1
EPI3041160|MP|A/White-tailed eagle/Norway/2022-07-1997-1/2022|EPI ISL 18901644||A / H5N1
EPI3041176|MP|A/Northern gannet/Norway/2022-07-2112-1/2022|EPI ISL 18901646||A / H5N1
A/Red fox/Norway/2022-07-2244-1/2022
EPI2191444|MP|A/Northern Gannet/Netherlands/2/2022|EPI ISL 15267023||A / H5N1
EPI2291575|MP|A/northern gannet/Spain/22VIR12774-29 3076-18/2022|EPI ISL 16507105||A / H5N1
EPI3041120|MP|A/Northern gannet/Norway/2022-07-1597-1/2022|EPI ISL 18901639||A / H5N1
EPI3041144|MP|A/Northern gannet/Norway/2022-07-1991-1/2022|EPI ISL 18901642||A / H5N1
EPI3040933|MP|A/Northern gannet/Norway/2022-07-1116-1/2022|EPI ISL 18901613||A / H5N1
EPI3041018|MP|A/Great black-backed gull/Norway/2022-07-1235-4/2022|EPI ISL 18901625||A / H5N1
EPI3041060|MP|A/Northern gannet/Norway/2022-07-1361-1/2022|EPI ISL 18901631||A / H5N1
EPI3041045|MP|A/Northern fulmar/Norway/2022-07-1356-1/2022|EPI ISL 18901629||A / H5N1
EPI2782080|MP|A/Northern gannet/Norway/2022-07-1146-1T/2022|EPI ISL 18455202||A / H5N1
EPI3041073|MP|A/Northern gannet/Norway/2022-07-1370-1/2022|EPI ISL 18901633||A / H5N1
EPI2780725|MP|A/Nothern gannet/France/22P019153/2022|EPI ISL 18445745||A / H5N1
EPI2291614|MP|A/northern gannet/Spain/22VIR12774-36 3583-6/2022|EPI ISL 16507176||A / H5N1
EPI2191464|MP|A/Northern Gannet/Netherlands/5/2022|EPI ISL 15267026||A / H5N1
EPI2150865|MP|A/northern gannet/Sweden/SVA220623SZ0232/FB001982/O-2022|EPI ISL 14811896||A / H5N1
EPI2150627|MP|A/porpoise /Sweden/SVA220712SZ0367/FB002184/O-2022|EPI ISL 14810369||A / H5N1
EPI2132551|MP|A/northern gannet/Germany-SH/AI03496/2022|EPI ISL 14573001||A / H5N1
A/Red fox/Norway/2022-04-22642-1-4/2022
EPI3041358|MP|A/White-tailed eagle/Norway/2022-07-984-1/2022|EPI ISL 18901688||A / H5N1
EPI3040927|MP|A/Northern gannet/Norway/2022-07-1115-1/2022|EPI ISL 18901612||A / H5N1
A/chicken/Iraq/1/2020
EPI3041182|MP|A/White-tailed eagle/Norway/2022-07-239-1/2022|EPI ISL 18901647||A / H5N1
EPI3041089|MP|A/White-tailed eagle/Norway/2022-07-157-1/2022|EPI ISL 18901635||A / H5N1
EPI3041313|MP|A/Raptor/Norway/2022-07-492-1/2022|EPI ISL 18901682||A / H5N1
EPI2816115|MP|A/White-tailed eagle/Norway/2022-07-7-1/2022|EPI ISL 18594180||A / H5N1
64
49
62
33
47
26
81
46
21
98
72
0.0020

G)

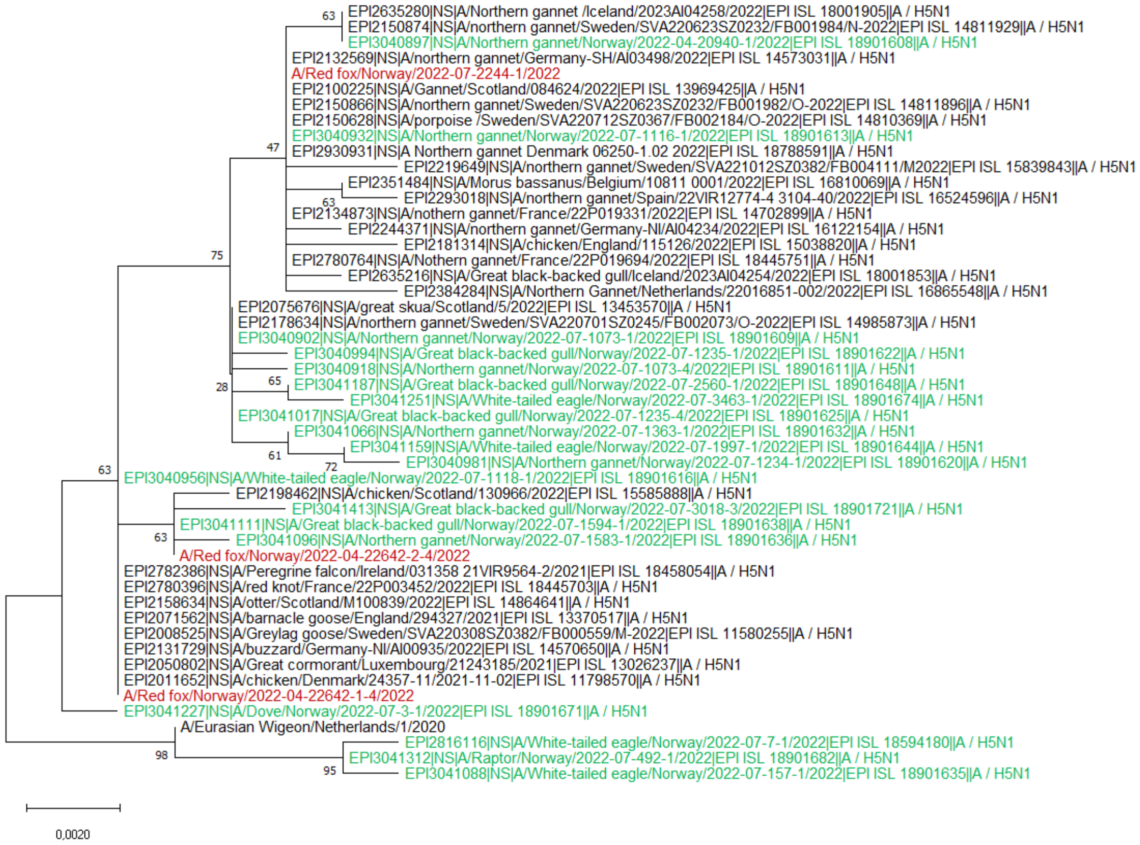

EPI2635280|NS|A/Northern gannet /Iceland/2023AI04258/2022|EPI ISL 18001905||A / H5N1
EPI2150874|NS|A/northern gannet/Sweden/SVA220623SZ0232/FB001984/N-2022|EPI ISL 14811929||A / H5N1
EPI3040897|NS|A/Northern gannet/Norway/2022-04-20940-1/2022|EPI ISL 18901608||A / H5N1
EPI2132569|NS|A/northern gannet/Germany-SH/AI03498/2022|EPI ISL 14573031||A / H5N1
A/Red fox/Norway/2022-07-2244-1/2022
EPI2100225|NS|A/Gannet/Scotland/084624/2022|EPI ISL 13969425||A / H5N1
EPI2150866|NS|A/northern gannet/Sweden/SVA220623SZ0232/FB001982/O-2022|EPI ISL 14811896||A / H5N1
EPI2150628|NS|A/porpoise /Sweden/SVA220712SZ0367/FB002184/O-2022|EPI ISL 14810369||A / H5N1
EPI3040932|NS|A/Northern gannet/Norway/2022-07-1116-1/2022|EPI ISL 18901613||A / H5N1
EPI2930931|NS|A Northern gannet Denmark 06250-1.02 2022|EPI ISL 18788591||A / H5N1
EPI2219649|NS|A/northern gannet/Sweden/SVA221012SZ0382/FB004111/M2022|EPI ISL 15839843||A / H5N1
EPI2351484|NS|A/Morus bassanus/Belgium/10811 0001/2022|EPI ISL 16810069||A / H5N1
EPI2293018|NS|A/northern gannet/Spain/22VIR12774-4 3104-40/2022|EPI ISL 16524596||A / H5N1
EPI2134873|NS|A/nothern gannet/France/22P019331/2022|EPI ISL 14702899||A / H5N1
EPI2244371|NS|A/northern gannet/Germany-NI/AI04234/2022|EPI ISL 16122154||A / H5N1
EPI2181314|NS|A/chicken/England/115126/2022|EPI ISL 15038820||A / H5N1
EPI2780764|NS|A/Nothern gannet/France/22P019694/2022|EPI ISL 18445751||A / H5N1
EPI2635216|NS|A/Great black-backed gull/Iceland/2023AI04254/2022|EPI ISL 18001853||A / H5N1
EPI2384284|NS|A/Northern Gannet/Netherlands/22016851-002/2022|EPI ISL 16865548||A / H5N1
EPI2075676|NS|A/great skua/Scotland/5/2022|EPI ISL 13453570||A / H5N1
EPI2178634|NS|A/northern gannet/Sweden/SVA220701SZ0245/FB002073/O-2022|EPI ISL 14985873||A / H5N1
EPI3040902|NS|A/Northern gannet/Norway/2022-07-1073-1/2022|EPI ISL 18901609||A / H5N1
EPI3040994|NS|A/Great black-backed gull/Norway/2022-07-1235-1/2022|EPI ISL 18901622||A / H5N1
EPI3040918|NS|A/Northern gannet/Norway/2022-07-1073-4/2022|EPI ISL 18901611||A / H5N1
EPI3041187|NS|A/Great black-backed gull/Norway/2022-07-2560-1/2022|EPI ISL 18901648||A / H5N1
EPI3041251|NS|A/White-tailed eagle/Norway/2022-07-3463-1/2022|EPI ISL 18901674||A / H5N1
EPI3041017|NS|A/Great black-backed gull/Norway/2022-07-1235-4/2022|EPI ISL 18901625||A / H5N1
EPI3041066|NS|A/Northern gannet/Norway/2022-07-1363-1/2022|EPI ISL 18901632||A / H5N1
EPI3041159|NS|A/White-tailed eagle/Norway/2022-07-1997-1/2022|EPI ISL 18901644||A / H5N1
EPI3040981|NS|A/Northern gannet/Norway/2022-07-1234-1/2022|EPI ISL 18901620||A / H5N1
EPI3040956|NS|A/White-tailed eagle/Norway/2022-07-1118-1/2022|EPI ISL 18901616||A / H5N1
EPI2198462|NS|A/chicken/Scotland/130966/2022|EPI ISL 15585888||A / H5N1
EPI3041413|NS|A/Great black-backed gull/Norway/2022-07-3018-3/2022|EPI ISL 18901721||A / H5N1
EPI3041111|NS|A/Great black-backed gull/Norway/2022-07-1594-1/2022|EPI ISL 18901638||A / H5N1
EPI3041096|NS|A/Northern gannet/Norway/2022-07-1583-1/2022|EPI ISL 18901636||A / H5N1
A/Red fox/Norway/2022-04-22642-2-4/2022
EPI2782386|NS|A/Peregrine falcon/Ireland/031358 21VIR9564-2/2021|EPI ISL 18458054||A / H5N1
EPI2780396|NS|A/red knot/France/22P003452/2022|EPI ISL 18445703||A / H5N1
EPI2158634|NS|A/otter/Scotland/M100839/2022|EPI ISL 14864641||A / H5N1
EPI2071562|NS|A/barnacle goose/England/294327/2021|EPI ISL 13370517||A / H5N1
EPI2008525|NS|A/Greylag goose/Sweden/SVA220308SZ0382/FB000559/M-2022|EPI ISL 11580255||A / H5N1
EPI2131729|NS|A/buzzard/Germany-NI/AI00935/2022|EPI ISL 14570650||A / H5N1
EPI2050802|NS|A/Great cormorant/Luxembourg/21243185/2021|EPI ISL 13026237||A / H5N1
EPI2011652|NS|A/chicken/Denmark/24357-11/2021-11-02|EPI ISL 11798570||A / H5N1
A/Red fox/Norway/2022-04-22642-1-4/2022
EPI3041227|NS|A/Dove/Norway/2022-07-3-1/2022|EPI ISL 18901671||A / H5N1
A/Eurasian Wigeon/Netherlands/1/2020
EPI2816116|NS|A/White-tailed eagle/Norway/2022-07-7-1/2022|EPI ISL 18594180||A / H5N1
EPI3041312|NS|A/Raptor/Norway/2022-07-492-1/2022|EPI ISL 18901682||A / H5N1
EPI3041088|NS|A/White-tailed eagle/Norway/2022-07-157-1/2022|EPI ISL 18901635||A / H5N1
63
47
63
75
28
65
61
72
63
63
98
95
0,0020

**Appendix Figure 4.** Midpoint rooted maximum likelihood phylogenetic trees showing the genetic relationship between highly pathogenic avian influenza H5N1 EA-2022-BB viruses from red foxes (red), wild birds (green) and other highly similar viruses obtained from GISAID (black). Trees are shown for the gene segments coding A) PB2, B) PB1, C) PA, D) NP, E) NA, F) M and G) NS.

A)

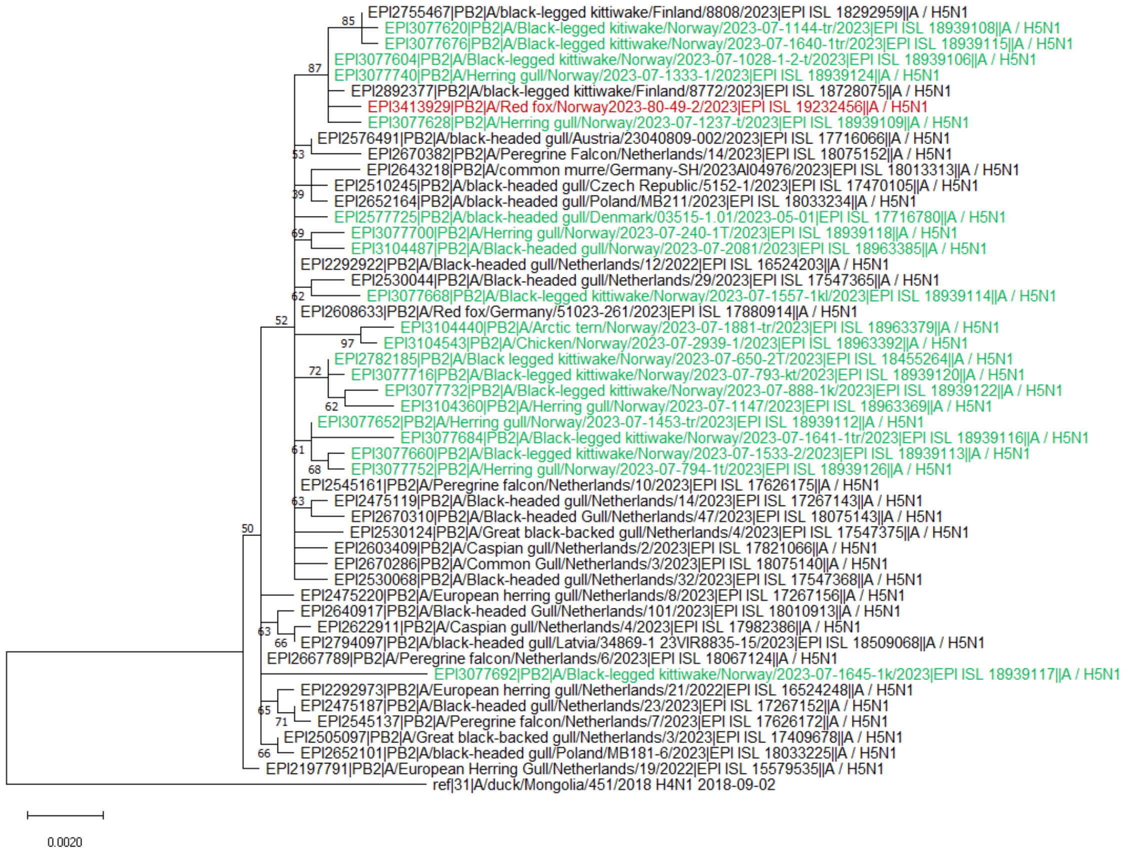

B)

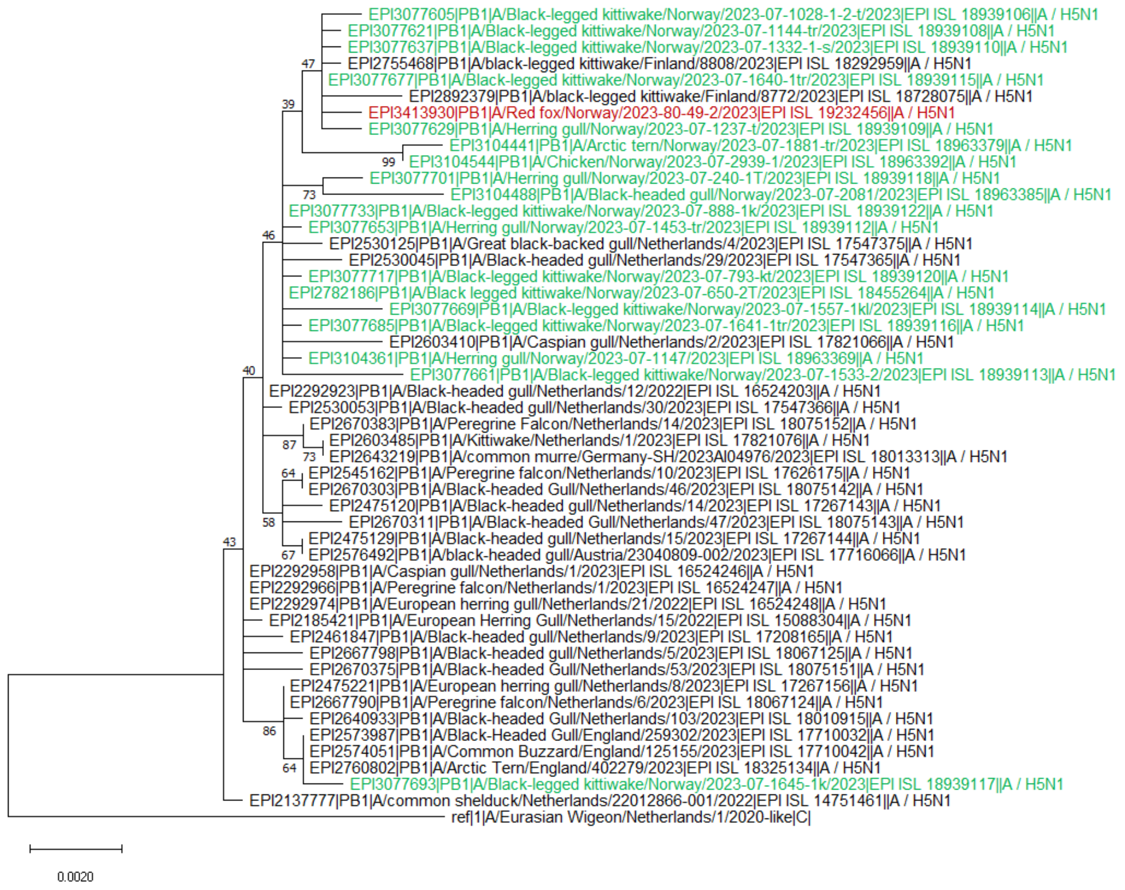
EPI3077605|PB1|A/Black-legged kittiwake/Norway/2023-07-1028-1-2-t/2023|EPI ISL 18939106||A / H5N1
EPI3077621|PB1|A/Black-legged kittiwake/Norway/2023-07-1144-tr/2023|EPI ISL 18939108||A / H5N1
EPI3077637|PB1|A/Black-legged kittiwake/Norway/2023-07-1332-1-s/2023|EPI ISL 18939110||A / H5N1
EPI2755468|PB1|A/black-legged kittiwake/Finland/8808/2023|EPI ISL 18292959||A / H5N1
EPI3077677|PB1|A/Black-legged kittiwake/Norway/2023-07-1640-1tr/2023|EPI ISL 18939115||A / H5N1
EPI2892379|PB1|A/black-legged kittiwake/Finland/8772/2023|EPI ISL 18728075||A / H5N1
EPI3413930|PB1|A/Red fox/Norway/2023-80-49-2/2023|EPI ISL 19232456||A / H5N1
EPI3077629|PB1|A/Herring gull/Norway/2023-07-1237-t/2023|EPI ISL 18939109||A / H5N1
EPI3104441|PB1|A/Arctic tern/Norway/2023-07-1881-tr/2023|EPI ISL 18963379||A / H5N1
EPI3104544|PB1|A/Chicken/Norway/2023-07-2939-1/2023|EPI ISL 18963392||A / H5N1
EPI3077701|PB1|A/Herring gull/Norway/2023-07-240-1T/2023|EPI ISL 18939118||A / H5N1
EPI3104488|PB1|A/Black-headed gull/Norway/2023-07-2081/2023|EPI ISL 18963385||A / H5N1
EPI3077733|PB1|A/Black-legged kittiwake/Norway/2023-07-888-1k/2023|EPI ISL 18939122||A / H5N1
EPI3077653|PB1|A/Herring gull/Norway/2023-07-1453-tr/2023|EPI ISL 18939112||A / H5N1
EPI2530125|PB1|A/Great black-backed gull/Netherlands/4/2023|EPI ISL 17547375||A / H5N1
EPI2530045|PB1|A/Black-headed gull/Netherlands/29/2023|EPI ISL 17547365||A / H5N1
EPI3077717|PB1|A/Black-legged kittiwake/Norway/2023-07-793-kt/2023|EPI ISL 18939120||A / H5N1
EPI2782186|PB1|A/Black legged kittiwake/Norway/2023-07-650-2T/2023|EPI ISL 18455264||A / H5N1
EPI3077669|PB1|A/Black-legged kittiwake/Norway/2023-07-1557-1kl/2023|EPI ISL 18939114||A / H5N1
EPI3077685|PB1|A/Black-legged kittiwake/Norway/2023-07-1641-1tr/2023|EPI ISL 18939116||A / H5N1
EPI2603410|PB1|A/Caspian gull/Netherlands/2/2023|EPI ISL 17821066||A / H5N1
EPI3104361|PB1|A/Herring gull/Norway/2023-07-1147/2023|EPI ISL 18963369||A / H5N1
EPI3077661|PB1|A/Black-legged kittiwake/Norway/2023-07-1533-2/2023|EPI ISL 18939113||A / H5N1
EPI2292923|PB1|A/Black-headed gull/Netherlands/12/2022|EPI ISL 16524203||A / H5N1
EPI2530053|PB1|A/Black-headed gull/Netherlands/30/2023|EPI ISL 17547366||A / H5N1
EPI2670383|PB1|A/Peregrine Falcon/Netherlands/14/2023|EPI ISL 18075152||A / H5N1
EPI2603485|PB1|A/Kittiwake/Netherlands/1/2023|EPI ISL 17821076||A / H5N1
EPI2643219|PB1|A/common murre/Germany-SH/2023AI04976/2023|EPI ISL 18013313||A / H5N1
EPI2545162|PB1|A/Peregrine falcon/Netherlands/10/2023|EPI ISL 17626175||A / H5N1
EPI2670303|PB1|A/Black-headed Gull/Netherlands/46/2023|EPI ISL 18075142||A / H5N1
EPI2475120|PB1|A/Black-headed gull/Netherlands/14/2023|EPI ISL 17267143||A / H5N1
EPI2670311|PB1|A/Black-headed Gull/Netherlands/47/2023|EPI ISL 18075143||A / H5N1
EPI2475129|PB1|A/Black-headed gull/Netherlands/15/2023|EPI ISL 17267144||A / H5N1
EPI2576492|PB1|A/black-headed gull/Austria/23040809-002/2023|EPI ISL 17716066||A / H5N1
EPI2292958|PB1|A/Caspian gull/Netherlands/1/2023|EPI ISL 16524246||A / H5N1
EPI2292966|PB1|A/Peregrine falcon/Netherlands/1/2023|EPI ISL 16524247||A / H5N1
EPI2292974|PB1|A/European herring gull/Netherlands/21/2022|EPI ISL 16524248||A / H5N1
EPI2185421|PB1|A/European Herring Gull/Netherlands/15/2022|EPI ISL 15088304||A / H5N1
EPI2461847|PB1|A/Black-headed gull/Netherlands/9/2023|EPI ISL 17208165||A / H5N1
EPI2667798|PB1|A/Black-headed gull/Netherlands/5/2023|EPI ISL 18067125||A / H5N1
EPI2670375|PB1|A/Black-headed Gull/Netherlands/53/2023|EPI ISL 18075151||A / H5N1
EPI2475221|PB1|A/European herring gull/Netherlands/8/2023|EPI ISL 17267156||A / H5N1
EPI2667790|PB1|A/Peregrine falcon/Netherlands/6/2023|EPI ISL 18067124||A / H5N1
EPI2640933|PB1|A/Black-headed Gull/Netherlands/103/2023|EPI ISL 18010915||A / H5N1
EPI2573987|PB1|A/Black-Headed Gull/England/259302/2023|EPI ISL 17710032||A / H5N1
EPI2574051|PB1|A/Common Buzzard/England/125155/2023|EPI ISL 17710042||A / H5N1
EPI2760802|PB1|A/Arctic Tern/England/402279/2023|EPI ISL 18325134||A / H5N1
EPI3077693|PB1|A/Black-legged kittiwake/Norway/2023-07-1645-1k/2023|EPI ISL 18939117||A / H5N1
EPI2137777|PB1|A/common shelduck/Netherlands/22012866-001/2022|EPI ISL 14751461||A / H5N1
ref|1|A/Eurasian Wigeon/Netherlands/1/2020-like|C|
0.0020

C)

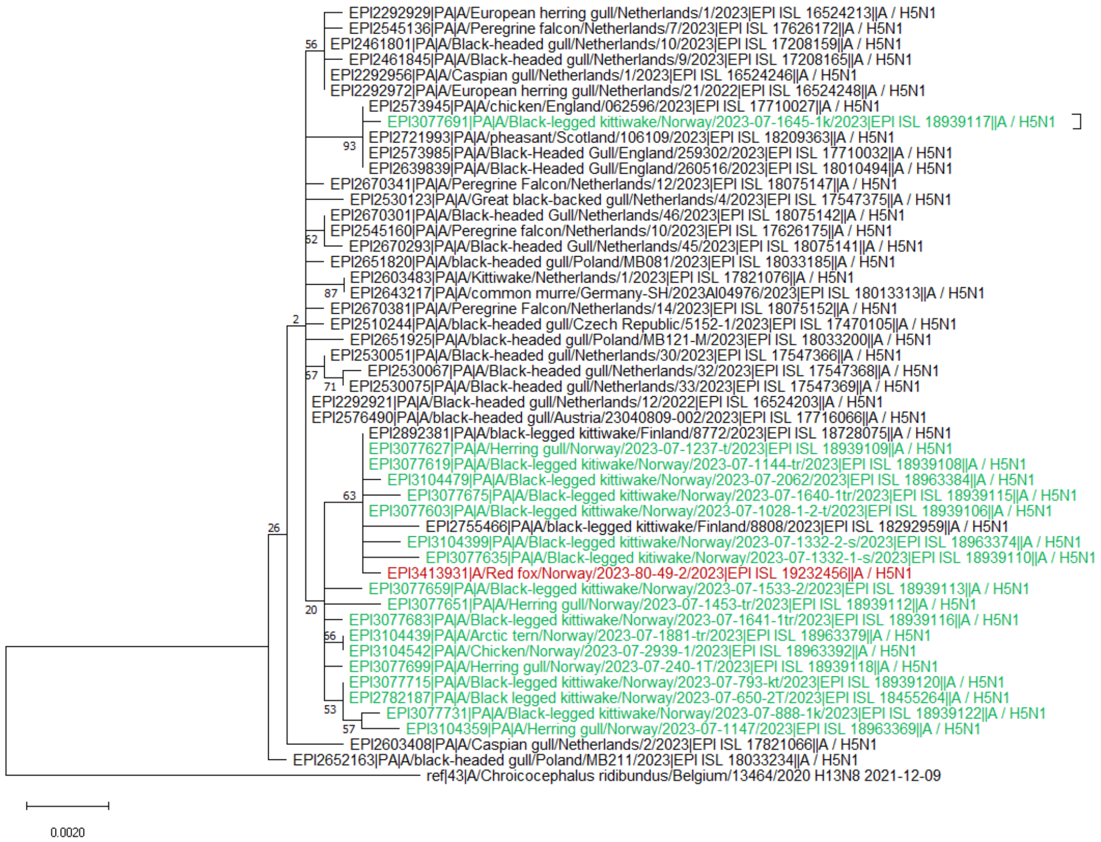
EPI2292929|PA|A/European herring gull/Netherlands/1/2023|EPI ISL 16524213||A / H5N1
EPI2545136|PA|A/Peregrine falcon/Netherlands/7/2023|EPI ISL 17626172||A / H5N1
56
EPI2461801|PA|A/Black-headed gull/Netherlands/10/2023|EPI ISL 17208159||A / H5N1
EPI2461845|PA|A/Black-headed gull/Netherlands/9/2023|EPI ISL 17208165||A / H5N1
EPI2292956|PA|A/Caspian gull/Netherlands/1/2023|EPI ISL 16524246||A / H5N1
EPI2292972|PA|A/European herring gull/Netherlands/21/2022|EPI ISL 16524248||A / H5N1
EPI2573945|PA|A/chicken/England/062596/2023|EPI ISL 17710027||A / H5N1
EPI3077691|PA|A/Black-legged kittiwake/Norway/2023-07-1645-1k/2023|EPI ISL 18939117||A / H5N1
EPI2721993|PA|A/pheasant/Scotland/106109/2023|EPI ISL 18209363||A / H5N1
93
EPI2573985|PA|A/Black-Headed Gull/England/259302/2023|EPI ISL 17710032||A / H5N1
EPI2639839|PA|A/Black-Headed Gull/England/260516/2023|EPI ISL 18010494||A / H5N1
EPI2670341|PA|A/Peregrine Falcon/Netherlands/12/2023|EPI ISL 18075147||A / H5N1
EPI2530123|PA|A/Great black-backed gull/Netherlands/4/2023|EPI ISL 17547375||A / H5N1
EPI2670301|PA|A/Black-headed Gull/Netherlands/46/2023|EPI ISL 18075142||A / H5N1
EPI2545160|PA|A/Peregrine falcon/Netherlands/10/2023|EPI ISL 17626175||A / H5N1
62
EPI2670293|PA|A/Black-headed Gull/Netherlands/45/2023|EPI ISL 18075141||A / H5N1
EPI2651820|PA|A/black-headed gull/Poland/MB081/2023|EPI ISL 18033185||A / H5N1
EPI2603483|PA|A/Kittiwake/Netherlands/1/2023|EPI ISL 17821076||A / H5N1
87
EPI2643217|PA|A/common murre/Germany-SH/2023AI04976/2023|EPI ISL 18013313||A / H5N1
EPI2670381|PA|A/Peregrine Falcon/Netherlands/14/2023|EPI ISL 18075152||A / H5N1
2
EPI2510244|PA|A/black-headed gull/Czech Republic/5152-1/2023|EPI ISL 17470105||A / H5N1
EPI2651925|PA|A/black-headed gull/Poland/MB121-M/2023|EPI ISL 18033200||A / H5N1
EPI2530051|PA|A/Black-headed gull/Netherlands/30/2023|EPI ISL 17547366||A / H5N1
EPI2530067|PA|A/Black-headed gull/Netherlands/32/2023|EPI ISL 17547368||A / H5N1
67
71
EPI2530075|PA|A/Black-headed gull/Netherlands/33/2023|EPI ISL 17547369||A / H5N1
EPI2292921|PA|A/Black-headed gull/Netherlands/12/2022|EPI ISL 16524203||A / H5N1
EPI2576490|PA|A/black-headed gull/Austria/23040809-002/2023|EPI ISL 17716066||A / H5N1
EPI2892381|PA|A/black-legged kittiwake/Finland/8772/2023|EPI ISL 18728075||A / H5N1
EPI3077627|PA|A/Herring gull/Norway/2023-07-1237-t/2023|EPI ISL 18939109||A / H5N1
EPI3077619|PA|A/Black-legged kitiwake/Norway/2023-07-1144-tr/2023|EPI ISL 18939108||A / H5N1
EPI3104479|PA|A/Black-legged kittiwake/Norway/2023-07-2062/2023|EPI ISL 18963384||A / H5N1
63
EPI3077675|PA|A/Black-legged kittiwake/Norway/2023-07-1640-1tr/2023|EPI ISL 18939115||A / H5N1
EPI3077603|PA|A/Black-legged kittiwake/Norway/2023-07-1028-1-2-t/2023|EPI ISL 18939106||A / H5N1
26
EPI2755466|PA|A/black-legged kittiwake/Finland/8808/2023|EPI ISL 18292959||A / H5N1
EPI3104399|PA|A/Black-legged kittiwake/Norway/2023-07-1332-2-s/2023|EPI ISL 18963374||A / H5N1
EPI3077635|PA|A/Black-legged kitiwake/Norway/2023-07-1332-1-s/2023|EPI ISL 18939110||A / H5N1
EPI3413931|A/Red fox/Norway/2023-80-49-2/2023|EPI ISL 19232456||A / H5N1
EPI3077659|PA|A/Black-legged kittiwake/Norway/2023-07-1533-2/2023|EPI ISL 18939113||A / H5N1
EPI3077651|PA|A/Herring gull/Norway/2023-07-1453-tr/2023|EPI ISL 18939112||A / H5N1
20
EPI3077683|PA|A/Black-legged kittiwake/Norway/2023-07-1641-1tr/2023|EPI ISL 18939116||A / H5N1
66
EPI3104439|PA|A/Arctic tern/Norway/2023-07-1881-tr/2023|EPI ISL 18963379||A / H5N1
EPI3104542|PA|A/Chicken/Norway/2023-07-2939-1/2023|EPI ISL 18963392||A / H5N1
EPI3077699|PA|A/Herring gull/Norway/2023-07-240-1T/2023|EPI ISL 18939118||A / H5N1
EPI3077715|PA|A/Black-legged kittiwake/Norway/2023-07-793-kt/2023|EPI ISL 18939120||A / H5N1
EPI2782187|PA|A/Black legged kittiwake/Norway/2023-07-650-2T/2023|EPI ISL 18455264||A / H5N1
53
EPI3077731|PA|A/Black-legged kittiwake/Norway/2023-07-888-1k/2023|EPI ISL 18939122||A / H5N1
57
EPI3104359|PA|A/Herring gull/Norway/2023-07-1147/2023|EPI ISL 18963369||A / H5N1
EPI2603408|PA|A/Caspian gull/Netherlands/2/2023|EPI ISL 17821066||A / H5N1
EPI2652163|PA|A/black-headed gull/Poland/MB211/2023|EPI ISL 18033234||A / H5N1
ref|43|A/Chroicocephalus ridibundus/Belgium/13464/2020 H13N8 2021-12-09
0.0020

D)

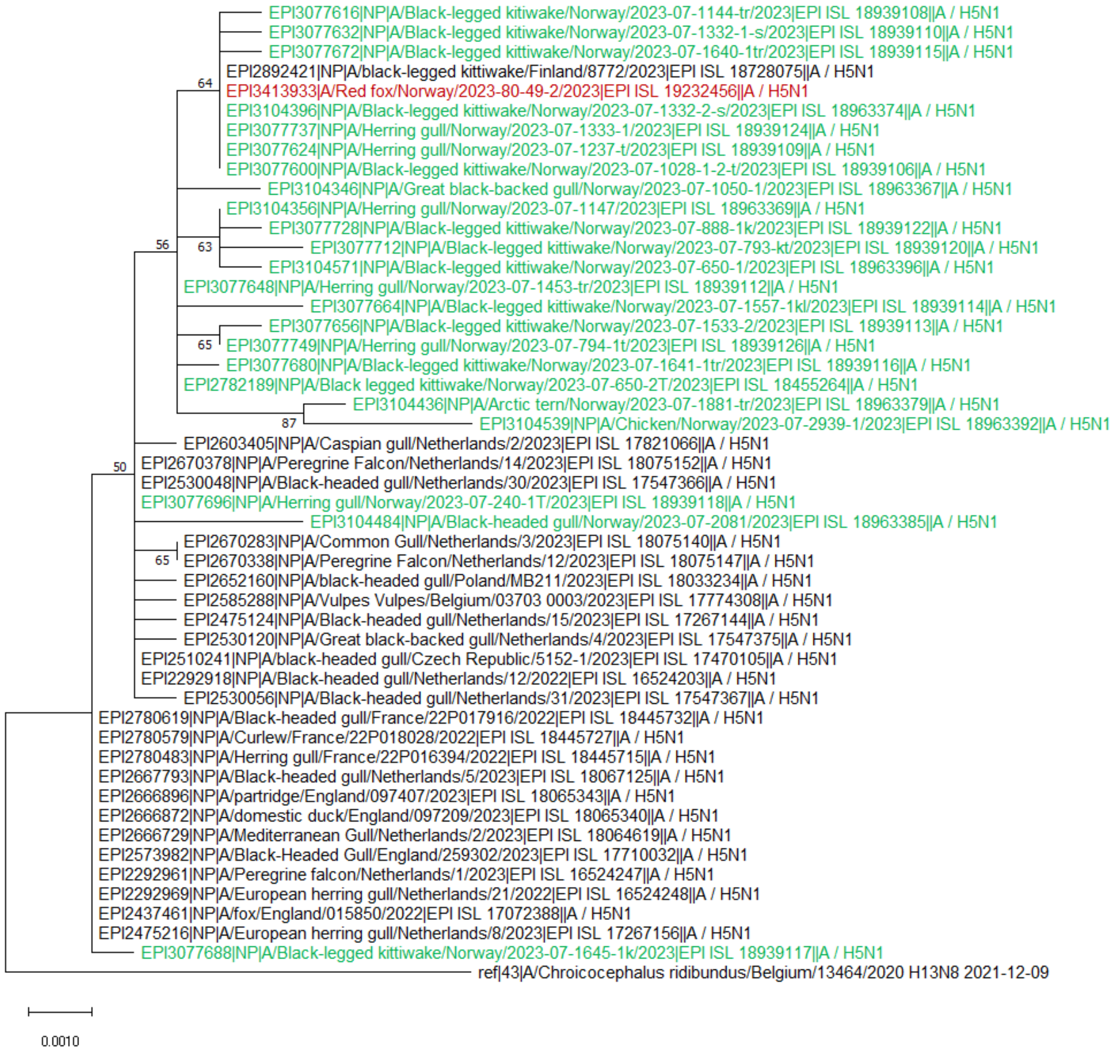

EPI3077616|NP|A/Black-legged kitiwake/Norway/2023-07-1144-tr/2023|EPI ISL 18939108||A / H5N1
EPI3077632|NP|A/Black-legged kitiwake/Norway/2023-07-1332-1-s/2023|EPI ISL 18939110||A / H5N1
EPI3077672|NP|A/Black-legged kittiwake/Norway/2023-07-1640-1tr/2023|EPI ISL 18939115||A / H5N1
64
EPI2892421|NP|A/black-legged kittiwake/Finland/8772/2023|EPI ISL 18728075||A / H5N1
EPI3413933|A/Red fox/Norway/2023-80-49-2/2023|EPI ISL 19232456||A / H5N1
EPI3104396|NP|A/Black-legged kittiwake/Norway/2023-07-1332-2-s/2023|EPI ISL 18963374||A / H5N1
EPI3077737|NP|A/Herring gull/Norway/2023-07-1333-1/2023|EPI ISL 18939124||A / H5N1
EPI3077624|NP|A/Herring gull/Norway/2023-07-1237-t/2023|EPI ISL 18939109||A / H5N1
EPI3077600|NP|A/Black-legged kittiwake/Norway/2023-07-1028-1-2-t/2023|EPI ISL 18939106||A / H5N1
EPI3104346|NP|A/Great black-backed gull/Norway/2023-07-1050-1/2023|EPI ISL 18963367||A / H5N1
EPI3104356|NP|A/Herring gull/Norway/2023-07-1147/2023|EPI ISL 18963369||A / H5N1
EPI3077728|NP|A/Black-legged kittiwake/Norway/2023-07-888-1k/2023|EPI ISL 18939122||A / H5N1
56
63
EPI3077712|NP|A/Black-legged kittiwake/Norway/2023-07-793-kt/2023|EPI ISL 18939120||A / H5N1
EPI3104571|NP|A/Black-legged kittiwake/Norway/2023-07-650-1/2023|EPI ISL 18963396||A / H5N1
EPI3077648|NP|A/Herring gull/Norway/2023-07-1453-tr/2023|EPI ISL 18939112||A / H5N1
EPI3077664|NP|A/Black-legged kittiwake/Norway/2023-07-1557-1kl/2023|EPI ISL 18939114||A / H5N1
EPI3077656|NP|A/Black-legged kittiwake/Norway/2023-07-1533-2/2023|EPI ISL 18939113||A / H5N1
65
EPI3077749|NP|A/Herring gull/Norway/2023-07-794-1t/2023|EPI ISL 18939126||A / H5N1
EPI3077680|NP|A/Black-legged kittiwake/Norway/2023-07-1641-1tr/2023|EPI ISL 18939116||A / H5N1
EPI2782189|NP|A/Black legged kittiwake/Norway/2023-07-650-2T/2023|EPI ISL 18455264||A / H5N1
EPI3104436|NP|A/Arctic tern/Norway/2023-07-1881-tr/2023|EPI ISL 18963379||A / H5N1
87
EPI3104539|NP|A/Chicken/Norway/2023-07-2939-1/2023|EPI ISL 18963392||A / H5N1
EPI2603405|NP|A/Caspian gull/Netherlands/2/2023|EPI ISL 17821066||A / H5N1
50
EPI2670378|NP|A/Peregrine Falcon/Netherlands/14/2023|EPI ISL 18075152||A / H5N1
EPI2530048|NP|A/Black-headed gull/Netherlands/30/2023|EPI ISL 17547366||A / H5N1
EPI3077696|NP|A/Herring gull/Norway/2023-07-240-1T/2023|EPI ISL 18939118||A / H5N1
EPI3104484|NP|A/Black-headed gull/Norway/2023-07-2081/2023|EPI ISL 18963385||A / H5N1
EPI2670283|NP|A/Common Gull/Netherlands/3/2023|EPI ISL 18075140||A / H5N1
65
EPI2670338|NP|A/Peregrine Falcon/Netherlands/12/2023|EPI ISL 18075147||A / H5N1
EPI2652160|NP|A/black-headed gull/Poland/MB211/2023|EPI ISL 18033234||A / H5N1
EPI2585288|NP|A/Vulpes Vulpes/Belgium/03703 0003/2023|EPI ISL 17774308||A / H5N1
EPI2475124|NP|A/Black-headed gull/Netherlands/15/2023|EPI ISL 17267144||A / H5N1
EPI2530120|NP|A/Great black-backed gull/Netherlands/4/2023|EPI ISL 17547375||A / H5N1
EPI2510241|NP|A/black-headed gull/Czech Republic/5152-1/2023|EPI ISL 17470105||A / H5N1
EPI2292918|NP|A/Black-headed gull/Netherlands/12/2022|EPI ISL 16524203||A / H5N1
EPI2530056|NP|A/Black-headed gull/Netherlands/31/2023|EPI ISL 17547367||A / H5N1
EPI2780619|NP|A/Black-headed gull/France/22P017916/2022|EPI ISL 18445732||A / H5N1
EPI2780579|NP|A/Curlew/France/22P018028/2022|EPI ISL 18445727||A / H5N1
EPI2780483|NP|A/Herring gull/France/22P016394/2022|EPI ISL 18445715||A / H5N1
EPI2667793|NP|A/Black-headed gull/Netherlands/5/2023|EPI ISL 18067125||A / H5N1
EPI2666896|NP|A/partridge/England/097407/2023|EPI ISL 18065343||A / H5N1
EPI2666872|NP|A/domestic duck/England/097209/2023|EPI ISL 18065340||A / H5N1
EPI2666729|NP|A/Mediterranean Gull/Netherlands/2/2023|EPI ISL 18064619||A / H5N1
EPI2573982|NP|A/Black-Headed Gull/England/259302/2023|EPI ISL 17710032||A / H5N1
EPI2292961|NP|A/Peregrine falcon/Netherlands/1/2023|EPI ISL 16524247||A / H5N1
EPI2292969|NP|A/European herring gull/Netherlands/21/2022|EPI ISL 16524248||A / H5N1
EPI2437461|NP|A/fox/England/015850/2022|EPI ISL 17072388||A / H5N1
EPI2475216|NP|A/European herring gull/Netherlands/8/2023|EPI ISL 17267156||A / H5N1
EPI3077688|NP|A/Black-legged kittiwake/Norway/2023-07-1645-1k/2023|EPI ISL 18939117||A / H5N1
ref|43|A/Chroicocephalus ridibundus/Belgium/13464/2020 H13N8 2021-12-09
0.0010

E)

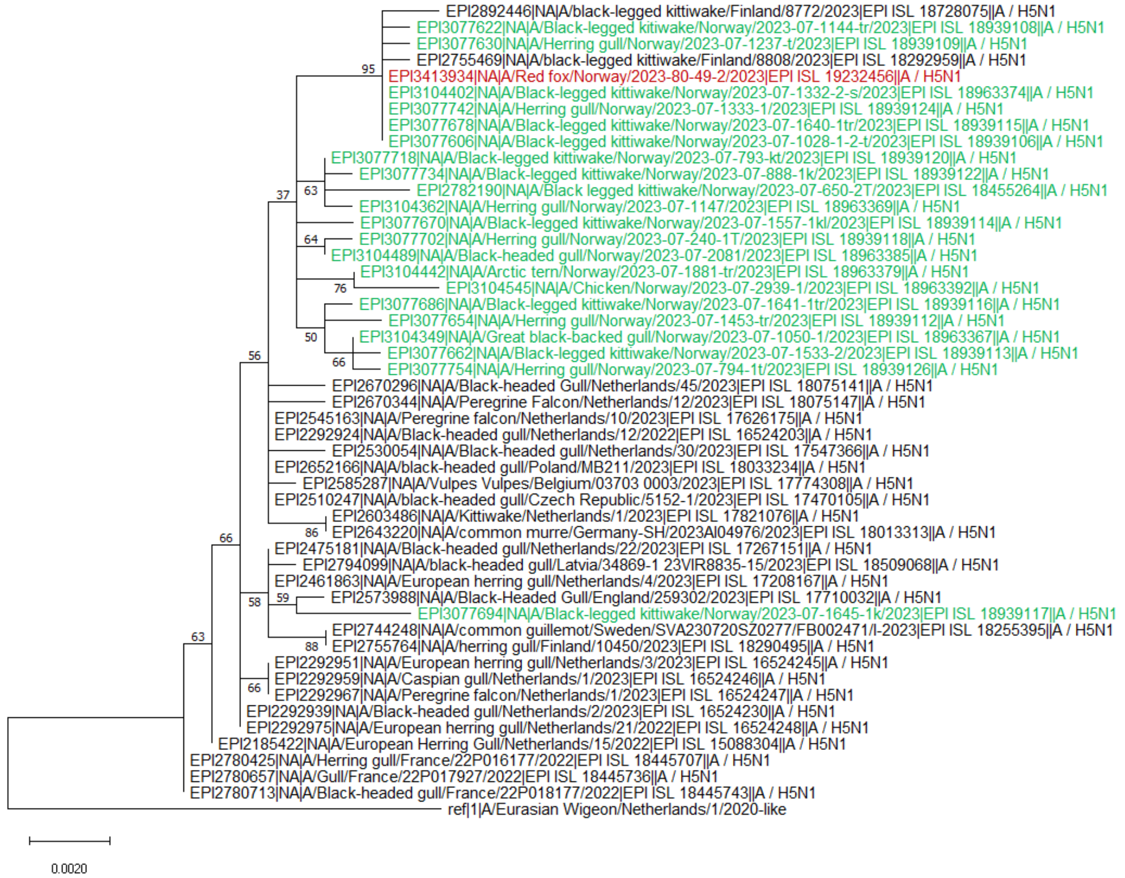

EPI2892446|NA|A/black-legged kittiwake/Finland/8772/2023|EPI ISL 18728075||A / H5N1
EPI3077622|NA|A/Black-legged kitiwake/Norway/2023-07-1144-tr/2023|EPI ISL 18939108||A / H5N1
EPI3077630|NA|A/Herring gull/Norway/2023-07-1237-t/2023|EPI ISL 18939109||A / H5N1
EPI2755469|NA|A/black-legged kittiwake/Finland/8808/2023|EPI ISL 18292959||A / H5N1
EPI3413934|NA|A/Red fox/Norway/2023-80-49-2/2023|EPI ISL 19232456||A / H5N1
EPI3104402|NA|A/Black-legged kittiwake/Norway/2023-07-1332-2-s/2023|EPI ISL 18963374||A / H5N1
EPI3077742|NA|A/Herring gull/Norway/2023-07-1333-1/2023|EPI ISL 18939124||A / H5N1
EPI3077678|NA|A/Black-legged kittiwake/Norway/2023-07-1640-1tr/2023|EPI ISL 18939115||A / H5N1
EPI3077606|NA|A/Black-legged kittiwake/Norway/2023-07-1028-1-2-t/2023|EPI ISL 18939106||A / H5N1
EPI3077718|NA|A/Black-legged kittiwake/Norway/2023-07-793-kt/2023|EPI ISL 18939120||A / H5N1
EPI3077734|NA|A/Black-legged kittiwake/Norway/2023-07-888-1k/2023|EPI ISL 18939122||A / H5N1
EPI2782190|NA|A/Black legged kittiwake/Norway/2023-07-650-2T/2023|EPI ISL 18455264||A / H5N1
EPI3104362|NA|A/Herring gull/Norway/2023-07-1147/2023|EPI ISL 18963369||A / H5N1
EPI3077670|NA|A/Black-legged kittiwake/Norway/2023-07-1557-1kl/2023|EPI ISL 18939114||A / H5N1
EPI3077702|NA|A/Herring gull/Norway/2023-07-240-1T/2023|EPI ISL 18939118||A / H5N1
EPI3104489|NA|A/Black-headed gull/Norway/2023-07-2081/2023|EPI ISL 18963385||A / H5N1
EPI3104442|NA|A/Arctic tern/Norway/2023-07-1881-tr/2023|EPI ISL 18963379||A / H5N1
EPI3104545|NA|A/Chicken/Norway/2023-07-2939-1/2023|EPI ISL 18963392||A / H5N1
EPI3077686|NA|A/Black-legged kittiwake/Norway/2023-07-1641-1tr/2023|EPI ISL 18939116||A / H5N1
EPI3077654|NA|A/Herring gull/Norway/2023-07-1453-tr/2023|EPI ISL 18939112||A / H5N1
EPI3104349|NA|A/Great black-backed gull/Norway/2023-07-1050-1/2023|EPI ISL 18963367||A / H5N1
EPI3077662|NA|A/Black-legged kittiwake/Norway/2023-07-1533-2/2023|EPI ISL 18939113||A / H5N1
EPI3077754|NA|A/Herring gull/Norway/2023-07-794-1t/2023|EPI ISL 18939126||A / H5N1
EPI2670296|NA|A/Black-headed Gull/Netherlands/45/2023|EPI ISL 18075141||A / H5N1
EPI2670344|NA|A/Peregrine Falcon/Netherlands/12/2023|EPI ISL 18075147||A / H5N1
EPI2545163|NA|A/Peregrine falcon/Netherlands/10/2023|EPI ISL 17626175||A / H5N1
EPI2292924|NA|A/Black-headed gull/Netherlands/12/2022|EPI ISL 16524203||A / H5N1
EPI2530054|NA|A/Black-headed gull/Netherlands/30/2023|EPI ISL 17547366||A / H5N1
EPI2652166|NA|A/black-headed gull/Poland/MB211/2023|EPI ISL 18033234||A / H5N1
EPI2585287|NA|A/Vulpes Vulpes/Belgium/03703 0003/2023|EPI ISL 17774308||A / H5N1
EPI2510247|NA|A/black-headed gull/Czech Republic/5152-1/2023|EPI ISL 17470105||A / H5N1
EPI2603486|NA|A/Kittiwake/Netherlands/1/2023|EPI ISL 17821076||A / H5N1
EPI2643220|NA|A/common murre/Germany-SH/2023AI04976/2023|EPI ISL 18013313||A / H5N1
EPI2475181|NA|A/Black-headed gull/Netherlands/22/2023|EPI ISL 17267151||A / H5N1
EPI2794099|NA|A/black-headed gull/Latvia/34869-1 23VIR8835-15/2023|EPI ISL 18509068||A / H5N1
EPI2461863|NA|A/European herring gull/Netherlands/4/2023|EPI ISL 17208167||A / H5N1
EPI2573988|NA|A/Black-Headed Gull/England/259302/2023|EPI ISL 17710032||A / H5N1
EPI3077694|NA|A/Black-legged kittiwake/Norway/2023-07-1645-1k/2023|EPI ISL 18939117||A / H5N1
EPI2744248|NA|A/common guillemot/Sweden/SVA230720SZ0277/FB002471/I-2023|EPI ISL 18255395||A / H5N1
EPI2755764|NA|A/herring gull/Finland/10450/2023|EPI ISL 18290495||A / H5N1
EPI2292951|NA|A/European herring gull/Netherlands/3/2023|EPI ISL 16524245||A / H5N1
EPI2292959|NA|A/Caspian gull/Netherlands/1/2023|EPI ISL 16524246||A / H5N1
EPI2292967|NA|A/Peregrine falcon/Netherlands/1/2023|EPI ISL 16524247||A / H5N1
EPI2292939|NA|A/Black-headed gull/Netherlands/2/2023|EPI ISL 16524230||A / H5N1
EPI2292975|NA|A/European herring gull/Netherlands/21/2022|EPI ISL 16524248||A / H5N1
EPI2185422|NA|A/European Herring Gull/Netherlands/15/2022|EPI ISL 15088304||A / H5N1
EPI2780425|NA|A/Herring gull/France/22P016177/2022|EPI ISL 18445707||A / H5N1
EPI2780657|NA|A/Gull/France/22P017927/2022|EPI ISL 18445736||A / H5N1
EPI2780713|NA|A/Black-headed gull/France/22P018177/2022|EPI ISL 18445743||A / H5N1
ref|1|A/Eurasian Wigeon/Netherlands/1/2020-like
95
63
37
64
76
50
66
56
86
66
58
59
88
66
63
0.0020

F)

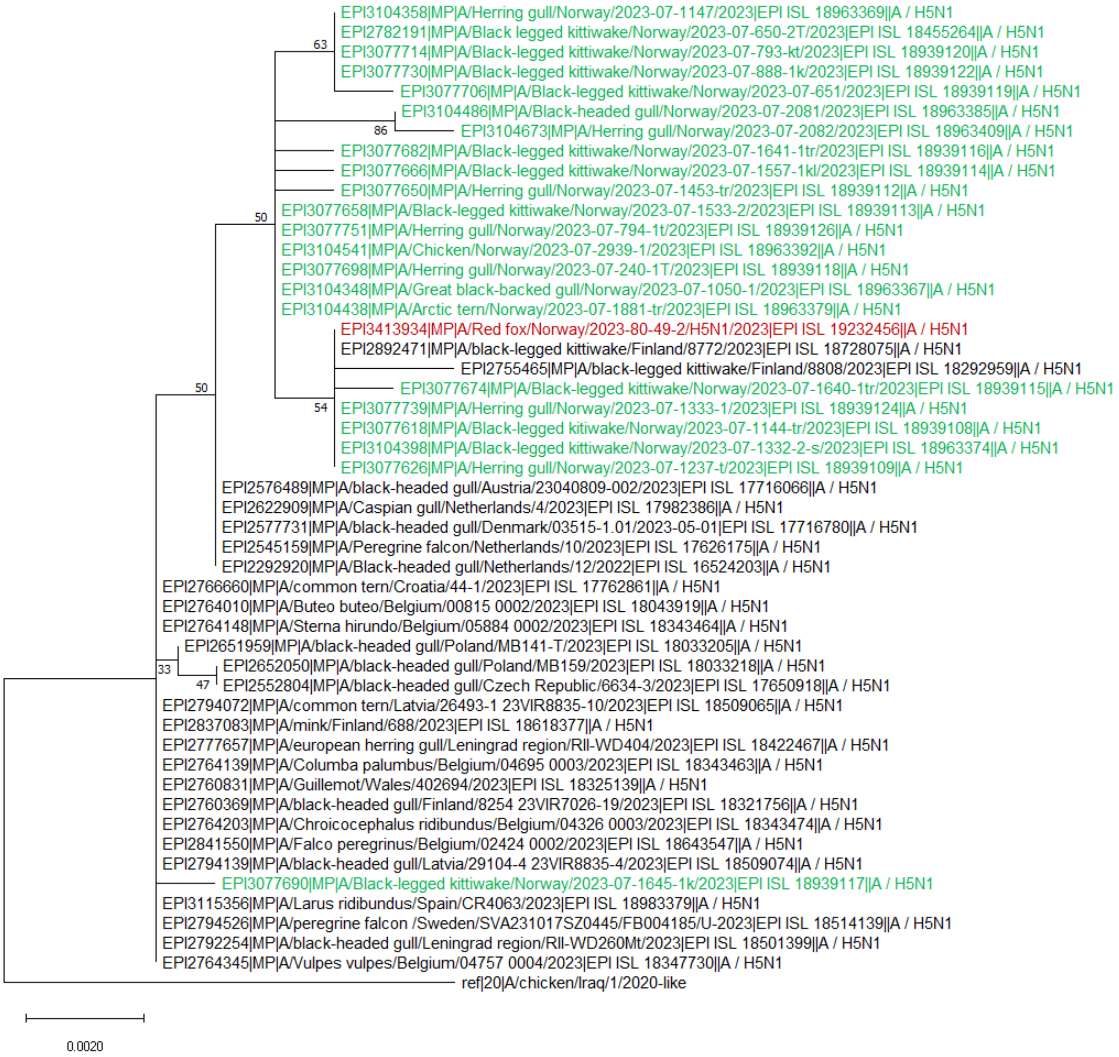
EPI3104358|MP|A/Herring gull/Norway/2023-07-1147/2023|EPI ISL 18963369||A / H5N1
EPI2782191|MP|A/Black legged kittiwake/Norway/2023-07-650-2T/2023|EPI ISL 18455264||A / H5N1
63
EPI3077714|MP|A/Black-legged kittiwake/Norway/2023-07-793-kt/2023|EPI ISL 18939120||A / H5N1
EPI3077730|MP|A/Black-legged kittiwake/Norway/2023-07-888-1k/2023|EPI ISL 18939122||A / H5N1
EPI3077706|MP|A/Black-legged kittiwake/Norway/2023-07-651/2023|EPI ISL 18939119||A / H5N1
EPI3104486|MP|A/Black-headed gull/Norway/2023-07-2081/2023|EPI ISL 18963385||A / H5N1
86
EPI3104673|MP|A/Herring gull/Norway/2023-07-2082/2023|EPI ISL 18963409||A / H5N1
EPI3077682|MP|A/Black-legged kittiwake/Norway/2023-07-1641-1tr/2023|EPI ISL 18939116||A / H5N1
EPI3077666|MP|A/Black-legged kittiwake/Norway/2023-07-1557-1kl/2023|EPI ISL 18939114||A / H5N1
EPI3077650|MP|A/Herring gull/Norway/2023-07-1453-tr/2023|EPI ISL 18939112||A / H5N1
50
EPI3077658|MP|A/Black-legged kittiwake/Norway/2023-07-1533-2/2023|EPI ISL 18939113||A / H5N1
EPI3077751|MP|A/Herring gull/Norway/2023-07-794-1t/2023|EPI ISL 18939126||A / H5N1
EPI3104541|MP|A/Chicken/Norway/2023-07-2939-1/2023|EPI ISL 18963392||A / H5N1
EPI3077698|MP|A/Herring gull/Norway/2023-07-240-1T/2023|EPI ISL 18939118||A / H5N1
EPI3104348|MP|A/Great black-backed gull/Norway/2023-07-1050-1/2023|EPI ISL 18963367||A / H5N1
EPI3104438|MP|A/Arctic tern/Norway/2023-07-1881-tr/2023|EPI ISL 18963379||A / H5N1
EPI3413934|MP|A/Red fox/Norway/2023-80-49-2/H5N1/2023|EPI ISL 19232456||A / H5N1
EPI2892471|MP|A/black-legged kittiwake/Finland/8772/2023|EPI ISL 18728075||A / H5N1
EPI2755465|MP|A/black-legged kittiwake/Finland/8808/2023|EPI ISL 18292959||A / H5N1
50
EPI3077674|MP|A/Black-legged kittiwake/Norway/2023-07-1640-1tr/2023|EPI ISL 18939115||A / H5N1
54
EPI3077739|MP|A/Herring gull/Norway/2023-07-1333-1/2023|EPI ISL 18939124||A / H5N1
EPI3077618|MP|A/Black-legged kitiwake/Norway/2023-07-1144-tr/2023|EPI ISL 18939108||A / H5N1
EPI3104398|MP|A/Black-legged kittiwake/Norway/2023-07-1332-2-s/2023|EPI ISL 18963374||A / H5N1
EPI3077626|MP|A/Herring gull/Norway/2023-07-1237-t/2023|EPI ISL 18939109||A / H5N1
EPI2576489|MP|A/black-headed gull/Austria/23040809-002/2023|EPI ISL 17716066||A / H5N1
EPI2622909|MP|A/Caspian gull/Netherlands/4/2023|EPI ISL 17982386||A / H5N1
EPI2577731|MP|A/black-headed gull/Denmark/03515-1.01/2023-05-01|EPI ISL 17716780||A / H5N1
EPI2545159|MP|A/Peregrine falcon/Netherlands/10/2023|EPI ISL 17626175||A / H5N1
EPI2292920|MP|A/Black-headed gull/Netherlands/12/2022|EPI ISL 16524203||A / H5N1
EPI2766660|MP|A/common tern/Croatia/44-1/2023|EPI ISL 17762861||A / H5N1
EPI2764010|MP|A/Buteo buteo/Belgium/00815 0002/2023|EPI ISL 18043919||A / H5N1
EPI2764148|MP|A/Sterna hirundo/Belgium/05884 0002/2023|EPI ISL 18343464||A / H5N1
EPI2651959|MP|A/black-headed gull/Poland/MB141-T/2023|EPI ISL 18033205||A / H5N1
33
EPI2652050|MP|A/black-headed gull/Poland/MB159/2023|EPI ISL 18033218||A / H5N1
47
EPI2552804|MP|A/black-headed gull/Czech Republic/6634-3/2023|EPI ISL 17650918||A / H5N1
EPI2794072|MP|A/common tern/Latvia/26493-1 23VIR8835-10/2023|EPI ISL 18509065||A / H5N1
EPI2837083|MP|A/mink/Finland/688/2023|EPI ISL 18618377||A / H5N1
EPI2777657|MP|A/european herring gull/Leningrad region/RII-WD404/2023|EPI ISL 18422467||A / H5N1
EPI2764139|MP|A/Columba palumbus/Belgium/04695 0003/2023|EPI ISL 18343463||A / H5N1
EPI2760831|MP|A/Guillemot/Wales/402694/2023|EPI ISL 18325139||A / H5N1
EPI2760369|MP|A/black-headed gull/Finland/8254 23VIR7026-19/2023|EPI ISL 18321756||A / H5N1
EPI2764203|MP|A/Chroicocephalus ridibundus/Belgium/04326 0003/2023|EPI ISL 18343474||A / H5N1
EPI2841550|MP|A/Falco peregrinus/Belgium/02424 0002/2023|EPI ISL 18643547||A / H5N1
EPI2794139|MP|A/black-headed gull/Latvia/29104-4 23VIR8835-4/2023|EPI ISL 18509074||A / H5N1
EPI3077690|MP|A/Black-legged kittiwake/Norway/2023-07-1645-1k/2023|EPI ISL 18939117||A / H5N1
EPI3115356|MP|A/Larus ridibundus/Spain/CR4063/2023|EPI ISL 18983379||A / H5N1
EPI2794526|MP|A/peregrine falcon /Sweden/SVA231017SZ0445/FB004185/U-2023|EPI ISL 18514139||A / H5N1
EPI2792254|MP|A/black-headed gull/Leningrad region/RII-WD260Mt/2023|EPI ISL 18501399||A / H5N1
EPI2764345|MP|A/Vulpes vulpes/Belgium/04757 0004/2023|EPI ISL 18347730||A / H5N1
ref|20|A/chicken/Iraq/1/2020-like
0.0020

G)

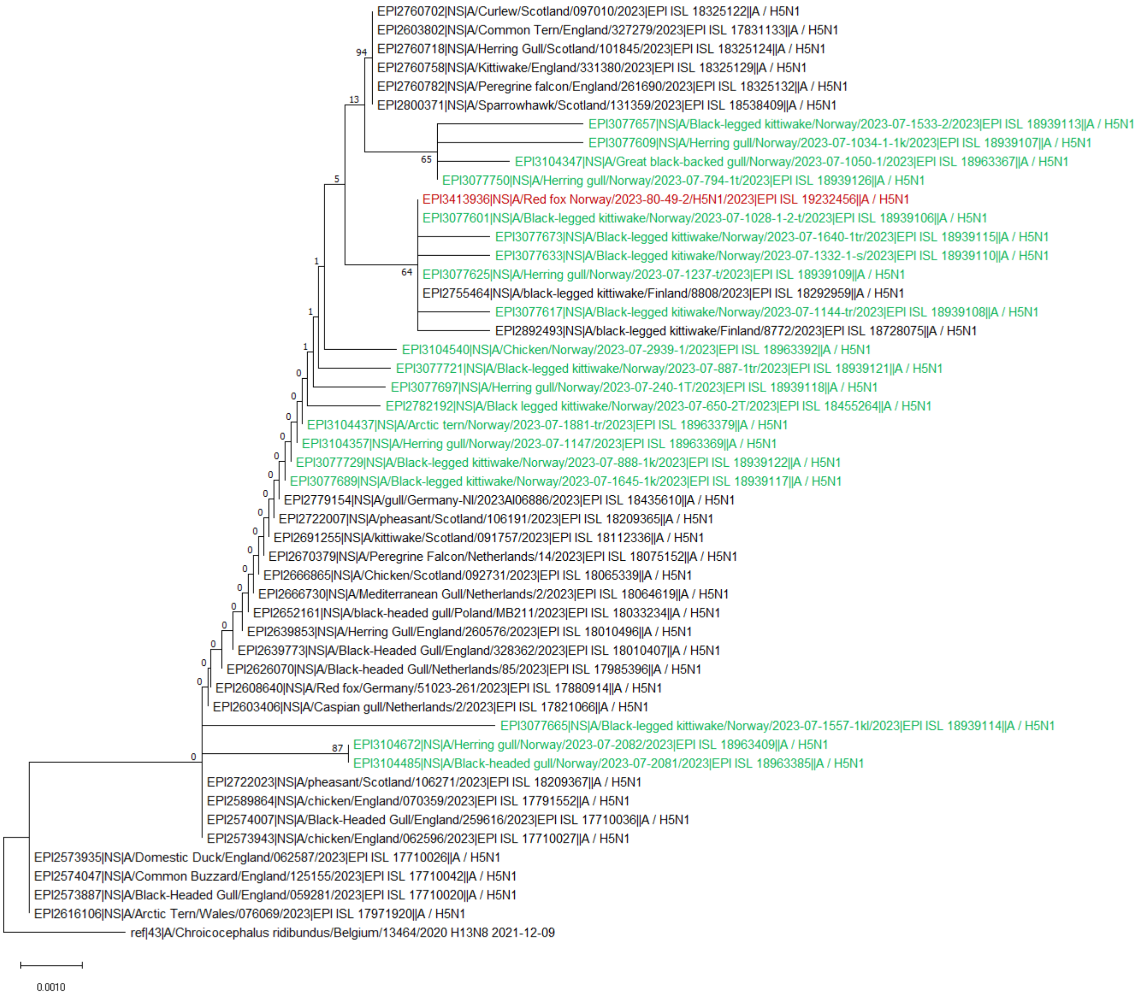

EPI2760702|NS|A/Curlew/Scotland/097010/2023|EPI ISL 18325122||A / H5N1
EPI2603802|NS|A/Common Tern/England/327279/2023|EPI ISL 17831133||A / H5N1
EPI2760718|NS|A/Herring Gull/Scotland/101845/2023|EPI ISL 18325124||A / H5N1
EPI2760758|NS|A/Kittiwake/England/331380/2023|EPI ISL 18325129||A / H5N1
EPI2760782|NS|A/Peregrine falcon/England/261690/2023|EPI ISL 18325132||A / H5N1
EPI2800371|NS|A/Sparrowhawk/Scotland/131359/2023|EPI ISL 18538409||A / H5N1
EPI3077657|NS|A/Black-legged kittiwake/Norway/2023-07-1533-2/2023|EPI ISL 18939113||A / H5N1
EPI3077609|NS|A/Herring gull/Norway/2023-07-1034-1-1k/2023|EPI ISL 18939107||A / H5N1
EPI3104347|NS|A/Great black-backed gull/Norway/2023-07-1050-1/2023|EPI ISL 18963367||A / H5N1
EPI3077750|NS|A/Herring gull/Norway/2023-07-794-1t/2023|EPI ISL 18939126||A / H5N1
EPI3413936|NS|A/Red fox Norway/2023-80-49-2/H5N1/2023|EPI ISL 19232456||A / H5N1
EPI3077601|NS|A/Black-legged kittiwake/Norway/2023-07-1028-1-2-t/2023|EPI ISL 18939106||A / H5N1
EPI3077673|NS|A/Black-legged kittiwake/Norway/2023-07-1640-1tr/2023|EPI ISL 18939115||A / H5N1
EPI3077633|NS|A/Black-legged kitiwake/Norway/2023-07-1332-1-s/2023|EPI ISL 18939110||A / H5N1
EPI3077625|NS|A/Herring gull/Norway/2023-07-1237-t/2023|EPI ISL 18939109||A / H5N1
EPI2755464|NS|A/black-legged kittiwake/Finland/8808/2023|EPI ISL 18292959||A / H5N1
EPI3077617|NS|A/Black-legged kitiwake/Norway/2023-07-1144-tr/2023|EPI ISL 18939108||A / H5N1
EPI2892493|NS|A/black-legged kittiwake/Finland/8772/2023|EPI ISL 18728075||A / H5N1
EPI3104540|NS|A/Chicken/Norway/2023-07-2939-1/2023|EPI ISL 18963392||A / H5N1
EPI3077721|NS|A/Black-legged kittiwake/Norway/2023-07-887-1tr/2023|EPI ISL 18939121||A / H5N1
EPI3077697|NS|A/Herring gull/Norway/2023-07-240-1T/2023|EPI ISL 18939118||A / H5N1
EPI2782192|NS|A/Black legged kittiwake/Norway/2023-07-650-2T/2023|EPI ISL 18455264||A / H5N1
EPI3104437|NS|A/Arctic tern/Norway/2023-07-1881-tr/2023|EPI ISL 18963379||A / H5N1
EPI3104357|NS|A/Herring gull/Norway/2023-07-1147/2023|EPI ISL 18963369||A / H5N1
EPI3077729|NS|A/Black-legged kittiwake/Norway/2023-07-888-1k/2023|EPI ISL 18939122||A / H5N1
EPI3077689|NS|A/Black-legged kittiwake/Norway/2023-07-1645-1k/2023|EPI ISL 18939117||A / H5N1
EPI2779154|NS|A/gull/Germany-NI/2023AI06886/2023|EPI ISL 18435610||A / H5N1
EPI2722007|NS|A/pheasant/Scotland/106191/2023|EPI ISL 18209365||A / H5N1
EPI2691255|NS|A/kittiwake/Scotland/091757/2023|EPI ISL 18112336||A / H5N1
EPI2670379|NS|A/Peregrine Falcon/Netherlands/14/2023|EPI ISL 18075152||A / H5N1
EPI2666865|NS|A/Chicken/Scotland/092731/2023|EPI ISL 18065339||A / H5N1
EPI2666730|NS|A/Mediterranean Gull/Netherlands/2/2023|EPI ISL 18064619||A / H5N1
EPI2652161|NS|A/black-headed gull/Poland/MB211/2023|EPI ISL 18033234||A / H5N1
EPI2639853|NS|A/Herring Gull/England/260576/2023|EPI ISL 18010496||A / H5N1
EPI2639773|NS|A/Black-Headed Gull/England/328362/2023|EPI ISL 18010407||A / H5N1
EPI2626070|NS|A/Black-headed Gull/Netherlands/85/2023|EPI ISL 17985396||A / H5N1
EPI2608640|NS|A/Red fox/Germany/51023-261/2023|EPI ISL 17880914||A / H5N1
EPI2603406|NS|A/Caspian gull/Netherlands/2/2023|EPI ISL 17821066||A / H5N1
EPI3077665|NS|A/Black-legged kittiwake/Norway/2023-07-1557-1kl/2023|EPI ISL 18939114||A / H5N1
EPI3104672|NS|A/Herring gull/Norway/2023-07-2082/2023|EPI ISL 18963409||A / H5N1
EPI3104485|NS|A/Black-headed gull/Norway/2023-07-2081/2023|EPI ISL 18963385||A / H5N1
EPI2722023|NS|A/pheasant/Scotland/106271/2023|EPI ISL 18209367||A / H5N1
EPI2589864|NS|A/chicken/England/070359/2023|EPI ISL 17791552||A / H5N1
EPI2574007|NS|A/Black-Headed Gull/England/259616/2023|EPI ISL 17710036||A / H5N1
EPI2573943|NS|A/chicken/England/062596/2023|EPI ISL 17710027||A / H5N1
EPI2573935|NS|A/Domestic Duck/England/062587/2023|EPI ISL 17710026||A / H5N1
EPI2574047|NS|A/Common Buzzard/England/125155/2023|EPI ISL 17710042||A / H5N1
EPI2573887|NS|A/Black-Headed Gull/England/059281/2023|EPI ISL 17710020||A / H5N1
EPI2616106|NS|A/Arctic Tern/Wales/076069/2023|EPI ISL 17971920||A / H5N1
ref|43|A/Chroicocephalus ridibundus/Belgium/13464/2020 H13N8 2021-12-09
0.0010

**Appendix Figure 5.** Midpoint rooted maximum likelihood phylogenetic trees showing the genetic relationship between highly pathogenic avian influenza H5N5 EA-2021-I viruses from red foxes (red), wild birds (green) and other highly similar viruses obtained from GISAID (black). Trees are shown for the gene segments coding A) PB2, B) PB1, C) PA, D) NP, E) NA, F) M and G) NS.

A)

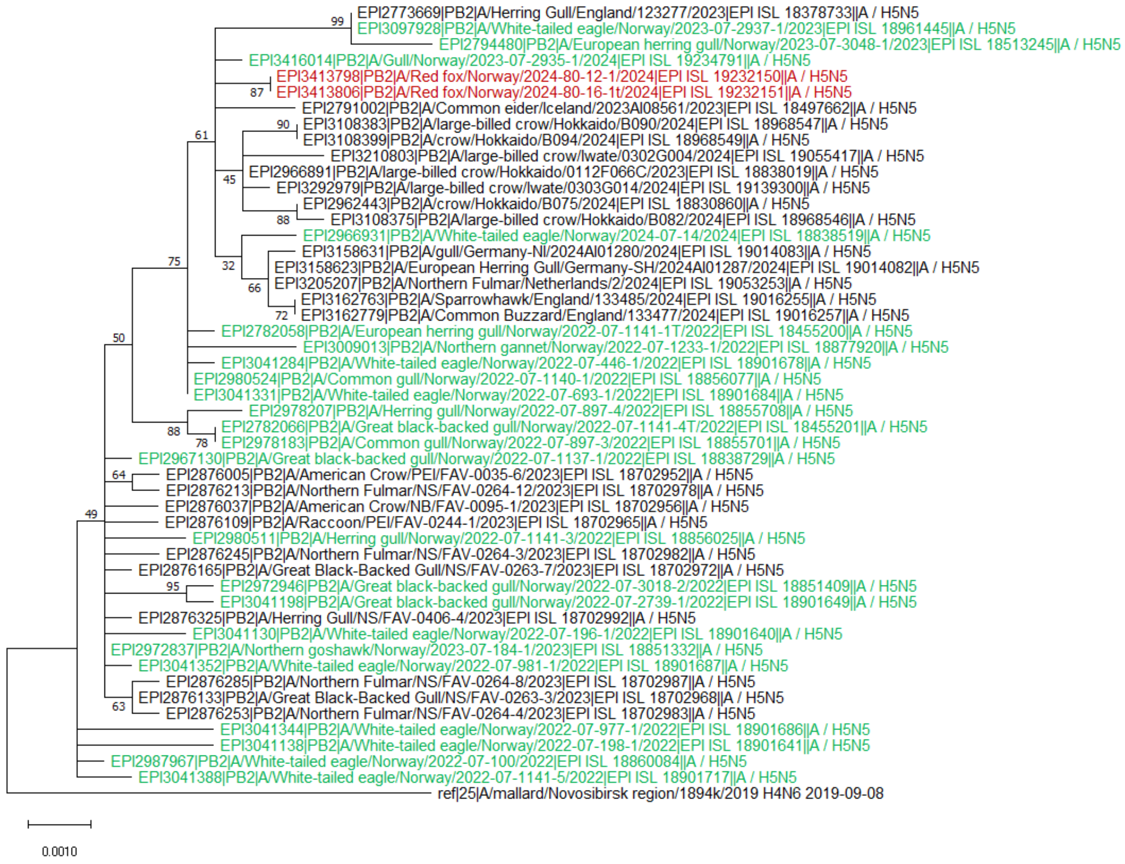

B)

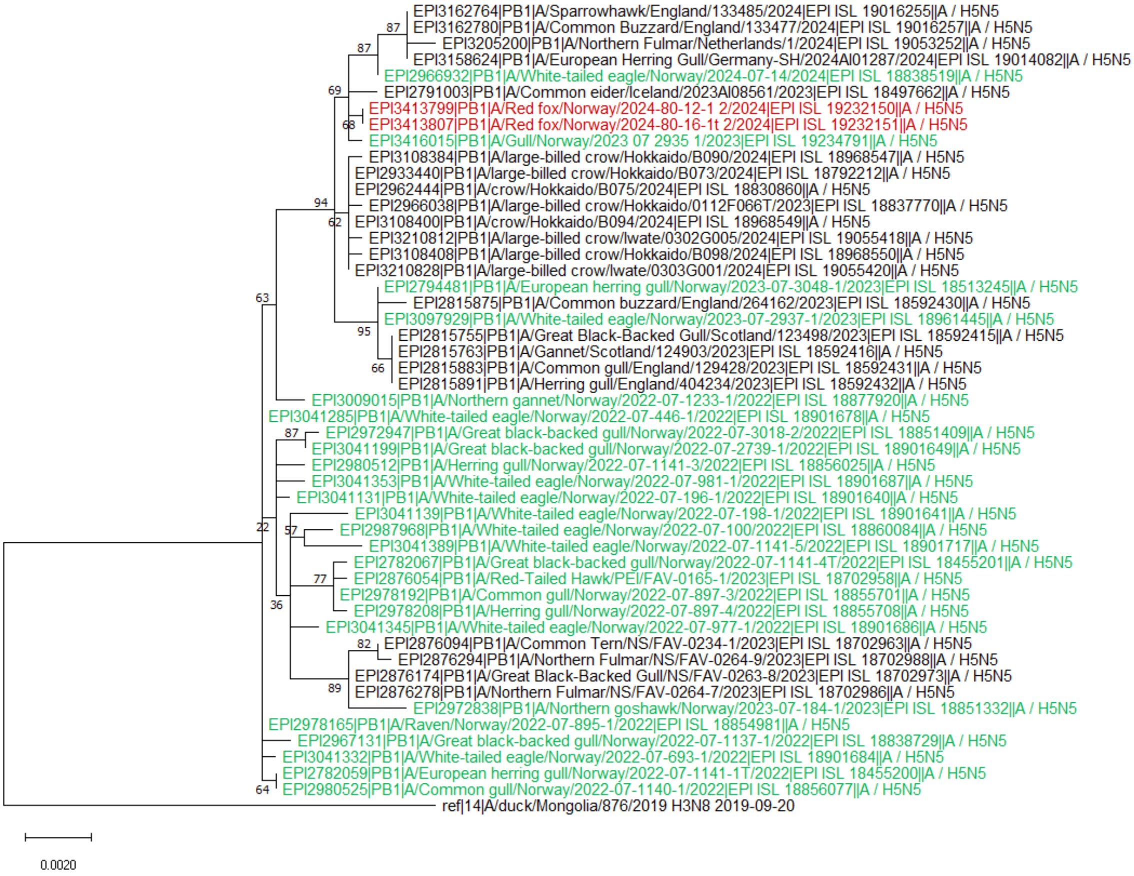
EPI3162764|PB1|A/Sparrowhawk/England/133485/2024|EPI ISL 19016255||A / H5N5
EPI3162780|PB1|A/Common Buzzard/England/133477/2024|EPI ISL 19016257||A / H5N5
EPI3205200|PB1|A/Northern Fulmar/Netherlands/1/2024|EPI ISL 19053252||A / H5N5
EPI3158624|PB1|A/European Herring Gull/Germany-SH/2024AI01287/2024|EPI ISL 19014082||A / H5N5
EPI2966932|PB1|A/White-tailed eagle/Norway/2024-07-14/2024|EPI ISL 18838519||A / H5N5
EPI2791003|PB1|A/Common eider/Iceland/2023AI08561/2023|EPI ISL 18497662||A / H5N5
EPI3413799|PB1|A/Red fox/Norway/2024-80-12-1 2/2024|EPI ISL 19232150||A / H5N5
EPI3413807|PB1|A/Red fox/Norway/2024-80-16-1t 2/2024|EPI ISL 19232151||A / H5N5
EPI3416015|PB1|A/Gull/Norway/2023 07 2935 1/2023|EPI ISL 19234791||A / H5N5
EPI3108384|PB1|A/large-billed crow/Hokkaido/B090/2024|EPI ISL 18968547||A / H5N5
EPI2933440|PB1|A/large-billed crow/Hokkaido/B073/2024|EPI ISL 18792212||A / H5N5
EPI2962444|PB1|A/crow/Hokkaido/B075/2024|EPI ISL 18830860||A / H5N5
EPI2966038|PB1|A/large-billed crow/Hokkaido/0112F066T/2023|EPI ISL 18837770||A / H5N5
EPI3108400|PB1|A/crow/Hokkaido/B094/2024|EPI ISL 18968549||A / H5N5
EPI3210812|PB1|A/large-billed crow/Iwate/0302G005/2024|EPI ISL 19055418||A / H5N5
EPI3108408|PB1|A/large-billed crow/Hokkaido/B098/2024|EPI ISL 18968550||A / H5N5
EPI3210828|PB1|A/large-billed crow/Iwate/0303G001/2024|EPI ISL 19055420||A / H5N5
EPI2794481|PB1|A/European herring gull/Norway/2023-07-3048-1/2023|EPI ISL 18513245||A / H5N5
EPI2815875|PB1|A/Common buzzard/England/264162/2023|EPI ISL 18592430||A / H5N5
EPI3097929|PB1|A/White-tailed eagle/Norway/2023-07-2937-1/2023|EPI ISL 18961445||A / H5N5
EPI2815755|PB1|A/Great Black-Backed Gull/Scotland/123498/2023|EPI ISL 18592415||A / H5N5
EPI2815763|PB1|A/Gannet/Scotland/124903/2023|EPI ISL 18592416||A / H5N5
EPI2815883|PB1|A/Common gull/England/129428/2023|EPI ISL 18592431||A / H5N5
EPI2815891|PB1|A/Herring gull/England/404234/2023|EPI ISL 18592432||A / H5N5
EPI3009015|PB1|A/Northern gannet/Norway/2022-07-1233-1/2022|EPI ISL 18877920||A / H5N5
EPI3041285|PB1|A/White-tailed eagle/Norway/2022-07-446-1/2022|EPI ISL 18901678||A / H5N5
EPI2972947|PB1|A/Great black-backed gull/Norway/2022-07-3018-2/2022|EPI ISL 18851409||A / H5N5
EPI3041199|PB1|A/Great black-backed gull/Norway/2022-07-2739-1/2022|EPI ISL 18901649||A / H5N5
EPI2980512|PB1|A/Herring gull/Norway/2022-07-1141-3/2022|EPI ISL 18856025||A / H5N5
EPI3041353|PB1|A/White-tailed eagle/Norway/2022-07-981-1/2022|EPI ISL 18901687||A / H5N5
EPI3041131|PB1|A/White-tailed eagle/Norway/2022-07-196-1/2022|EPI ISL 18901640||A / H5N5
EPI3041139|PB1|A/White-tailed eagle/Norway/2022-07-198-1/2022|EPI ISL 18901641||A / H5N5
EPI2987968|PB1|A/White-tailed eagle/Norway/2022-07-100/2022|EPI ISL 18860084||A / H5N5
EPI3041389|PB1|A/White-tailed eagle/Norway/2022-07-1141-5/2022|EPI ISL 18901717||A / H5N5
EPI2782067|PB1|A/Great black-backed gull/Norway/2022-07-1141-4T/2022|EPI ISL 18455201||A / H5N5
EPI2876054|PB1|A/Red-Tailed Hawk/PEI/FAV-0165-1/2023|EPI ISL 18702958||A / H5N5
EPI2978192|PB1|A/Common gull/Norway/2022-07-897-3/2022|EPI ISL 18855701||A / H5N5
EPI2978208|PB1|A/Herring gull/Norway/2022-07-897-4/2022|EPI ISL 18855708||A / H5N5
EPI3041345|PB1|A/White-tailed eagle/Norway/2022-07-977-1/2022|EPI ISL 18901686||A / H5N5
EPI2876094|PB1|A/Common Tern/NS/FAV-0234-1/2023|EPI ISL 18702963||A / H5N5
EPI2876294|PB1|A/Northern Fulmar/NS/FAV-0264-9/2023|EPI ISL 18702988||A / H5N5
EPI2876174|PB1|A/Great Black-Backed Gull/NS/FAV-0263-8/2023|EPI ISL 18702973||A / H5N5
EPI2876278|PB1|A/Northern Fulmar/NS/FAV-0264-7/2023|EPI ISL 18702986||A / H5N5
EPI2972838|PB1|A/Northern goshawk/Norway/2023-07-184-1/2023|EPI ISL 18851332||A / H5N5
EPI2978165|PB1|A/Raven/Norway/2022-07-895-1/2022|EPI ISL 18854981||A / H5N5
EPI2967131|PB1|A/Great black-backed gull/Norway/2022-07-1137-1/2022|EPI ISL 18838729||A / H5N5
EPI3041332|PB1|A/White-tailed eagle/Norway/2022-07-693-1/2022|EPI ISL 18901684||A / H5N5
EPI2782059|PB1|A/European herring gull/Norway/2022-07-1141-1T/2022|EPI ISL 18455200||A / H5N5
EPI2980525|PB1|A/Common gull/Norway/2022-07-1140-1/2022|EPI ISL 18856077||A / H5N5
ref|14|A/duck/Mongolia/876/2019 H3N8 2019-09-20
0.0020

C)

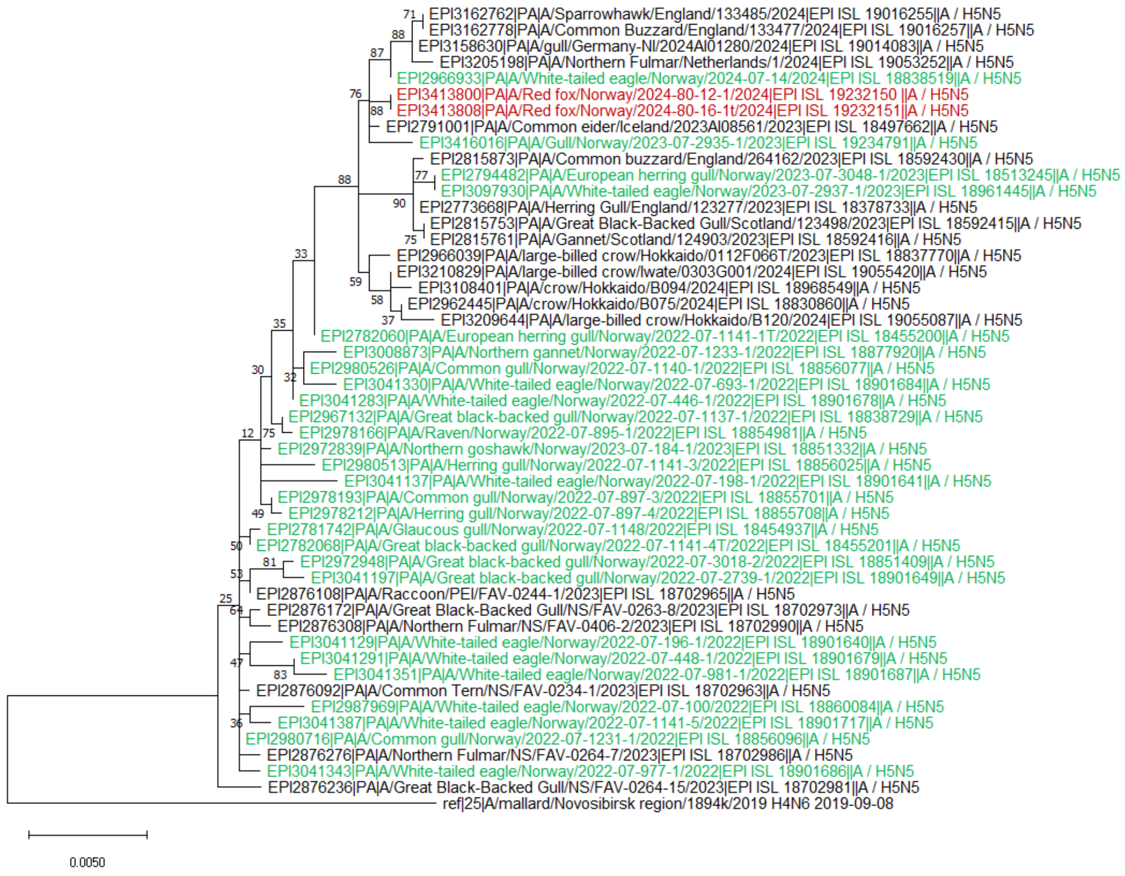

EPI3162762|PA|A/Sparrowhawk/England/133485/2024|EPI ISL 19016255||A / H5N5
EPI3162778|PA|A/Common Buzzard/England/133477/2024|EPI ISL 19016257||A / H5N5
EPI3158630|PA|A/gull/Germany-NI/2024AI01280/2024|EPI ISL 19014083||A / H5N5
EPI3205198|PA|A/Northern Fulmar/Netherlands/1/2024|EPI ISL 19053252||A / H5N5
EPI2966933|PA|A/White-tailed eagle/Norway/2024-07-14/2024|EPI ISL 18838519||A / H5N5
EPI3413800|PA|A/Red fox/Norway/2024-80-12-1/2024|EPI ISL 19232150 ||A / H5N5
EPI3413808|PA|A/Red fox/Norway/2024-80-16-1t/2024|EPI ISL 19232151||A / H5N5
EPI2791001|PA|A/Common eider/Iceland/2023AI08561/2023|EPI ISL 18497662||A / H5N5
EPI3416016|PA|A/Gull/Norway/2023-07-2935-1/2023|EPI ISL 19234791||A / H5N5
EPI2815873|PA|A/Common buzzard/England/264162/2023|EPI ISL 18592430||A / H5N5
EPI2794482|PA|A/European herring gull/Norway/2023-07-3048-1/2023|EPI ISL 18513245||A / H5N5
EPI3097930|PA|A/White-tailed eagle/Norway/2023-07-2937-1/2023|EPI ISL 18961445||A / H5N5
EPI2773668|PA|A/Herring Gull/England/123277/2023|EPI ISL 18378733||A / H5N5
EPI2815753|PA|A/Great Black-Backed Gull/Scotland/123498/2023|EPI ISL 18592415||A / H5N5
EPI2815761|PA|A/Gannet/Scotland/124903/2023|EPI ISL 18592416||A / H5N5
EPI2966039|PA|A/large-billed crow/Hokkaido/0112F066T/2023|EPI ISL 18837770||A / H5N5
EPI3210829|PA|A/large-billed crow/Iwate/0303G001/2024|EPI ISL 19055420||A / H5N5
EPI3108401|PA|A/crow/Hokkaido/B094/2024|EPI ISL 18968549||A / H5N5
EPI2962445|PA|A/crow/Hokkaido/B075/2024|EPI ISL 18830860||A / H5N5
EPI3209644|PA|A/large-billed crow/Hokkaido/B120/2024|EPI ISL 19055087||A / H5N5
EPI2782060|PA|A/European herring gull/Norway/2022-07-1141-1T/2022|EPI ISL 18455200||A / H5N5
EPI3008873|PA|A/Northern gannet/Norway/2022-07-1233-1/2022|EPI ISL 18877920||A / H5N5
EPI2980526|PA|A/Common gull/Norway/2022-07-1140-1/2022|EPI ISL 18856077||A / H5N5
EPI3041330|PA|A/White-tailed eagle/Norway/2022-07-693-1/2022|EPI ISL 18901684||A / H5N5
EPI3041283|PA|A/White-tailed eagle/Norway/2022-07-446-1/2022|EPI ISL 18901678||A / H5N5
EPI2967132|PA|A/Great black-backed gull/Norway/2022-07-1137-1/2022|EPI ISL 18838729||A / H5N5
EPI2978166|PA|A/Raven/Norway/2022-07-895-1/2022|EPI ISL 18854981||A / H5N5
EPI2972839|PA|A/Northern goshawk/Norway/2023-07-184-1/2023|EPI ISL 18851332||A / H5N5
EPI2980513|PA|A/Herring gull/Norway/2022-07-1141-3/2022|EPI ISL 18856025||A / H5N5
EPI3041137|PA|A/White-tailed eagle/Norway/2022-07-198-1/2022|EPI ISL 18901641||A / H5N5
EPI2978193|PA|A/Common gull/Norway/2022-07-897-3/2022|EPI ISL 18855701||A / H5N5
EPI2978212|PA|A/Herring gull/Norway/2022-07-897-4/2022|EPI ISL 18855708||A / H5N5
EPI2781742|PA|A/Glaucous gull/Norway/2022-07-1148/2022|EPI ISL 18454937||A / H5N5
EPI2782068|PA|A/Great black-backed gull/Norway/2022-07-1141-4T/2022|EPI ISL 18455201||A / H5N5
EPI2972948|PA|A/Great black-backed gull/Norway/2022-07-3018-2/2022|EPI ISL 18851409||A / H5N5
EPI3041197|PA|A/Great black-backed gull/Norway/2022-07-2739-1/2022|EPI ISL 18901649||A / H5N5
EPI2876108|PA|A/Raccoon/PEI/FAV-0244-1/2023|EPI ISL 18702965||A / H5N5
EPI2876172|PA|A/Great Black-Backed Gull/NS/FAV-0263-8/2023|EPI ISL 18702973||A / H5N5
EPI2876308|PA|A/Northern Fulmar/NS/FAV-0406-2/2023|EPI ISL 18702990||A / H5N5
EPI3041129|PA|A/White-tailed eagle/Norway/2022-07-196-1/2022|EPI ISL 18901640||A / H5N5
EPI3041291|PA|A/White-tailed eagle/Norway/2022-07-448-1/2022|EPI ISL 18901679||A / H5N5
EPI3041351|PA|A/White-tailed eagle/Norway/2022-07-981-1/2022|EPI ISL 18901687||A / H5N5
EPI2876092|PA|A/Common Tern/NS/FAV-0234-1/2023|EPI ISL 18702963||A / H5N5
EPI2987969|PA|A/White-tailed eagle/Norway/2022-07-100/2022|EPI ISL 18860084||A / H5N5
EPI3041387|PA|A/White-tailed eagle/Norway/2022-07-1141-5/2022|EPI ISL 18901717||A / H5N5
EPI2980716|PA|A/Common gull/Norway/2022-07-1231-1/2022|EPI ISL 18856096||A / H5N5
EPI2876276|PA|A/Northern Fulmar/NS/FAV-0264-7/2023|EPI ISL 18702986||A / H5N5
EPI3041343|PA|A/White-tailed eagle/Norway/2022-07-977-1/2022|EPI ISL 18901686||A / H5N5
EPI2876236|PA|A/Great Black-Backed Gull/NS/FAV-0264-15/2023|EPI ISL 18702981||A / H5N5
ref|25|A/mallard/Novosibirsk region/1894k/2019 H4N6 2019-09-08
0.0050

D)

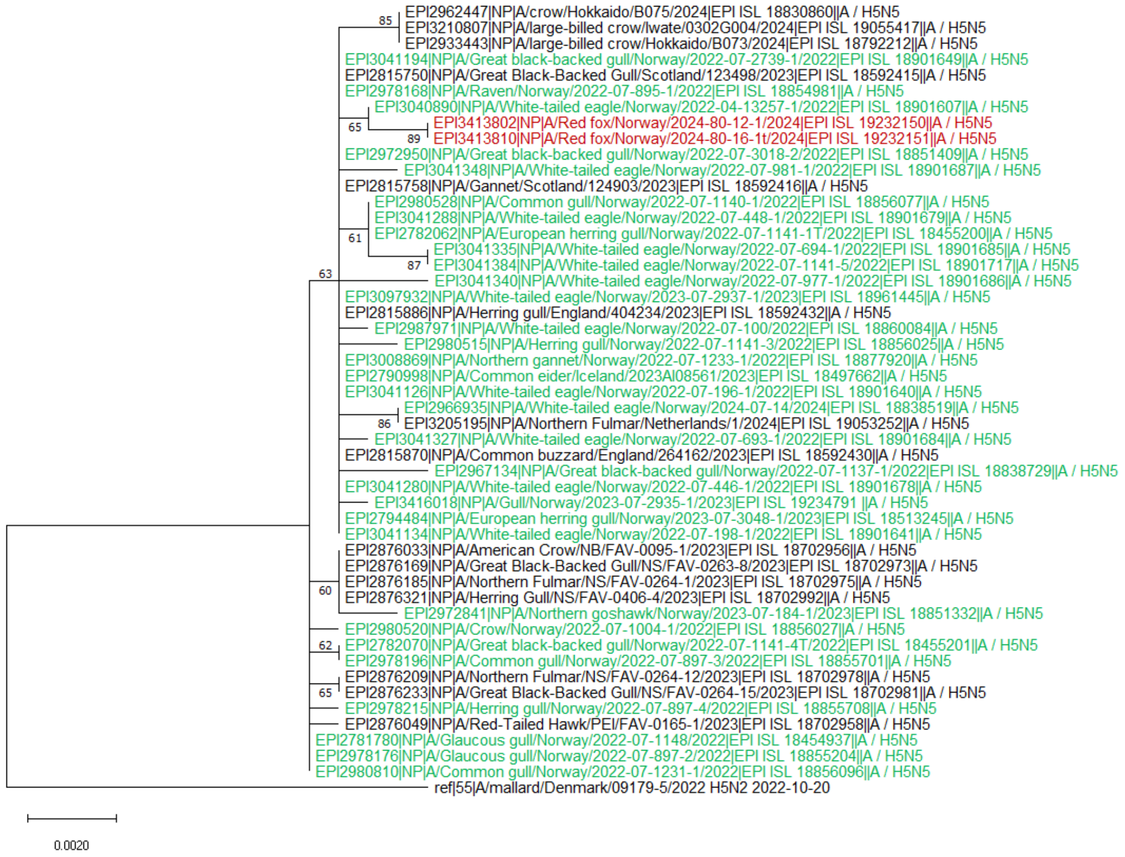

85
EPI2962447|NP|A/crow/Hokkaido/B075/2024|EPI ISL 18830860||A / H5N5
EPI3210807|NP|A/large-billed crow/Iwate/0302G004/2024|EPI ISL 19055417||A / H5N5
EPI2933443|NP|A/large-billed crow/Hokkaido/B073/2024|EPI ISL 18792212||A / H5N5
EPI3041194|NP|A/Great black-backed gull/Norway/2022-07-2739-1/2022|EPI ISL 18901649||A / H5N5
EPI2815750|NP|A/Great Black-Backed Gull/Scotland/123498/2023|EPI ISL 18592415||A / H5N5
EPI2978168|NP|A/Raven/Norway/2022-07-895-1/2022|EPI ISL 18854981||A / H5N5
EPI3040890|NP|A/White-tailed eagle/Norway/2022-04-13257-1/2022|EPI ISL 18901607||A / H5N5
65
EPI3413802|NP|A/Red fox/Norway/2024-80-12-1/2024|EPI ISL 19232150||A / H5N5
89
EPI3413810|NP|A/Red fox/Norway/2024-80-16-1t/2024|EPI ISL 19232151||A / H5N5
EPI2972950|NP|A/Great black-backed gull/Norway/2022-07-3018-2/2022|EPI ISL 18851409||A / H5N5
EPI3041348|NP|A/White-tailed eagle/Norway/2022-07-981-1/2022|EPI ISL 18901687||A / H5N5
EPI2815758|NP|A/Gannet/Scotland/124903/2023|EPI ISL 18592416||A / H5N5
EPI2980528|NP|A/Common gull/Norway/2022-07-1140-1/2022|EPI ISL 18856077||A / H5N5
EPI3041288|NP|A/White-tailed eagle/Norway/2022-07-448-1/2022|EPI ISL 18901679||A / H5N5
61
EPI2782062|NP|A/European herring gull/Norway/2022-07-1141-1T/2022|EPI ISL 18455200||A / H5N5
EPI3041335|NP|A/White-tailed eagle/Norway/2022-07-694-1/2022|EPI ISL 18901685||A / H5N5
87
EPI3041384|NP|A/White-tailed eagle/Norway/2022-07-1141-5/2022|EPI ISL 18901717||A / H5N5
63
EPI3041340|NP|A/White-tailed eagle/Norway/2022-07-977-1/2022|EPI ISL 18901686||A / H5N5
EPI3097932|NP|A/White-tailed eagle/Norway/2023-07-2937-1/2023|EPI ISL 18961445||A / H5N5
EPI2815886|NP|A/Herring gull/England/404234/2023|EPI ISL 18592432||A / H5N5
EPI2987971|NP|A/White-tailed eagle/Norway/2022-07-100/2022|EPI ISL 18860084||A / H5N5
EPI2980515|NP|A/Herring gull/Norway/2022-07-1141-3/2022|EPI ISL 18856025||A / H5N5
EPI3008869|NP|A/Northern gannet/Norway/2022-07-1233-1/2022|EPI ISL 18877920||A / H5N5
EPI2790998|NP|A/Common eider/Iceland/2023AI08561/2023|EPI ISL 18497662||A / H5N5
EPI3041126|NP|A/White-tailed eagle/Norway/2022-07-196-1/2022|EPI ISL 18901640||A / H5N5
EPI2966935|NP|A/White-tailed eagle/Norway/2024-07-14/2024|EPI ISL 18838519||A / H5N5
86
EPI3205195|NP|A/Northern Fulmar/Netherlands/1/2024|EPI ISL 19053252||A / H5N5
EPI3041327|NP|A/White-tailed eagle/Norway/2022-07-693-1/2022|EPI ISL 18901684||A / H5N5
EPI2815870|NP|A/Common buzzard/England/264162/2023|EPI ISL 18592430||A / H5N5
EPI2967134|NP|A/Great black-backed gull/Norway/2022-07-1137-1/2022|EPI ISL 18838729||A / H5N5
EPI3041280|NP|A/White-tailed eagle/Norway/2022-07-446-1/2022|EPI ISL 18901678||A / H5N5
EPI3416018|NP|A/Gull/Norway/2023-07-2935-1/2023|EPI ISL 19234791 ||A / H5N5
EPI2794484|NP|A/European herring gull/Norway/2023-07-3048-1/2023|EPI ISL 18513245||A / H5N5
EPI3041134|NP|A/White-tailed eagle/Norway/2022-07-198-1/2022|EPI ISL 18901641||A / H5N5
EPI2876033|NP|A/American Crow/NB/FAV-0095-1/2023|EPI ISL 18702956||A / H5N5
EPI2876169|NP|A/Great Black-Backed Gull/NS/FAV-0263-8/2023|EPI ISL 18702973||A / H5N5
EPI2876185|NP|A/Northern Fulmar/NS/FAV-0264-1/2023|EPI ISL 18702975||A / H5N5
60
EPI2876321|NP|A/Herring Gull/NS/FAV-0406-4/2023|EPI ISL 18702992||A / H5N5
EPI2972841|NP|A/Northern goshawk/Norway/2023-07-184-1/2023|EPI ISL 18851332||A / H5N5
EPI2980520|NP|A/Crow/Norway/2022-07-1004-1/2022|EPI ISL 18856027||A / H5N5
62
EPI2782070|NP|A/Great black-backed gull/Norway/2022-07-1141-4T/2022|EPI ISL 18455201||A / H5N5
EPI2978196|NP|A/Common gull/Norway/2022-07-897-3/2022|EPI ISL 18855701||A / H5N5
EPI2876209|NP|A/Northern Fulmar/NS/FAV-0264-12/2023|EPI ISL 18702978||A / H5N5
65
EPI2876233|NP|A/Great Black-Backed Gull/NS/FAV-0264-15/2023|EPI ISL 18702981||A / H5N5
EPI2978215|NP|A/Herring gull/Norway/2022-07-897-4/2022|EPI ISL 18855708||A / H5N5
EPI2876049|NP|A/Red-Tailed Hawk/PEI/FAV-0165-1/2023|EPI ISL 18702958||A / H5N5
EPI2781780|NP|A/Glaucous gull/Norway/2022-07-1148/2022|EPI ISL 18454937||A / H5N5
EPI2978176|NP|A/Glaucous gull/Norway/2022-07-897-2/2022|EPI ISL 18855204||A / H5N5
EPI2980810|NP|A/Common gull/Norway/2022-07-1231-1/2022|EPI ISL 18856096||A / H5N5
ref|55|A/mallard/Denmark/09179-5/2022 H5N2 2022-10-20
0.0020

E)

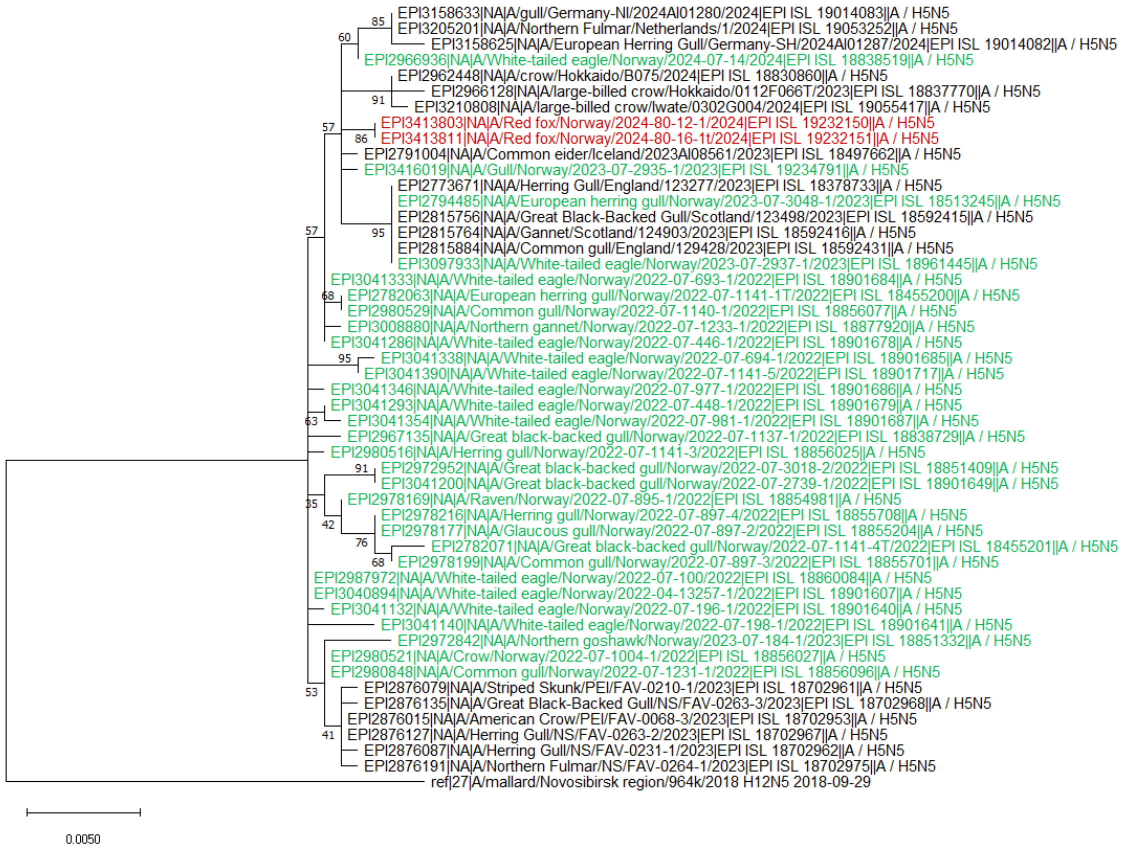
EPI3158633|NA|A/gull/Germany-NI/2024AI01280/2024|EPI ISL 19014083||A / H5N5
EPI3205201|NA|A/Northern Fulmar/Netherlands/1/2024|EPI ISL 19053252||A / H5N5
EPI3158625|NA|A/European Herring Gull/Germany-SH/2024AI01287/2024|EPI ISL 19014082||A / H5N5
EPI2966936|NA|A/White-tailed eagle/Norway/2024-07-14/2024|EPI ISL 18838519||A / H5N5
EPI2962448|NA|A/crow/Hokkaido/B075/2024|EPI ISL 18830860||A / H5N5
EPI2966128|NA|A/large-billed crow/Hokkaido/0112F066T/2023|EPI ISL 18837770||A / H5N5
EPI3210808|NA|A/large-billed crow/Iwate/0302G004/2024|EPI ISL 19055417||A / H5N5
EPI3413803|NA|A/Red fox/Norway/2024-80-12-1/2024|EPI ISL 19232150||A / H5N5
EPI3413811|NA|A/Red fox/Norway/2024-80-16-1t/2024|EPI ISL 19232151||A / H5N5
EPI2791004|NA|A/Common eider/Iceland/2023AI08561/2023|EPI ISL 18497662||A / H5N5
EPI3416019|NA|A/Gull/Norway/2023-07-2935-1/2023|EPI ISL 19234791||A / H5N5
EPI2773671|NA|A/Herring Gull/England/123277/2023|EPI ISL 18378733||A / H5N5
EPI2794485|NA|A/European herring gull/Norway/2023-07-3048-1/2023|EPI ISL 18513245||A / H5N5
EPI2815756|NA|A/Great Black-Backed Gull/Scotland/123498/2023|EPI ISL 18592415||A / H5N5
EPI2815764|NA|A/Gannet/Scotland/124903/2023|EPI ISL 18592416||A / H5N5
EPI2815884|NA|A/Common gull/England/129428/2023|EPI ISL 18592431||A / H5N5
EPI3097933|NA|A/White-tailed eagle/Norway/2023-07-2937-1/2023|EPI ISL 18961445||A / H5N5
EPI3041333|NA|A/White-tailed eagle/Norway/2022-07-693-1/2022|EPI ISL 18901684||A / H5N5
EPI2782063|NA|A/European herring gull/Norway/2022-07-1141-1T/2022|EPI ISL 18455200||A / H5N5
EPI2980529|NA|A/Common gull/Norway/2022-07-1140-1/2022|EPI ISL 18856077||A / H5N5
EPI3008880|NA|A/Northern gannet/Norway/2022-07-1233-1/2022|EPI ISL 18877920||A / H5N5
EPI3041286|NA|A/White-tailed eagle/Norway/2022-07-446-1/2022|EPI ISL 18901678||A / H5N5
EPI3041338|NA|A/White-tailed eagle/Norway/2022-07-694-1/2022|EPI ISL 18901685||A / H5N5
EPI3041390|NA|A/White-tailed eagle/Norway/2022-07-1141-5/2022|EPI ISL 18901717||A / H5N5
EPI3041346|NA|A/White-tailed eagle/Norway/2022-07-977-1/2022|EPI ISL 18901686||A / H5N5
EPI3041293|NA|A/White-tailed eagle/Norway/2022-07-448-1/2022|EPI ISL 18901679||A / H5N5
EPI3041354|NA|A/White-tailed eagle/Norway/2022-07-981-1/2022|EPI ISL 18901687||A / H5N5
EPI2967135|NA|A/Great black-backed gull/Norway/2022-07-1137-1/2022|EPI ISL 18838729||A / H5N5
EPI2980516|NA|A/Herring gull/Norway/2022-07-1141-3/2022|EPI ISL 18856025||A / H5N5
EPI2972952|NA|A/Great black-backed gull/Norway/2022-07-3018-2/2022|EPI ISL 18851409||A / H5N5
EPI3041200|NA|A/Great black-backed gull/Norway/2022-07-2739-1/2022|EPI ISL 18901649||A / H5N5
EPI2978169|NA|A/Raven/Norway/2022-07-895-1/2022|EPI ISL 18854981||A / H5N5
EPI2978216|NA|A/Herring gull/Norway/2022-07-897-4/2022|EPI ISL 18855708||A / H5N5
EPI2978177|NA|A/Glaucous gull/Norway/2022-07-897-2/2022|EPI ISL 18855204||A / H5N5
EPI2782071|NA|A/Great black-backed gull/Norway/2022-07-1141-4T/2022|EPI ISL 18455201||A / H5N5
EPI2978199|NA|A/Common gull/Norway/2022-07-897-3/2022|EPI ISL 18855701||A / H5N5
EPI2987972|NA|A/White-tailed eagle/Norway/2022-07-100/2022|EPI ISL 18860084||A / H5N5
EPI3040894|NA|A/White-tailed eagle/Norway/2022-04-13257-1/2022|EPI ISL 18901607||A / H5N5
EPI3041132|NA|A/White-tailed eagle/Norway/2022-07-196-1/2022|EPI ISL 18901640||A / H5N5
EPI3041140|NA|A/White-tailed eagle/Norway/2022-07-198-1/2022|EPI ISL 18901641||A / H5N5
EPI2972842|NA|A/Northern goshawk/Norway/2023-07-184-1/2023|EPI ISL 18851332||A / H5N5
EPI2980521|NA|A/Crow/Norway/2022-07-1004-1/2022|EPI ISL 18856027||A / H5N5
EPI2980848|NA|A/Common gull/Norway/2022-07-1231-1/2022|EPI ISL 18856096||A / H5N5
EPI2876079|NA|A/Striped Skunk/PEI/FAV-0210-1/2023|EPI ISL 18702961||A / H5N5
EPI2876135|NA|A/Great Black-Backed Gull/NS/FAV-0263-3/2023|EPI ISL 18702968||A / H5N5
EPI2876015|NA|A/American Crow/PEI/FAV-0068-3/2023|EPI ISL 18702953||A / H5N5
EPI2876127|NA|A/Herring Gull/NS/FAV-0263-2/2023|EPI ISL 18702967||A / H5N5
EPI2876087|NA|A/Herring Gull/NS/FAV-0231-1/2023|EPI ISL 18702962||A / H5N5
EPI2876191|NA|A/Northern Fulmar/NS/FAV-0264-1/2023|EPI ISL 18702975||A / H5N5
ref|27|A/mallard/Novosibirsk region/964k/2018 H12N5 2018-09-29
85
60
91
57
86
57
95
68
95
63
91
35
42
76
68
53
41
0.0050

F)

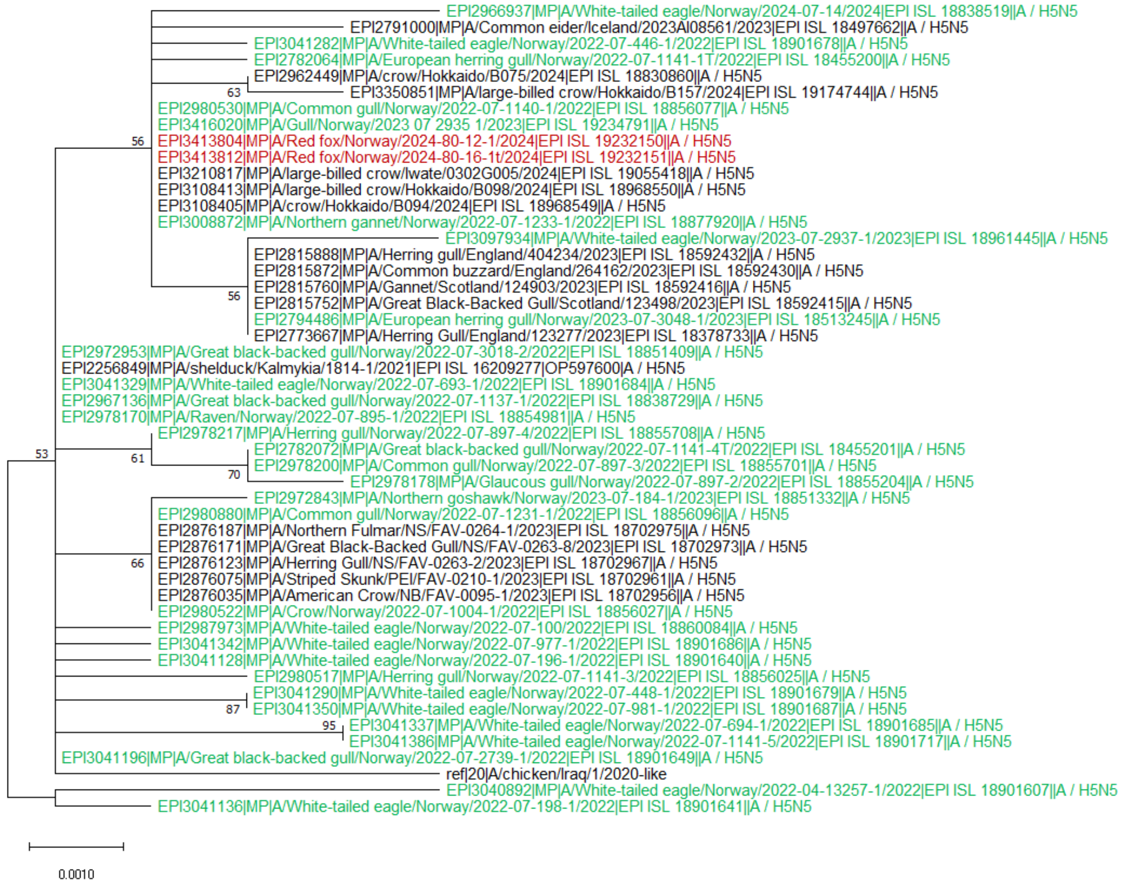
EPI2966937|MP|A/White-tailed eagle/Norway/2024-07-14/2024|EPI ISL 18838519||A / H5N5
EPI2791000|MP|A/Common eider/Iceland/2023AI08561/2023|EPI ISL 18497662||A / H5N5
EPI3041282|MP|A/White-tailed eagle/Norway/2022-07-446-1/2022|EPI ISL 18901678||A / H5N5
EPI2782064|MP|A/European herring gull/Norway/2022-07-1141-1T/2022|EPI ISL 18455200||A / H5N5
EPI2962449|MP|A/crow/Hokkaido/B075/2024|EPI ISL 18830860||A / H5N5
63
EPI3350851|MP|A/large-billed crow/Hokkaido/B157/2024|EPI ISL 19174744||A / H5N5
EPI2980530|MP|A/Common gull/Norway/2022-07-1140-1/2022|EPI ISL 18856077||A / H5N5
EPI3416020|MP|A/Gull/Norway/2023 07 2935 1/2023|EPI ISL 19234791||A / H5N5
56
EPI3413804|MP|A/Red fox/Norway/2024-80-12-1/2024|EPI ISL 19232150||A / H5N5
EPI3413812|MP|A/Red fox/Norway/2024-80-16-1t/2024|EPI ISL 19232151||A / H5N5
EPI3210817|MP|A/large-billed crow/Iwate/0302G005/2024|EPI ISL 19055418||A / H5N5
EPI3108413|MP|A/large-billed crow/Hokkaido/B098/2024|EPI ISL 18968550||A / H5N5
EPI3108405|MP|A/crow/Hokkaido/B094/2024|EPI ISL 18968549||A / H5N5
EPI3008872|MP|A/Northern gannet/Norway/2022-07-1233-1/2022|EPI ISL 18877920||A / H5N5
EPI3097934|MP|A/White-tailed eagle/Norway/2023-07-2937-1/2023|EPI ISL 18961445||A / H5N5
EPI2815888|MP|A/Herring gull/England/404234/2023|EPI ISL 18592432||A / H5N5
EPI2815872|MP|A/Common buzzard/England/264162/2023|EPI ISL 18592430||A / H5N5
EPI2815760|MP|A/Gannet/Scotland/124903/2023|EPI ISL 18592416||A / H5N5
56
EPI2815752|MP|A/Great Black-Backed Gull/Scotland/123498/2023|EPI ISL 18592415||A / H5N5
EPI2794486|MP|A/European herring gull/Norway/2023-07-3048-1/2023|EPI ISL 18513245||A / H5N5
EPI2773667|MP|A/Herring Gull/England/123277/2023|EPI ISL 18378733||A / H5N5
EPI2972953|MP|A/Great black-backed gull/Norway/2022-07-3018-2/2022|EPI ISL 18851409||A / H5N5
EPI2256849|MP|A/shelduck/Kalmykia/1814-1/2021|EPI ISL 16209277|OP597600|A / H5N5
EPI3041329|MP|A/White-tailed eagle/Norway/2022-07-693-1/2022|EPI ISL 18901684||A / H5N5
EPI2967136|MP|A/Great black-backed gull/Norway/2022-07-1137-1/2022|EPI ISL 18838729||A / H5N5
EPI2978170|MP|A/Raven/Norway/2022-07-895-1/2022|EPI ISL 18854981||A / H5N5
EPI2978217|MP|A/Herring gull/Norway/2022-07-897-4/2022|EPI ISL 18855708||A / H5N5
53
61
EPI2782072|MP|A/Great black-backed gull/Norway/2022-07-1141-4T/2022|EPI ISL 18455201||A / H5N5
EPI2978200|MP|A/Common gull/Norway/2022-07-897-3/2022|EPI ISL 18855701||A / H5N5
70
EPI2978178|MP|A/Glaucous gull/Norway/2022-07-897-2/2022|EPI ISL 18855204||A / H5N5
EPI2972843|MP|A/Northern goshawk/Norway/2023-07-184-1/2023|EPI ISL 18851332||A / H5N5
EPI2980880|MP|A/Common gull/Norway/2022-07-1231-1/2022|EPI ISL 18856096||A / H5N5
EPI2876187|MP|A/Northern Fulmar/NS/FAV-0264-1/2023|EPI ISL 18702975||A / H5N5
EPI2876171|MP|A/Great Black-Backed Gull/NS/FAV-0263-8/2023|EPI ISL 18702973||A / H5N5
66
EPI2876123|MP|A/Herring Gull/NS/FAV-0263-2/2023|EPI ISL 18702967||A / H5N5
EPI2876075|MP|A/Striped Skunk/PEI/FAV-0210-1/2023|EPI ISL 18702961||A / H5N5
EPI2876035|MP|A/American Crow/NB/FAV-0095-1/2023|EPI ISL 18702956||A / H5N5
EPI2980522|MP|A/Crow/Norway/2022-07-1004-1/2022|EPI ISL 18856027||A / H5N5
EPI2987973|MP|A/White-tailed eagle/Norway/2022-07-100/2022|EPI ISL 18860084||A / H5N5
EPI3041342|MP|A/White-tailed eagle/Norway/2022-07-977-1/2022|EPI ISL 18901686||A / H5N5
EPI3041128|MP|A/White-tailed eagle/Norway/2022-07-196-1/2022|EPI ISL 18901640||A / H5N5
EPI2980517|MP|A/Herring gull/Norway/2022-07-1141-3/2022|EPI ISL 18856025||A / H5N5
EPI3041290|MP|A/White-tailed eagle/Norway/2022-07-448-1/2022|EPI ISL 18901679||A / H5N5
87
EPI3041350|MP|A/White-tailed eagle/Norway/2022-07-981-1/2022|EPI ISL 18901687||A / H5N5
95
EPI3041337|MP|A/White-tailed eagle/Norway/2022-07-694-1/2022|EPI ISL 18901685||A / H5N5
EPI3041386|MP|A/White-tailed eagle/Norway/2022-07-1141-5/2022|EPI ISL 18901717||A / H5N5
EPI3041196|MP|A/Great black-backed gull/Norway/2022-07-2739-1/2022|EPI ISL 18901649||A / H5N5
ref|20|A/chicken/Iraq/1/2020-like
EPI3040892|MP|A/White-tailed eagle/Norway/2022-04-13257-1/2022|EPI ISL 18901607||A / H5N5
EPI3041136|MP|A/White-tailed eagle/Norway/2022-07-198-1/2022|EPI ISL 18901641||A / H5N5
0.0010

G)

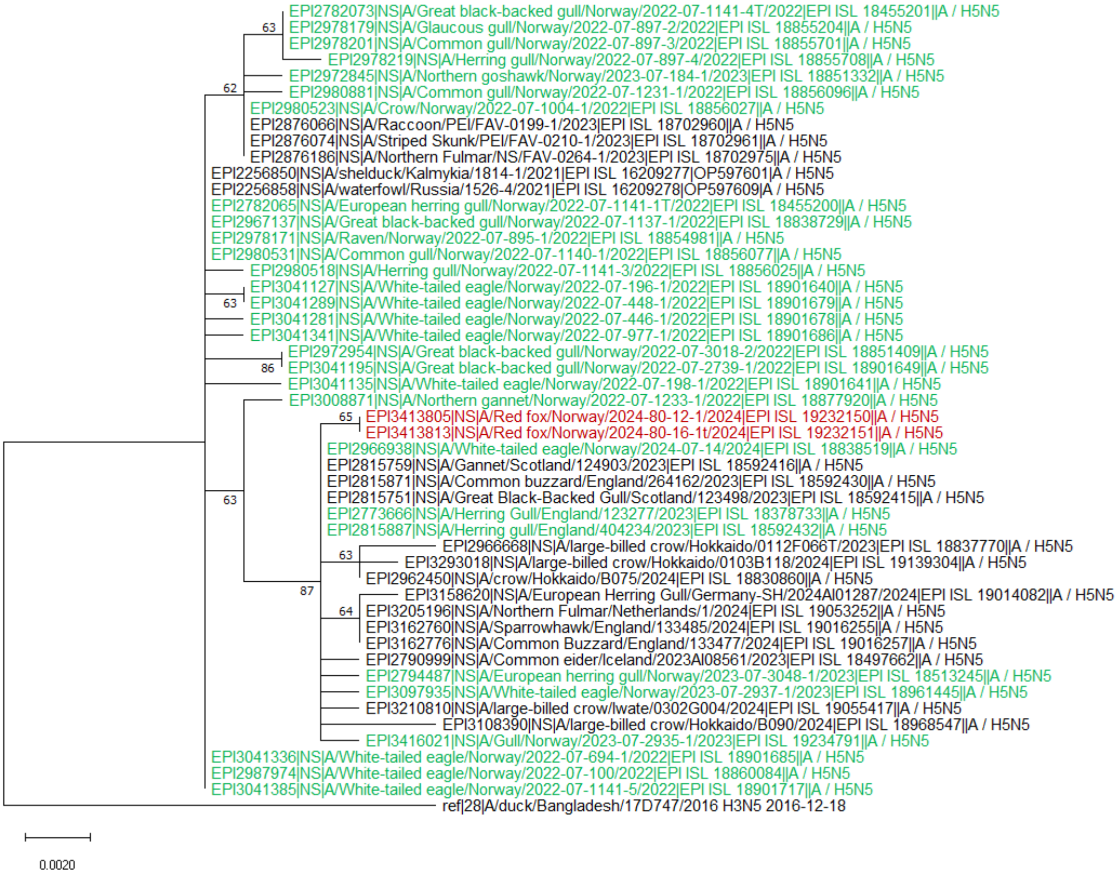
EPI2782073|NS|A/Great black-backed gull/Norway/2022-07-1141-4T/2022|EPI ISL 18455201||A / H5N5
63
EPI2978179|NS|A/Glaucous gull/Norway/2022-07-897-2/2022|EPI ISL 18855204||A / H5N5
EPI2978201|NS|A/Common gull/Norway/2022-07-897-3/2022|EPI ISL 18855701||A / H5N5
EPI2978219|NS|A/Herring gull/Norway/2022-07-897-4/2022|EPI ISL 18855708||A / H5N5
EPI2972845|NS|A/Northern goshawk/Norway/2023-07-184-1/2023|EPI ISL 18851332||A / H5N5
62
EPI2980881|NS|A/Common gull/Norway/2022-07-1231-1/2022|EPI ISL 18856096||A / H5N5
EPI2980523|NS|A/Crow/Norway/2022-07-1004-1/2022|EPI ISL 18856027||A / H5N5
EPI2876066|NS|A/Raccoon/PEI/FAV-0199-1/2023|EPI ISL 18702960||A / H5N5
EPI2876074|NS|A/Striped Skunk/PEI/FAV-0210-1/2023|EPI ISL 18702961||A / H5N5
EPI2876186|NS|A/Northern Fulmar/NS/FAV-0264-1/2023|EPI ISL 18702975||A / H5N5
EPI2256850|NS|A/shelduck/Kalmykia/1814-1/2021|EPI ISL 16209277|OP597601|A / H5N5
EPI2256858|NS|A/waterfowl/Russia/1526-4/2021|EPI ISL 16209278|OP597609|A / H5N5
EPI2782065|NS|A/European herring gull/Norway/2022-07-1141-1T/2022|EPI ISL 18455200||A / H5N5
EPI2967137|NS|A/Great black-backed gull/Norway/2022-07-1137-1/2022|EPI ISL 18838729||A / H5N5
EPI2978171|NS|A/Raven/Norway/2022-07-895-1/2022|EPI ISL 18854981||A / H5N5
EPI2980531|NS|A/Common gull/Norway/2022-07-1140-1/2022|EPI ISL 18856077||A / H5N5
EPI2980518|NS|A/Herring gull/Norway/2022-07-1141-3/2022|EPI ISL 18856025||A / H5N5
EPI3041127|NS|A/White-tailed eagle/Norway/2022-07-196-1/2022|EPI ISL 18901640||A / H5N5
63
EPI3041289|NS|A/White-tailed eagle/Norway/2022-07-448-1/2022|EPI ISL 18901679||A / H5N5
EPI3041281|NS|A/White-tailed eagle/Norway/2022-07-446-1/2022|EPI ISL 18901678||A / H5N5
EPI3041341|NS|A/White-tailed eagle/Norway/2022-07-977-1/2022|EPI ISL 18901686||A / H5N5
EPI2972954|NS|A/Great black-backed gull/Norway/2022-07-3018-2/2022|EPI ISL 18851409||A / H5N5
86
EPI3041195|NS|A/Great black-backed gull/Norway/2022-07-2739-1/2022|EPI ISL 18901649||A / H5N5
EPI3041135|NS|A/White-tailed eagle/Norway/2022-07-198-1/2022|EPI ISL 18901641||A / H5N5
EPI3008871|NS|A/Northern gannet/Norway/2022-07-1233-1/2022|EPI ISL 18877920||A / H5N5
65
EPI3413805|NS|A/Red fox/Norway/2024-80-12-1/2024|EPI ISL 19232150||A / H5N5
EPI3413813|NS|A/Red fox/Norway/2024-80-16-1t/2024|EPI ISL 19232151||A / H5N5
EPI2966938|NS|A/White-tailed eagle/Norway/2024-07-14/2024|EPI ISL 18838519||A / H5N5
EPI2815759|NS|A/Gannet/Scotland/124903/2023|EPI ISL 18592416||A / H5N5
EPI2815871|NS|A/Common buzzard/England/264162/2023|EPI ISL 18592430||A / H5N5
63
EPI2815751|NS|A/Great Black-Backed Gull/Scotland/123498/2023|EPI ISL 18592415||A / H5N5
EPI2773666|NS|A/Herring Gull/England/123277/2023|EPI ISL 18378733||A / H5N5
EPI2815887|NS|A/Herring gull/England/404234/2023|EPI ISL 18592432||A / H5N5
EPI2966668|NS|A/large-billed crow/Hokkaido/0112F066T/2023|EPI ISL 18837770||A / H5N5
63
EPI3293018|NS|A/large-billed crow/Hokkaido/0103B118/2024|EPI ISL 19139304||A / H5N5
EPI2962450|NS|A/crow/Hokkaido/B075/2024|EPI ISL 18830860||A / H5N5
87
EPI3158620|NS|A/European Herring Gull/Germany-SH/2024AI01287/2024|EPI ISL 19014082||A / H5N5
64
EPI3205196|NS|A/Northern Fulmar/Netherlands/1/2024|EPI ISL 19053252||A / H5N5
EPI3162760|NS|A/Sparrowhawk/England/133485/2024|EPI ISL 19016255||A / H5N5
EPI3162776|NS|A/Common Buzzard/England/133477/2024|EPI ISL 19016257||A / H5N5
EPI2790999|NS|A/Common eider/Iceland/2023AI08561/2023|EPI ISL 18497662||A / H5N5
EPI2794487|NS|A/European herring gull/Norway/2023-07-3048-1/2023|EPI ISL 18513245||A / H5N5
EPI3097935|NS|A/White-tailed eagle/Norway/2023-07-2937-1/2023|EPI ISL 18961445||A / H5N5
EPI3210810|NS|A/large-billed crow/Iwate/0302G004/2024|EPI ISL 19055417||A / H5N5
EPI3108390|NS|A/large-billed crow/Hokkaido/B090/2024|EPI ISL 18968547||A / H5N5
EPI3416021|NS|A/Gull/Norway/2023-07-2935-1/2023|EPI ISL 19234791||A / H5N5
EPI3041336|NS|A/White-tailed eagle/Norway/2022-07-694-1/2022|EPI ISL 18901685||A / H5N5
EPI2987974|NS|A/White-tailed eagle/Norway/2022-07-100/2022|EPI ISL 18860084||A / H5N5
EPI3041385|NS|A/White-tailed eagle/Norway/2022-07-1141-5/2022|EPI ISL 18901717||A / H5N5
ref|28|A/duck/Bangladesh/17D747/2016 H3N5 2016-12-18
0.0020

### Appendix Tables

**Appendix Table 1.** Summary of clinical observations, gross pathology, and histopathological findings in six red foxes (*Vulpes vulpes*) infected with highly pathogenic avian influenza viruses H5N1 and H5N5 in Norway, 2022–2024.

| Fox (Month, Year) | General information | HPAIV Subtype | Observations | Gross pathology | Histopathology |
|---|---|---|---|---|---|
| Fox Selje (July 2022) | Juv. male; Low BCS; 2.9 kg | H5N1 | Circling | Moderate cadaverosis; pulmonary edema, congestion, multifocal hemorrhages; empty stomach. Fractured skull (euthanized). | Multifocal necrotizing encephalitis; fibrinonecrotizing pneumonia. |
| Fox Refvik (July 2022) | Juv. male; Moderate-low BCS; 3.9 kg | H5N1 | Circling | Moderate cadaverosis; pulmonary edema, congestion and multifocal hemorrhages; empty stomach. | Multifocal necrotizing encephalitis; pneumonia. |
| Fox Averøy (August 2022) | Juv. male; Low BCS; 3.1 kg | H5N1 | Abnormal behavior, approached a house, and died. | Marked cadaverosis; cachectic; pulmonary edema, congestion and multifocal hemorrhages; empty stomach. | Multifocal necrotizing encephalitis; vasculitis; pneumonia. |
| Fox Tromsø (June 2023) | Juv. male; Moderate-low BCS; 2.2 kg | H5N1 | Found dead under a house. | Mild cadaverosis; pulmonary edema and congestion; rodent remains in stomach. | Diffuse pneumonia; no brain lesions. |
| Fox Skibotn 1 (February 2024) | Adult female; moderate-low BCS; 4.3 kg | H5N5 | Circling, forelimb crawling. | Mild cadaverosis; lungs not assessable (gunshot trauma); empty stomach. | Mild lymphohistiocytic encephalitis. |
| Fox Skibotn 2 (February 2024) | Adult female; moderate BCS; 5.2 kg | H5N5 | Circling and hind limb lameness. | Mild-moderate cadaverosis; few, scattered pulmonary hemorrhages; empty stomach, fractured skull (gunshot trauma). | Multifocal necrotizing encephalitis; minimal pulmonary lesions. |

**Appendix Table 2.** Presence of mutations that may indicate mammalian adaptation in highly pathogenic avian influenza viruses H5N1 and H5N5 clade 2.3.4.4b from red foxes (red) and wild birds in Norway 2022-2024.

| **Virus name, GISAID acc. No.** | **Subtype** | **Genotype** | **HA** | **PB2** |
|---|---|---|---|---|
| A/Great black-backed gull/Norway/2022-07-2560-1/2022<br>EPI_ISL_18901648 | H5N1 | EA-2020-C | S133A; T156A;(K218Q+S223R) | |
| A/Gull/Norway/2022-07-3180-1/2022<br>EPI_ISL_18901673 | H5N1 | EA-2020-C | S133A; T156A;(K218Q+S223R) | |
| A/Red fox/Norway/2022-07-2244-1/2022<br>EPI_ISL_18901720 | H5N1 | EA-2020-C | S133A; T156A;(K218Q+S223R) | E627K |
| A/Great black-backed gull/Norway/2022-07-1594-1/2022<br>EPI_ISL_18901638 | H5N1 | EA-2020-C | S133A; T156A;(K218Q+S223R) | |
| A/Red fox/Norway/2022-04-22642-1-4/2022<br>EPI_ISL_19232148 | H5N1 | EA-2020-C | S133A; T156A;(K218Q+S223R) | |
| A/Northern gannet/Norway/2022-07-2085-1/2022<br>EPI_ISL_18901645 | H5N1 | EA-2020-C | S133A;S155N;T156A;(K218Q+S223R) | |

| | | | |
|---|---|---|---|
| A/Northern gannet/Norway/2022-07-1991-1/2022<br><br>EPI_ISL_18901642 | H5N1 | EA-2020-C | S133A;T156A;(K218Q+S223R) |
| A/Red fox/Norway/2022-04-22642-2-4/2022<br><br>EPI_ISL_19232149 | H5N1 | EA-2020-C | S133A;T156A;(K218Q+S223R) |
| A/Herring_gull/Norway/2023-07-1237-t/2023<br><br>EPI_ISL_18939109 | H5N1 | EA-2022-BB | S133A;T156A;(K218Q+S223R) |
| A/Black-legged_kittiwake/Norway/2023-07-1640-1tr/2023<br><br>EPI_ISL_18939115 | H5N1 | EA-2022-BB | S133A;T156A;(K218Q+S223R) |
| A/Red_fox/Norway/2023-80-49-2/H5N1/2023<br><br>EPI_ISL_19232456 | H5N1 | EA-2022-BB | S133A;T156A;(K218Q+S223R) |
| A/ Black-legged_kitiwake/Norway/2023-04-21371-2/2023<br><br>EPI_ISL_18939098 | H5N1 | EA-2022-BB | S133A;T156A;(K218Q+S223R) |
| A/Black-legged_kitiwake/Norway/2023-07-1332-1-s/2023<br><br>EPI_ISL_18939110 | H5N1 | EA-2022-BB | S133A;T156A;(K218Q+S223R) |
| A/Red_fox/Norway/2024-80-16-1t/2024<br><br>EPI_ISL_19232151 | H5N5 | EA-2021-I | S133A;(K218Q+S223R) |

| | | | | |
|---|---|---|---|---|
| A/Red_Fox/Norway/2024-80-12-1/2024<br><br>EPI_ISL_19232150 | H5N5 | EA-2021-I | S133A;(K218Q+S223R) | |
| A/Gull/Norway/2023-07-2935-1/2024<br><br>EPI_ISL_19234791 | H5N5 | EA-2021-I | S133A;(K218Q+S223R) | |
| A/White-tailed_eagle/Norway/2024-07-14/2024<br><br>EPI_ISL_18838519 | H5N5 | EA-2021-I | S133A;S223R | E627K |
| A/White-tailed_eagle/Norway/2023-07-2937-1/2023<br><br>EPI_ISL_18961445 | H5N5 | EA-2021-I | S133A;(K218Q+S223R) | |
| A/European_herring_gull/Norway/2023-07-3048-1/2023<br><br>EPI_ISL_18513245 | H5N5 | EA-2021-I | S133A;(K218Q+S223R) | |
