## Supplementary File 2 for "Highly Pathogenic Avian Influenza Viruses H5N1 and H5N5 in Red Foxes (*Vulpes vulpes*) in Norway during 2022-2024"

### Supplementary Appendix

All genome sequences and associated metadata supporting the findings of this study can be accessed through the persistent digital object identifier https://doi.org/10.55876/gis8.260923pq

In addition to the minted DOI, GISAID also communicates the aggregation of GISAID accession numbers (EPI_ISL_IDs) through the corresponding EPI_SET_260923pq identifier to facilitate both, the acknowledgment of all data contributors and the direct retrieval of the underlying data from GISAID used in this study.

#### Influenza Virus Data Summary

| GISAID Identifier | Digital Object Identifier | Number of individual viruses | Data Collection range | Number of countries/territories |
| --- | --- | --- | --- | --- |
| EPI_SET_260923pq | https://doi.org/10.55876/gis8.260923pq | 380 | 2016-12-18 to 2024-04-30 | 26 |
